# Host-use Drives Convergent Evolution in Clownfish and Disentangles the Mystery of an Iconic Adaptive Radiation

**DOI:** 10.1101/2024.07.08.602550

**Authors:** T. Gaboriau, A. Marcionetti, A. Garcia Jimenez, S. Schmid, L. M. Fitzgerald, B. Micheli, B. Titus, N. Salamin

## Abstract

Clownfishes (Amphiprioninae) are a fascinating example of marine radiation. From a central Pacific ancestor, they quickly colonized the coral reefs of the Indo-Pacific and diversified independently on each side of the Indo-Australian Archipelago. Their association with venomous sea anemones is often thought to be the key innovation that enabled the clownfish radiation. However, this intuition has little empirical or theoretical support given our current knowledge of the clade. To date, no ecological variable has been identified that can explain clownfish niche partitioning, phenotypic evolution, species co-occurrence, and thus, the adaptive radiation of the group. Our synthetic work solves this long-standing mystery by testing the influence of sea anemone host use on phenotypic divergence. We provide the first major revision to the known clownfish-sea anemone host associations in over 30 years, accounting for host associations in a biologically relevant way. We gathered whole-genome data for all 28 clownfish species and reconstructed a fully supported species tree for the Amphiprioninae. Integrating this new data into comparative phylogenomic approaches, we demonstrate for the first time, that the host sea anemones are the drivers of convergent evolution in clownfish color pattern and morphology. During the adaptive radiation of this group, clownfishes in different regions that associate with the same hosts have evolved the same phenotypes. Comparative genomics also reveals several genes under convergent positive selection linked to host specialisation events. Our results identify the sea anemone host as the key ecological variable that disentangles the entire adaptive radiation. As one of the most recognizable animals on the planet and an emerging model organism in the biological sciences, our findings bear on the interpretation of dozens of prior studies on clownfishes and will radically reshape research agendas for these iconic organisms.

## Introduction

**B**roadly distributed on tropical coral reefs throughout the Indian and Pacific Oceans [1], clownfishes (Amphiprioninae) are a charismatic group from the Pomacentridae family. They are comprised of 28 recognized species, which famously form obligate mutualisms with venomous sea anemones (Anthozoa: Actiniaria) and are among the most recognizable animals on the planet. Their small size, hierarchical social structure, and amenability to aquaculture have made them tractable model systems for understanding a wide range of fundamental biological processes [2]. Yet the diversification of clownfishes remains an evolutionary mystery [3]. The group reflects many of the telltale characteristics of classic adaptive radiation [4–7]. Clownfishes have diversified more rapidly than the rest of the Pomacentridae, both in terms of speciation rates [8] and phenotypic change [9], and the mutualism with sea anemones could be considered the key innovation that provided novel ecological opportunity and triggered the radiation of the clade [9]. Despite its apparent conformation to adaptive radiation theory, the enigma of the clownfish radiation is two-fold.

First, unlike most empirical examples of adaptive radiations [10–12], divergent phenotypes in clownfishes are not associated with well-defined adaptive zones (*sensu* Simpson [4]). The question of why clownfishes are phenotypically diverse compared to other Pomacentridae despite occupying the same trophic position and showing similar behaviors remains therefore unanswered. Second, there is no strong evidence of ecological speciation in clownfishes, and most recently diverged species are distributed allopatrically (Fig. S9, S10, S11, S12), which suggests that a standard geographical mode of speciation [13] could have driven clownfish diversification. Hence, finding the hidden determinants of clownfish diversification will significantly increase our understanding of the mechanisms driving evolutionary radiations and will add new light to interpret the evolution of the intensively studied clownfish symbiosis.

Divergent sea anemone host use is the most immediately obvious ecological variable that could potentially explain clownfish evolution [9, 14]. Clownfishes are the only fish species that form obligate mutualisms with sea anemones for their entire lives, which makes their habitat fragmented and population sizes greatly limited by host availability [1]. They establish associations with 10 nominal species of sea anemones [15, 16]. The number of sea anemone hosts each clownfish species associates with is highly variable, with most clownfishes having been observed with multiple host species. Despite the apparent flexibility in host association observed in most species, available host habitat is often saturated, limiting recruitment and reproductive opportunities. Once recruited, migration from one host anemone to another is risky and thus it is likely that most clownfishes stay in the same host their entire lives [2]. Clownfish reproduction is also strictly linked to their host. Only the two largest inhabitants of a host sea anemone are reproductive adults, which form a monogomous relationship until one of them dies. Clownfishes lay and brood eggs on hard substrate directly next to or under the host anemone, and there is evidence that host imprinting may drive clownfish larvae to preferentially settle on the same species of host [17]. This mutualism thus holds the potential to impact diversification in multiple ways. On one hand, host specialization may provide a competitive advantage by monopolizing host resources more effectively in a habitat that is saturated and highly competitive. Conversely, host switching or host generalization could potentially open new unconstrained adaptive zones with reduced competition and predation pressure, theoretically leading to increased diversification through ecological release [4, 7]. In such a competitive environment, strategies circumventing the hierarchical reproductive structure of clownfish might be also advantageous [18, 19]. This would include the foundation of associations with new hosts, which is potentially inducing reproductive isolation and generating incipient species in sympatry. Yet no pattern of host-driven diversification has provided broad explanatory power for explaining the clownfish radiation.

**Table 1.**
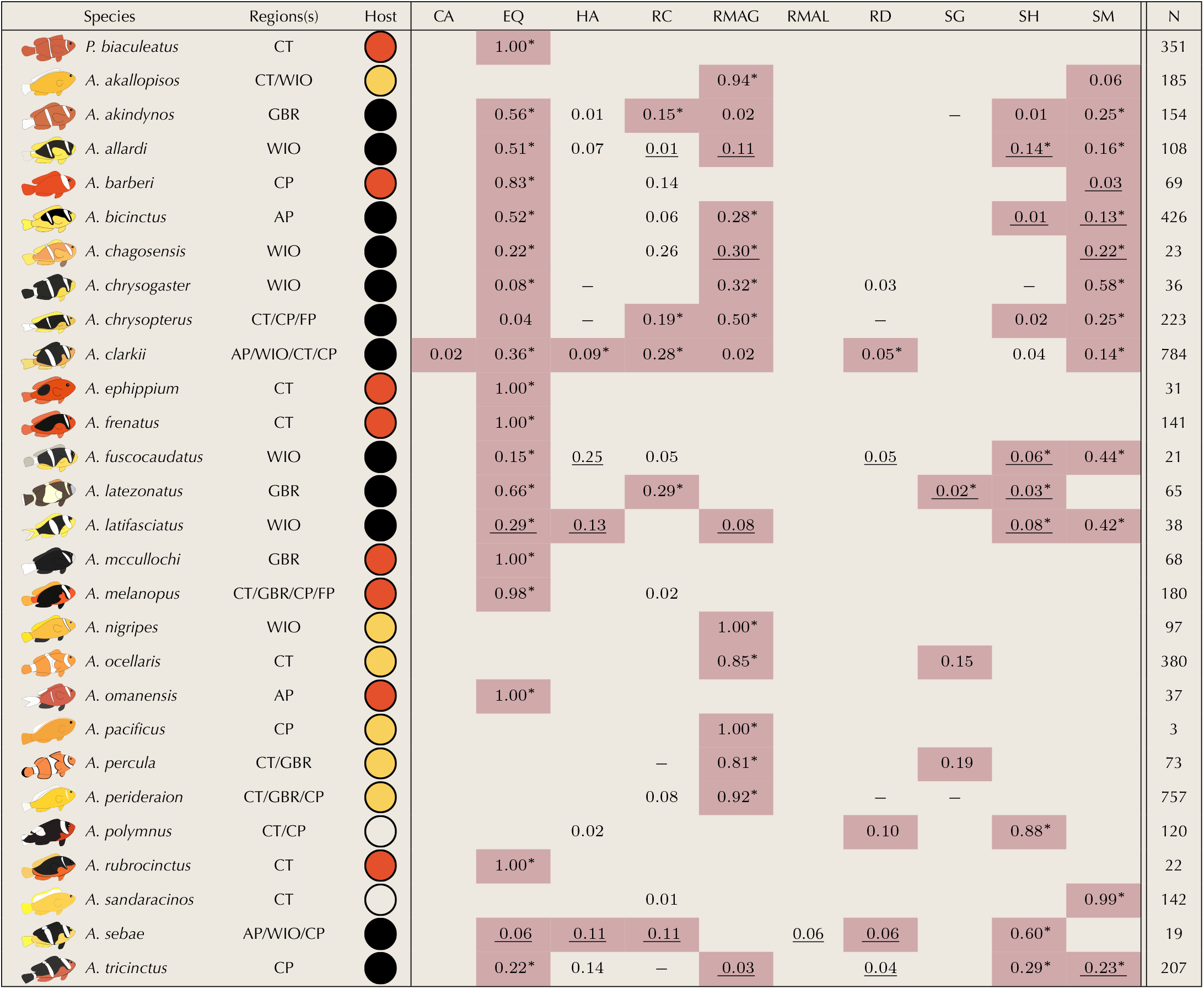
Redefinition of host-use associations based on reproductive habitat. The third column represents the region of occurrence of clownfish species (CT: Coral Triangle, WIO: Western Indian Ocean, GBR: Great Barrier Reef, CP: Central Pacific, AP: Arabian Peninsula, FP: French Polynesia.). The fourth column represents the reproductive host classification that we retained in the following analyses (red: *Entacmaea* specialist, yellow: *Radianthus magnifica* specialist, white: *Stichodactyla* specialist, black: generalist. Columns 5 to 15 represent the proportions of occurrences by species in each species of anemone. Associations highlighted in gray are hosts that serve as reproductive adult habitat. Associations with an * represent primary reproductive host association. Associations that are underlined represent newly documented host associations. Associations with an – are listed by Fautin & Allen 1992 but could not be independently verified (CA: *Cryptodendrum adhaesivum*, EQ: *Entacmaea quadricolor*, HA: *Heteractis aurora*, RC: *Radianthus crispa*, RMAG: *Radianthus magnifica*, RMAL: *Radianthus malu*, RD: *Radianthus doreensis*, SG: *Stichodactyla gigantea*, SH: *Stichodactyla haddoni*, SM: *Stichodactyla mertensii*). The last column represents the number of evaluated associations. Data sources used to quantify host frequency: iNaturalist, field collected data by authors of this study, Elliott & Mariscal (2002) Mar. Biol. 138:23-36, den Hartog 1997, Zool. Med. Leidin 71:181-188, Arvedlund et al. (2000) Environ. Biol. Fishes 58:203-213, Allen et al. (2008) Aqua 14:105-114, Scott et al., 2016, Mar. Biodiv. 46:327-328, Allen et al. (2010) Aqua 16:129-138.

| Species | Regions(s) | Host | CA | EQ | HA | RC | RMAG | RMAL | RD | SG | SH | SM | N |
| --- | --- | --- | --- | --- | --- | --- | --- | --- | --- | --- | --- | --- | --- |
| *P. biaculeatus* | CT |  |  | 1.00* |  |  |  |  |  |  |  |  | 351 |
| *A. akallopisos* | CT/WIO |  |  |  |  |  | 0.94* |  |  |  |  | 0.06 | 185 |
| *A. akindynos* | GBR |  |  | 0.56* | 0.01 | 0.15* | 0.02 |  |  | — | 0.01 | 0.25* | 154 |
| *A. allardi* | WIO |  |  | 0.51* | 0.07 | 0.01 | 0.11 |  |  |  | 0.14* | 0.16* | 108 |
| *A. barberi* | CP |  |  | 0.83* |  | 0.14 |  |  |  |  |  | 0.03 | 69 |
| *A. bicinctus* | AP |  |  | 0.52* |  | 0.06 | 0.28* |  |  |  | 0.01 | 0.13* | 426 |
| *A. chagosensis* | WIO |  |  | 0.22* |  | 0.26 | 0.30* |  |  |  |  | 0.22* | 23 |
| *A. chrysogaster* | WIO |  |  | 0.08* | — |  | 0.32* |  | 0.03 |  | — | 0.58* | 36 |
| *A. chrysopterus* | CT/CP/FP |  |  | 0.04 | — | 0.19* | 0.50* |  | — |  | 0.02 | 0.25* | 223 |
| *A. clarkii* | AP/WIO/CT/CP |  | 0.02 | 0.36* | 0.09* | 0.28* | 0.02 |  | 0.05* |  | 0.04 | 0.14* | 784 |
| *A. ephippium* | CT |  |  | 1.00* |  |  |  |  |  |  |  |  | 31 |
| *A. frenatus* | CT |  |  | 1.00* |  |  |  |  |  |  |  |  | 141 |
| *A. fuscocaudatus* | WIO |  |  | 0.15* | 0.25 | 0.05 |  |  | 0.05 |  | 0.06* | 0.44* | 21 |
| *A. latezonatus* | GBR |  |  | 0.66* |  | 0.29* |  |  |  | 0.02* | 0.03* |  | 65 |
| *A. latifasciatus* | WIO |  |  | 0.29* | 0.13 |  | 0.08 |  |  |  | 0.08* | 0.42* | 38 |
| *A. mccullochi* | GBR |  |  | 1.00* |  |  |  |  |  |  |  |  | 68 |
| *A. melanopus* | CT/GBR/CP/FP |  |  | 0.98* |  | 0.02 |  |  |  |  |  |  | 180 |
| *A. nigripes* | WIO |  |  |  |  |  | 1.00* |  |  |  |  |  | 97 |
| *A. ocellaris* | CT |  |  |  |  |  | 0.85* |  |  | 0.15 |  |  | 380 |
| *A. omanensis* | AP |  |  | 1.00* |  |  |  |  |  |  |  |  | 37 |
| *A. pacificus* | CP |  |  |  |  |  | 1.00* |  |  |  |  |  | 3 |
| *A. percula* | CT/GBR |  |  |  |  | — | 0.81* |  |  | 0.19 |  |  | 73 |
| *A. perideraion* | CT/GBR/CP |  |  |  |  | 0.08 | 0.92* |  | — | — |  |  | 757 |
| *A. polymnus* | CT/CP |  |  |  | 0.02 |  |  |  | 0.10 |  | 0.88* |  | 120 |
| *A. rubrocinctus* | CT |  |  | 1.00* |  |  |  |  |  |  |  |  | 22 |
| *A. sandaracinos* | CT |  |  |  |  | 0.01 |  |  |  |  |  | 0.99* | 142 |
| *A. sebae* | AP/WIO/CP |  |  | 0.06 | 0.11 | 0.11 |  | 0.06 | 0.06 |  | 0.60* |  | 19 |
| *A. tricinctus* | CP |  |  | 0.22* | 0.14 | — | 0.03 |  | 0.04 |  | 0.29* | 0.23* | 207 |

Major fish clades that have undergone adaptive radiation often show a wide range of color patterns and mating preferences, highlighting the role of these factors in the evolution of new species [20, 21]. Amphiprioninae are no exception to this observation as each clownfish species harbours a distinctive coloration defined by the number and thickness of white bars, the shade of orange, and the presence or absence of black patterns. The diversification of color patterns consequently constitutes another potential variable that could explain the clownfish radiation. As sequential hermaphrodites who are reproductively tied to their host, there is no sexual dimorphism in color pattern or apparent sexual selection for a mate. Other hypotheses that have been proposed to explain clownfish color patterns include host venom toxicity [14], species recognition [22, 23], and social signaling [23], but these have recovered only loose correlations between color patterns, ecology and behavior, and do not have broad explanatory power to explain diversification across the entire clade. Diversifying coloration patterns, which allows interspecific co-existence in the same sea anemone [24], could be an alternative strategy to circumvent the restricted reproductive structure of clownfishes.

We revisit and test the role of clownfish and seaanemone host associations on clownfish phenotypic diversification. We publish the first major revision to the known clownfish-sea anemone host associations in over 30 years and propose a novel ecological classification of clown-fishes based on the biological relevance of their host associations. We further generate whole-genome sequencing data for all 28 clownfish species and produce a fully supported species tree for the Amphiprioninae subfamily. We characterize the coloration and morphological phenotypes for all clownfish species and test the influence of host use and biogeography on phenotypic evolution. Finally, we investigate the presence of protein-coding genes showing patterns of convergent positive selection associated with shifts in host use, and thus potentially playing a role in host specialisation. Our findings transform our understanding of clownfish evolution and identify the host sea anemones as the key ecological variable that can broadly explain the clownfish adaptive radiation.

## Results & Discussion

In previous phylogenetic studies of clownfish evolution, sea anemone host association data have been gathered from the seminal work of Fautin & Allen [25, 26].

These reviews provide host association matrices based on direct field observations for each clownfish and sea anemone species. Our understanding of sea anemone host associations has remained largely unchanged for the past 30 years, and only four additional clownfish-sea anemone associations have been documented and synthesized by the most recent review on the subject [27]. Host association data are then taken from these summaries and mapped onto, or incorporated into, resulting phylogenetic analyses to disentangle whether there is any statistical support, or broader pattern, linking clownfish diversification with host-use [9, 14, 22, 28]. In these analyses, all host anemones a clownfish is known to associate with are treated equally, despite well-documented evidence that some clownfish-sea anemone combinations occur much more frequently than others in nature.

Leveraging newly collected host association data throughout the Indo-Pacific, the published literature, and the citizen science database iNaturalist, we first comprehensively update the clownfish-sea anemone association matrix and record 29 new host association across 10 clownfish species that had not been previously documented by Fautin and Allen [25] (Tab.1). Clownfish species are known to compete for “preferred” anemone hosts [27], some of which serve as reproductive adult habitat [29], leaving the less preferred hosts to exclusively harbor juveniles and non-breeding sub-adults [1, 30].We argue that the frequency with which each clownfish species utilizes sea anemone hosts as adult reproductive habitat, termed “reproductive host associations”, captures the most significant evolutionary aspects of the clownfish-sea anemone mutualism. Juvenile clownfish recruits opportunistically to both preferred and non-preferred habitat [29, 31]. However, there is no species-wide ontogenetic migration from less preferred to preferred host species as body size increases [32]. Consequently, these less preferred hosts are likely developmental dead ends.

Focusing solely on reproductive host associations, our updated results bin the clownfishes into four sea anemone host groups: *Radianthus magnifica* specialists, *Entacmaea quadricolor* specialists, *Stichodactyla* specialists and host generalists. Remarkably, the clownfishes comprising each of the four reproductive association groups have similar color patterns (Fig. 2.c) and morphologies (Fig. 2.b). The *R. magnifica* specialists are broadly defined by a light-orange base body color, reduced white stripe width, higher elongation and stronger peduncle. The *E. quadricolor* specialists are defined by a reddish-orange base body color and reduced white stripe number and width. Morphologically, they present a shorter snout and a deeper body. Finally, the generalists are defined by a black trunk with two or three white bands of varying width, along with a morphology more similar to *Entacmaea* specialists. This seems to confirm the intuition that reproductive host-use drives phenotypic diversification in clownfish [9, 14, 22]. We formally test this hypothesis using phylogenetic comparative methods and genomic approaches.

The species tree reconstructed using 10,720 orthologous genes shows a considerable level of gene tree incongruence, as illustrated by the quartet support values (overall normalized quartet score of 0.76, Fig. S1). Nevertheless, the tree is well supported, with all nodes having local posterior probabilities of 1 (Fig. S1). The main clades of clownfishes are confirmed, but we find that *Premnas biaculeatus* is placed as a basal lineage to all other clownfishes (Fig. 1, Fig. S1, S2) contradicting the most recent phylogenetic works on *Amphiprioninae* [9, 33–36]. To date, this constitutes the most robust placement of *Premnas biaculeatus* considering both the scale and quality of the data we analyzed and the support that we found (Fig. S1, S2).

**Fig. 1.**
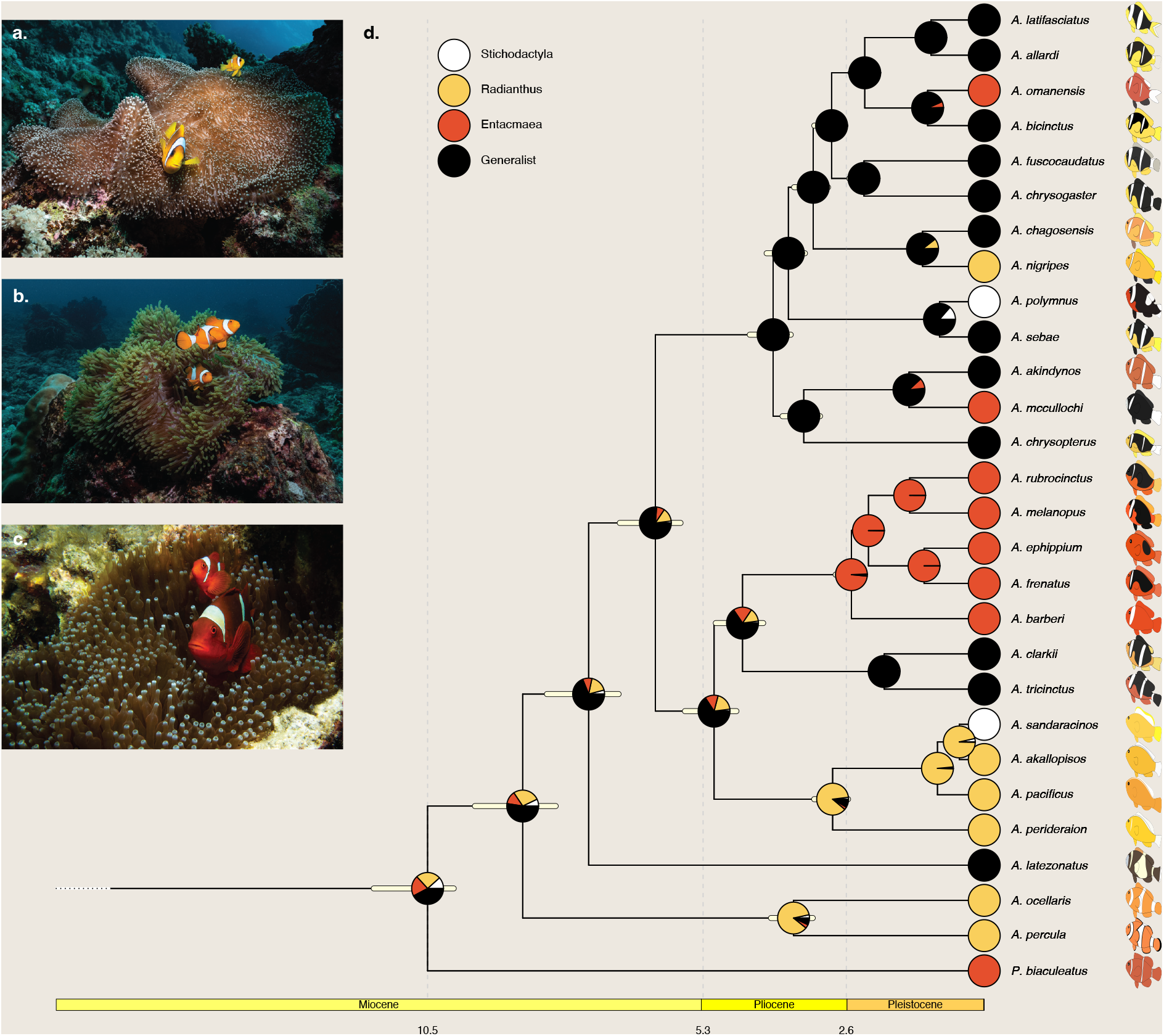
Phylogenetic tree of clownfish and reconstruction of ancestral host associations. **a**.: Picture of *Stichodactyla mertensii*. **b**.: Picture of *Radianthus magnifica*. **c**.: Picture of *Entacmaea quadricolor*. Pictures credit: Lucy Fitzgerald. **d**. Clownfish phylogenetic tree reconstructed from Whole Genome Sequencing Data. Yellow bars represent the highest posterior density of node ages in my. Pies at the tips represent the reproductive host association classification. Pies at the nodes represent the marginal probability of each ancestral reproductive host association.

Ancestral state reconstruction of reproductive host associations coupled with biogeography suggests a stepping-stone pattern of clownfish diversification, starting with the colonization of new regions by generalists followed by subsequent specialisation events (Fig. 1, S13). Our results indicate that the common ancestor of all clownfishes was likely a generalist. Previous studies demonstrated that melanin production and white stripes formation and development appears to only be lost across the phylogenetic tree as specialist lineages evolved from generalist ancestors [37, 38]. This is concordant with the present broad phenotype of generalists characterized by black body color and multiple white stripes. We also show that specialisation to the same host anemone happened multiple times in the clownfish evolutionary history, with four specialisation events to *Entacmaea*, two to *Stichodactyla* and three to *Radianthus*. The most recent specialisation events (*A. nigripes, A. omanensis, A. mccullochi, A. sebae*) happened in marginal areas of clownfish distribution in regions that are not occupied by other specialist species (Fig. S9,S10,S11). Our classification coupled with the most recent evaluation of present clownfish distribution [39] reveals that most species sharing reproductive host association categorisation do not co-occur (Fig. S9,S10,S11,S12). Notable exceptions to this pattern are caused by the basal clades *P. biaculeatus* and *A. ocellaris/A. percula* co-occuring with distantly related specialists, the recent adaptation of *A. sandaracinos* to *Stichodactyla* in sympatry with *A. polymnus* [40], and generalist species on the Great Barrier Reef. This repeated specialisation events following a rapid colonization of the Indo-Pacific from a Indo-Australian Archipelago ancestor created similar assemblages of clownfish phenotypes in each region (Fig. S8).

Clownfish phenotypes, characterized by analyzing color patterns and morphology from standardized photographs (museum specialist collections and photographs from public databases), are found to be convergent within reproductive host groups. Phylogenetic MANOVAs show a significant effect of host association on both color patterns (Fig. 2.c Pillai: *S* = 0.8181, *p* = 0.003) and Procrustes morphology phenotype (Fig. 2.b Pillai: *S* = 1.5280, *p* = 0.001), being independent of phylogenetic effects. Similarly, univariate and multivariate phylogenetic comparative analysis indicate that clownfish phenotypic evolution is driven by the reproductive host association (Fig.S23, S24, S25, S26, S22). Models comparison indicates that several axes of color pattern (PC 1 and 2) and morphology (PC 1 and 3) follow an Ornstein-Ulhenbeck mode of evolution [41] with different evolutionary optima depending on their reproductive host association. These results provide, for the first time, robust empirical support that sea anemone hosts are driving the convergent evolution of clownfish phenotypes. Our simulations (Fig. 3.a-d) illustrate the strength of attraction towards those optima associated with host shifts. We predict that color and morphology can converge rapidly following specialisation to a particular host anemone.

**Fig. 2.**
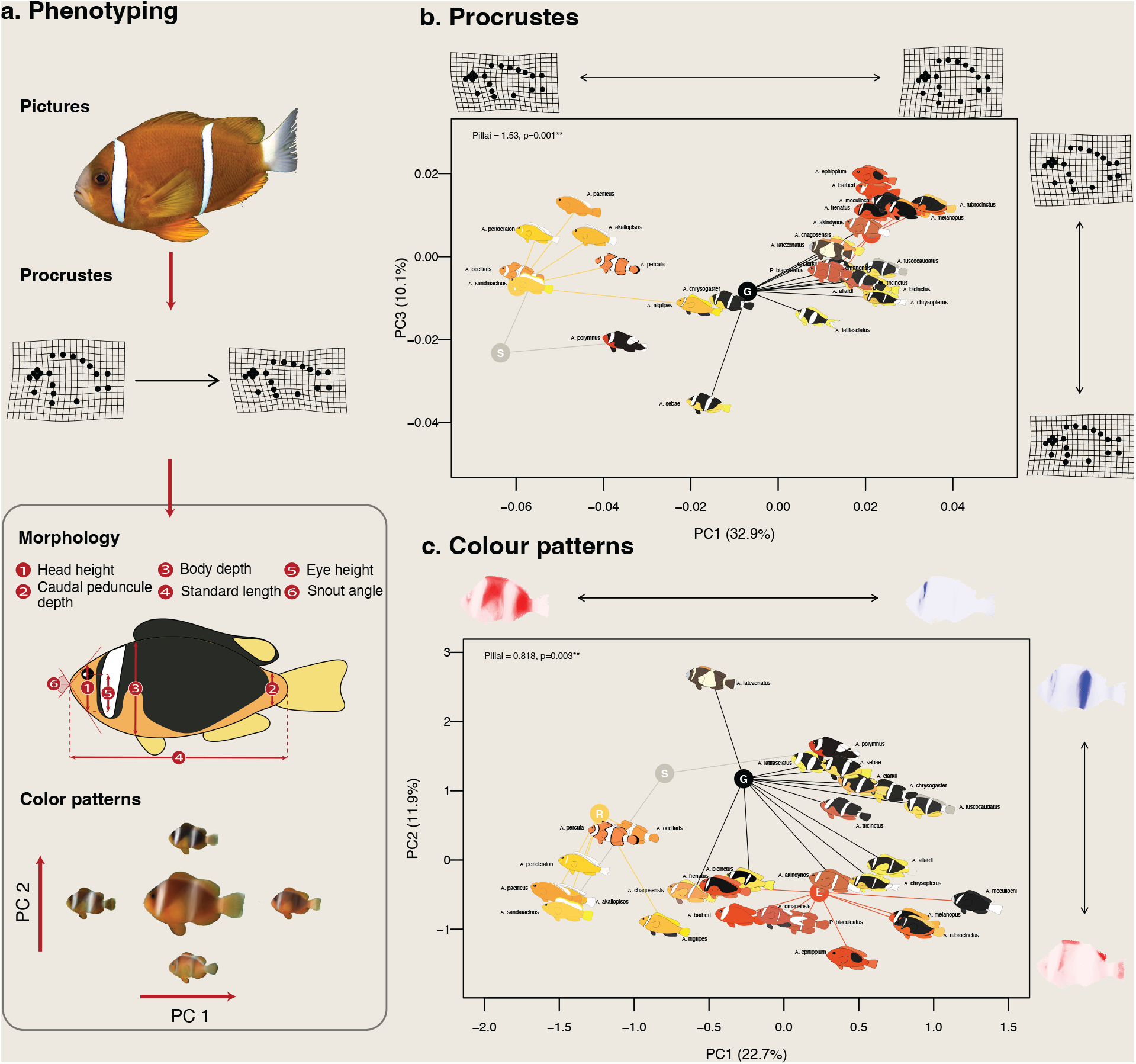
Characterization of clownfish phenotype. **a**.: Workflow for the characterization of clownfish phenotype. For morphological information, pictures of the left side of specimens were obtained from museum specimens. Following Rolland et al. 2018, we collected pictures in museum and specialist collections for several individuals per species (total of 307 individuals, with an average of 9 individuals per species, minimum of 2 and maximum of 31). For color patterns, we collected pictures of the left side of live specimens taken on the field or from iNaturalist (total of 214 individuals, with an average of 8 individuals by species, minimum of 4 and maximum of 10). **b**.: Projection of clownfish species on axes 1 and 3 of the morphospace. Warp grids represent morphological variation along axes. PC1 represents variation in elongation with deep-bodied individuals to the right of PC1 and elongated individuals to the left of PC1. PC3 represents variation in peduncle thickness and jaw, individuals at the top having a thinner peduncle and a lower jaw. **c**.: Projection of clownfish species on axes 1 and 2 of the color pattern space. Heat maps represent color patterns variation along axes. Intensity of red represent the strength of each pixel negative contribution to the axis. Intensity of blue represent the strength of each pixel positive contribution to the axis. PC1 represents variation in off stripe patterns and head white stripe. Individuals to the left of PC1 have bright orange off stripe patterns and no head stripe while individuals to the right of PC1 have dark off stripe patterns and a head stripe. PC2 represents variation in dorsal fin color and second stripe patterns. Colored dots represent the effects of host associations on the variance of clownfish phenotypes estimated using a phylogenetic multivariate ANOVA (R. *Radianthus* specialists, E. *Entacmaea* specialists, S. *Stichodactyla* specialists, G. Generalists).

**Fig. 3.**
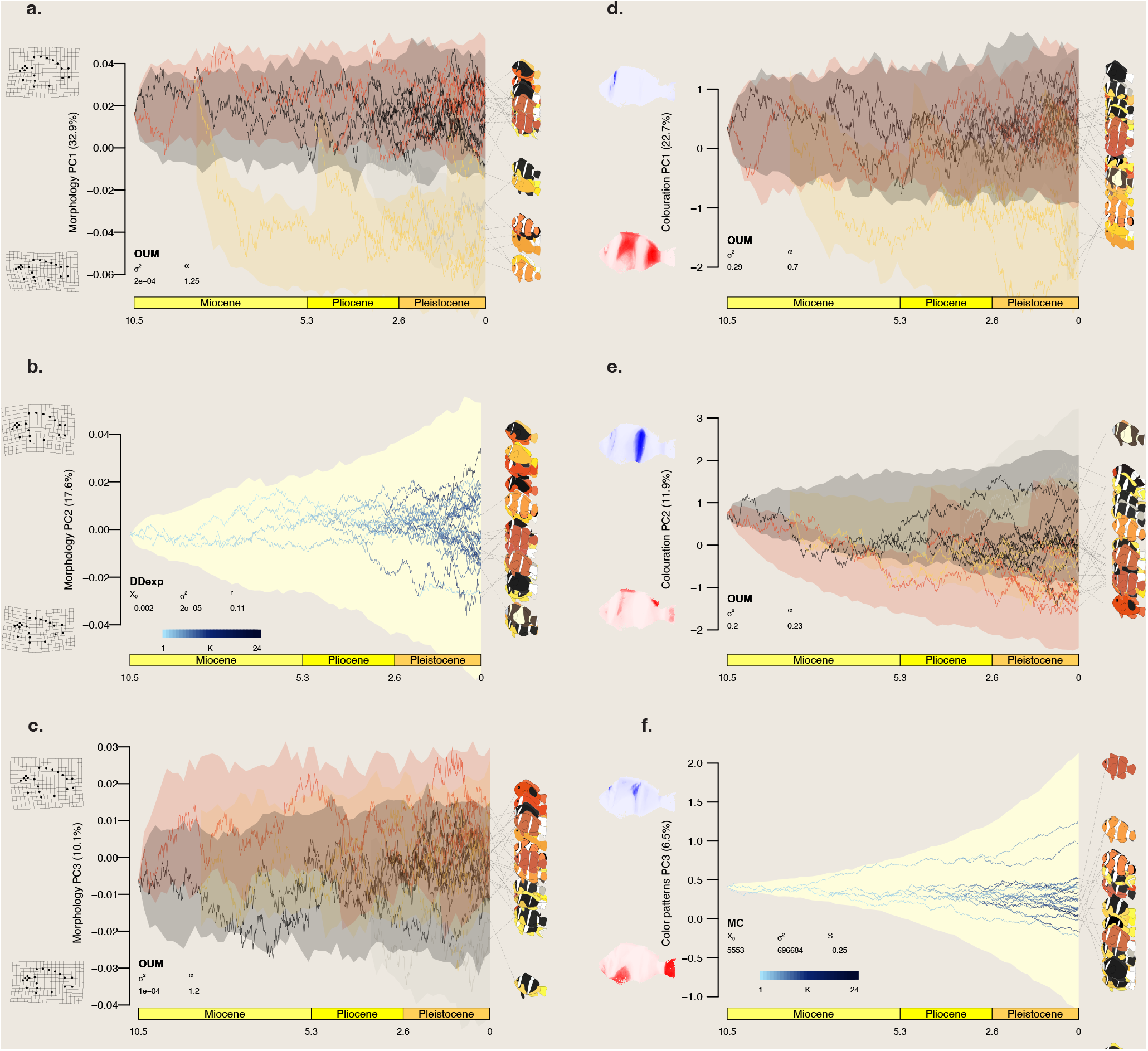
Influence of host-use on clownfish species phenotypes. For each trait, we simulated 100 evolutionary scenarios based on the selected univariate model with corresponding estimated parameters. Lines represent a random selected simulated evolutionary scenario along the phylogenetic tree. Bifurcations correspond to estimated divergence times from the clownfish chronogram. Areas correspond to the range of values taken by our simulations at a given time. In **a**.,**c**.,**d**. and **e**., colors represent the marginal reconstruction of reproductive host associations (Black: generalists; Grey: *Stichodactyla* specialists; Yellow: *Radianthus* specialists; Red: *Entacmea* specialists). In **b**. and **f**., the blue gradient represents the number of lineages co-occurring with the lineage represented by the branch. Graphs in the left column represent evolutionary trajectories of morphology PCA axis, warp grids represent morphological variation along axes. Graphs in the right column represent evolutionary trajectories of RGB color patterns PCA axes, heat maps represent color patterns variation along axes. Intensity of red represent the strength of each pixel negative contribution to the axis. Intensity of blue represent the strength of each pixel positive contribution to the axis. **a**. Morphology PC1 captures the variation in body shape, with deep-bodied species positioned on the top and elongated species on the bottom side of the PC1 axis. **b**. Morphology PC2 discriminates species with a smaller snout angle (small values) to those with a stronger snout angle (large values) .**c**. Morphology PC3 captures the variation in peduncle thickness and lower jaw position,species at the top having a thinner peduncle and a lower jaw. **d**. RGB color patterns PC1 represents variation in off stripe patterns and head white stripe. Species at the bottom of PC1 have bright orange off stripe patterns and no head stripe while species at the top of PC1 have dark off stripe patterns and a head stripe. **e**. RGB color patterns PC2 captures the variation in dorsal fin color and second stripe patterns. Low values are associated with an absence of second stripe and a bright dorsal fin, High values are associated with a white second stripe and a dark dorsal fin. **f**. RGB color patterns PC3 captures the variation in caudal fin brightness (bright species have low values) and stripes shape

Comparative analyses of protein-coding genes reveal that the repeated specialisation to *Radianthus magnifica* is associated with convergent positive selection at the gene level. We identify 8 genes (3 genes when long branches are excluded from the analysis) positively selected along the branches leading to specialisation to *Radianthus magnifica* among which we find the serine protease FAM111A involved in DNA replication with antiviral function in humans [42] and the Crumbs homolog 1 involved in photoreceptor morphogenesis [43]. Similarly the repeated specialisation to *Entacmaea quadricolor* is associated with convergent positive selection in 165 genes (76 genes when long branches are excluded from the analysis). Within them, we found genes coding for the *β*-carotene 15,15’-dioxygenase, the olfactory receptor 11A1, and the retinitispigmentosa 1-like 1 protein), which are associated with olfactory or vision functions. These analyses did not result in the identification of genes evidently linked to coloration and cannot directly explain the phenotypic convergence we observe. However, functional annotation is still lacking for many genes identified as being under convergent positive selection during host specialisation (Tab. S3S5). Furthermore, the number of genes experiencing convergent positive selection associated with host shifts could be underestimated, as shown by the low power of the analyses in the case of weak to moderate strength of selection (Fig. 4). Alternatively, differential levels of expression or changes in regulatory regions could also play role in modulating convergent phenotypes associated with reproductive-host associations.

**Fig. 4.**
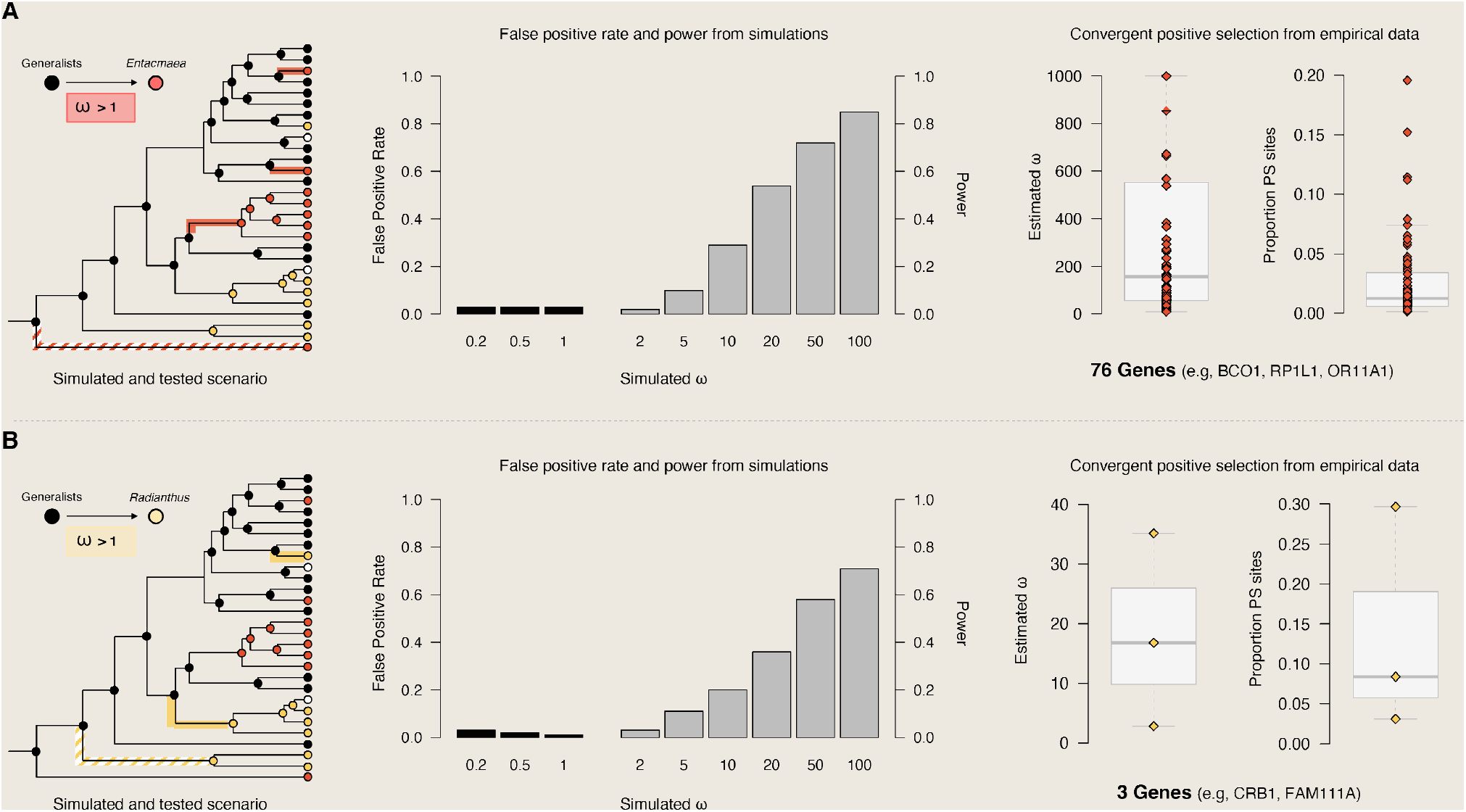
Convergent positive selection associated with shifts in sea anemones hosts during clownfish diversification. Shifts from generalists to *Entacmaea* (A) or *Radianthus* (B) specialists were considered. A, B, left panels) Sequences were simulated under scenarios of presence (*ω >* 1) or absence (*ω* ≤ 1) of convergent positive selection associated with shifts to *Entacmaea* (A) or *Radianthus* (B) sea anemone hosts. Colored branches represent foreground branches (with/without positive selection) for simulations and tests, while the hatched-colored branches were excluded from the tests on empirical data as potentially increasing the false positive rate (Fig. S7). For each *ω* value, a total of 100 sequences (100 replicates) with a length of 1,000 codons were simulated. A, B, middle panels) Tests for positive selection were performed on simulated sequences using the branch-site model implemented in codeml to obtain false positives (simulated *ω ≤* 1, estimated *ω >* 1) and true positives (simulated *ω >* 1, *ω >* 1). False positive rate and power were obtained as the proportion of false positives and true positives, respectively. A, B, right panels) Results of the estimated *ω* and the proportion of positive selected sites for genes under convergent positive selection during shifts to *Entacmaea* (A) and *Radianthus* (B) hosts. Each point corresponds to a significant gene, and the boxplots show the distribution of values of all significant genes. A total of 18,380 clownfish orthologous genes were tested using the branch-site model implemented in codeml, and the total number of significant genes after multiple testing corrections is reported. Genes under convergent positive selection include BCO1 (Q9I993), RP1L1 (Q8IWN7), ORA11A1 (Q9GZK7), CRB1 (P82279) and FAM11A (Q96PZ2).

Our results, however, clearly establish an association between reproductive-host specialisation and both phenotypic and genotypic convergence. Differences between host anemone habitat preferences [29, 44, 45] can partly explain morphological convergence relative to visual acuity and swimming abilities. For instance, *Entacmaea quadricolor* settles within holes and crevices of coral structures, often forming large colonies. Species occupying this anemone species can hide and swim from one individual to another easily. Their high body depth improves manoeuvrability in this environment unlike *Radiathus magnifica* specialists that are more exposed to open environments and currents [44] or *Stichodactyla* specialists that live in exposed flat surfaces. However, the role of color patterns in clownfish life-history seems less straightforward. Recent research suggests that color patterns are an important social signal in clownfish [38, 46]. Clownfish are well equipped to detect UV wavelengths [47] and modulation of UV reflectance through orange and white colors serve as social signaling of dominance within an anemone [48]. Vertical white bars seem to be particularly important in those interactions as clownfish tend to be more aggressive towards similarly patterned individuals to defend their host [23]. Specialization coincides with convergent changes in color and in photo-receptor genes which seems to align with the social function hypothesis. Nevertheless, the interpretation of orange/red/black patterns in an ecological context requires further understanding of the role of coloration in reef ecosystems.

Several phenotypic traits follow a density-dependent evolutionary trajectory based on co-occurrence between clownfishes rather than a convergent regime associated with host anemone. For instance, we found that evolutionary rates of RGB color patterns PC3 -head bar width and dorsal patterns-(Fig. S23, S32) and body-depth (represented by morphological PC2) (Fig. S25, S41) are higher in species co-occurring with several other lineages [49]. This suggests that interspecific interactions increase disparity in blackness and body-depth. Furthermore, the matching competition model [49] was found to be the best model for explaining color pattern PC3 evolution -pectoral and caudal fins patterns and mid-body vertical bar pattern-, meaning that trait tends to drift away from the normative value found in the community. These results highlight the complexity of phenotypic evolution in clownfishes, with certain patterns showing signatures of convergence based on the reproductive host while others exhibiting diverging trends influenced by the community composition and species interactions. Species recognition and hybrid avoidance may explain the high divergence between color patterns. Common instances of host-cohabiting species are characterised by individuals differing in their number of vertical white bars and often consist of generalist-specialist pairs [24, 50]. Taken together, our results reveal highly segregated community assembly in clownfishes at different scales. We first observe that co-occurring species rarely occupy the same reproductive hosts. When they do, species sharing a reproductive host strategy are phylogenetically distant and show a divergent vertical white bar pattern despite being convergent in other patterns (Fig. S8). This is supported by the non-random assembly of vertical white bar patterns in clownfish communities [38]. Although our data does not allow us to test further segregation at different scales, field observations tend to support it. For instance, segregation between species occupying the same anemones was observed along depth gradients in several clownfish communities [51, 52].

## Conclusion

We provide a robust and complete revision of the clownfish phylogeny using whole genome sequencing, a new classification of interactions between clownfish and sea-anemone species, and an in-depth analysis of coloration and morphological trait variability in the clade. This new data allowed us to identify a unique case of phenotypic and genotypic convergence driven by reproductive hosts. Our results suggest that evolutionary transitions from generalist to specialized reproductive host associations affected both coloration patterns, morphology and community assembly processes in clownfishes. It opens many new questions about the biology of clownfishes and adaptive radiations driven by competition.

## Methods

### Genetic samples selection and sequencing

Raw sequencing data for nine species (*Amphiprion akallopisos, A. bicinctus, A. clarkii, A. frenatus, A. melanopus, A. nigripes, A. percula, A. perideraion, A. sebae*) was available from previous studies (PRJNA433458, [53]; PRJNA515163, [35]; PRJNA1022585 [40]; PRJNA1025355[54]). Fin clips for the remaining 19 species were obtained from collaborators between 2013 and 2018. We extracted the genomic DNA with the DNeasy Blood and Tissue Kit (Qiagen, Hilden, Germany) following the manufacturer’s instructions, and we prepared short-insert (350 bp) paired-end libraries using the TruSeq Nano DNA Library Preparation Kit (Illumina, San Diego, California, USA). We validated the fragment length distribution of the libraries with a fragment analyzer (Agilent Technologies, Santa Clara, California, USA). Libraries generated in 2017 (Tab. S1) were pooled by five, and each pool was sequenced on a single lane of Illumina HiSeq2500 (read length: 100 bp) at the Lausanne Genomic Technologies Facility (Lausanne, Switzerland). The remaining libraries were sequenced on Illumina HiSeq4000 at the Genomics Platform of iGE3 (Geneva University, Switzerland), multiplexing eight samples per lane (Tab. S1).

### Reads processing, mapping, and gene sequence reconstruction

We removed adapter contaminations and trimmed the sequencing reads using Trimmomatic (v.0.36; [55]) with the following parameters: ILLUMINACLIP:TruSeq3-PE.fa:2:30:10 |LEADING:3 |TRAILING:3 |SLIDINGWINDOW:4:15 |MINLEN:36. We verified read quality before and after the processing with FastQC (v.0.11.5; [56]).

We mapped the trimmed reads of each species against the set of protein-coding genes of *A. frenatus* (31,054 genes, ca. 83.3 Mbp; https://doi.org/10.5061/dryad.nv1sv; [53]) using BWA (v.0.4.15; [57]). Following the framework reported in DePristo et al. [58], we processed the mapping results with samtools (v.0.1.19; [57]) to keep only primary alignments and mark potential read duplicates. We performed variant calling and filtering with the GATK software (v.3.7; [59]), following GATK best practices [58]. We extracted SNPs (task SelectVariants) and filtered them (task VariantFiltration) using the following parameters: *QD <* 2.0, *FS >* 200, *ReadP osRankSum <* −20.0, *SOR >* 10.

We reconstructed the gene sequences for each species using *A. frenatus* sequences as the reference and integrating the variants information for each species. This approach is legitimate because the divergence of the group is low [35], and in general, protein-coding genes are highly conserved. Nevertheless, to verify the accuracy of the reconstructed genes, we compare sequences obtained here for *A. akallopisos, A. bicinctus, A. frenatus, A. melanopus, A. nigripes, A. sebae* with the sequences previously obtained by independent genome annotation (https://doi.org/10.5281/zenodo.254024, [35]).

We evaluated the approximate position of the genes on the chromosome-level assembly of *A. percula* [60] by performing reciprocal best hits analyses with BLAST [61]. We filtered the results to keep only genes with sequences retrieved for all species, being at least 150 bp long and with a best reciprocal hit on *A. percula* chromosomes, for a total of 18,390 genes.

### Phylogenetic reconstruction and dating

To root the phylogenetic tree, we integrated a damselfish outgroup in the analysis. We retrieved protein-coding gene sequences for one individual of *Pomacentrus moluccensis* (32,027 genes; https://doi.org/10.5281/zenodo.254024, [35]). To obtain putative orthologs, we performed reciprocal best hits analyses with BLAST [61] between *A. frenatus* and *P. moluccensis* genes. From the 18,390 genes, only those with *P. moluccensis* orthologs were considered for phylogenetic reconstruction, corresponding to 10,720 genes.

We aligned the gene sequences of all clownfish species and *P. moluccensis* with MAFFT (strategy L-INS-I; v.7.841; [62]), and we reconstructed the gene trees with RAxML (GTR+G model, 100 bootstrap replicates; v.8.2.12; [63]). We estimated the species tree with ASTRAL-III (v.5.7.8; [64]) from the 10,720 gene trees. Since contracting very low-support branches in gene trees enhances ASTRAL-III accuracy [64], branches with bootstrap support below 10% were assigned a length of 0 using the Newick Utilities ([65]). We plotted ASTRAL results with the R package AstralPlane (v.0.1.1; https://github.com/chutter/AstralPlane.git).

We used SortaDate [66] to select the 20 most informative genes for the estimation of the calibrated phylogenetic tree of clownfishes. The selection considered the genes with the lowest topological conflict with the species tree and with the lowest variance in the root-to-tip distance [66]. We inferred the calibrated phylogenetic tree with BEAST2 (v.2.6.3; [67]). For each partition, we applied a GTR+G model of substitution and an uncorrelated relaxed clock with a lognormal distribution. Since no fossil records were available for clownfishes, we used a secondary calibration point, setting a uniform prior from 10 to 18 MYA for the crown age of clownfishes [3]. To facilitate the convergence of the MCMC, we constrained the seven major clades (as defined in [3]) to be monophyletic, in accord with the species tree obtained with ASTRAL-III (Indian clade: *A. omanensis, A. bicinctus, A. allardi, A. latifasciatus, A. nigripes, A. chagosensis, A chrysogaster, A. fuscocaudatus*; Polymnus clade : *A. polymnus, A. sebae*; Ephippium clade: *A. melanopus, A. ephippium, A. rubrocinctus, A. frenatus, A. barberi*; Australian clade: *A. mccullochi, A. akindynos*; Clarkii clade: *A. clarkii, A. tricinctus*; Akallopisos clade: *A. akallopisos, A. perideraion, A. sandaracinos*; *A. pacificus*; Percula clade: *A. ocellaris, A. percula*). We applied a birth-death model, and we ran four independent MCMC chains, each with 50 million generations and sampling every 1,000 generations. For each chain, we examined the results with TRACER (v.1.7.2; [68]) and convergence was assessed by effective sample sizes (ESS) above 200. The posterior distributions were combined using LogCombiner (BEAST2, [67]), and a maximum clade credibility tree was summarized with TreeAnnotator (burn-in percentage = 20%, posterior probability limit = 0.5; BEAST2, [67]). Plotting was performed with FigTree (v.1.4.4; https://github.com/rambaut/figtree.git).

### Image data collection and processing

We collected clownfish pictures from different public image databases (for instance, Reef Life Survey, Fishbase, Fishes of Australia; details for all images in Appendix). For color pattern analyses, images were selected when the fish were shown in a lateral view with sufficient illumination and quality to reveal the entire body. From each of the species inventory of images collected, we selected a minimum of four images with the highest quality and resolution. On those images, we standardized size and resolution, corrected the white balance, and homogenized the background with a neutral color contrasting with the targeted colors used in our analyses (RGB background color code was 0,165,255). These editions were carried out using Adobe Photoshop CC 2019 [69]. A total of 214 images accounting for the 28 different clownfish species met our criteria to proceed with further color analyses. For morphology analyses, we complemented this dataset with museum and private collections specimen to reach a total of 312 standardised left side pictures. We removed pictures background and set 29 morphological landmarks (landmarks shown in Fig S14) were set on each image using FIJI software [70]. Images were transformed with the R package *geomorph* [71] using Procrustes superimposition to a ‘mean shape’ ensuring all images were aligned when performing coloration pattern and morphology analyses.

### Coloration patterns

We estimated color pattern variation among individuals with four independent datasets. Clownfish patterns are constituted by three dominant colors: black (RGB color code: 0,0,0), orange (RGB color code: 255,165,0) and white (RGB color code: 255,255,255). We used the R package Patternize [72] to extract layers of patterns of the three colors from transformed images using *patLanRGB* function. This generated three datasets containing, respectively, black, white and orange patterns. The fourth dataset was built by extracting RGB channels from each image and concatenating the values from the three channels. This dataset represent the total color variation among clownfish individuals. For each, dataset we performed a PCA analysis using the *patPCA* function to capture color pattern variation.

### Morphology

We generated two morphology datasets. We used the 29 landmarks positions after Procrustes superimposition to generate a PCA of individual morphological variation. This allowed us to capture the total variation in individuals morphology. We also estimated five traits of interest for fish relating to swimming abilities, position in the water column or visual acuity (body ratio, peduncle ratio, eye-height ratio, head ratio, snout angle, Fig. 2) [73, 74].

### Ancestral state reconstructions

We inferred co-occurrence between lineages in space and time using our newly reconstructed phylogenetic tree and species occurrences obtained from the GBIF and INaturalist databases. Occurrences were thoroughly checked and filtered and ranges were assessed using species distribution modelling from [39]. We performed ancestral state reconstruction of clownfish biogeography using maximum likelihood estimation of DEC, DEC+J, DIVA, DIVA+J and BAYAREA, BAYAREA+J models parameters using the R package BioGeoBEARS [75]. We ran the model with 6 bio-regions identified based on coral reef fish communities [76] and calculated adjacencies and dispersal distances in meters between regions. We selected the best fitting model using AIC comparisons (Tab. S6). We then performed Biogeographic Stochastic Mapping using the best model (DIVA) S13 and generated 100 stochastic maps of co-occurrence between lineages throughout the phylogenetic tree.

We also reconstructed ancestral states of adult host occupancy based on the four categories identified as carrying the same biological characteristics: *Entacmaea* specialists, *Radianthus* specialists, *Stichodactyla* specialists and generalists. We used the R package *corHMM* [77] to test for five different transition matrices between the four states, with and without hidden states: (1) all rates different, (2) symmetric rates between reproductive association categories, (3) all rates equal, (4) equal rates between specialists but different rates for specialists to generalists and generalists to specialist, and (5) equal rates between specialists and equal rates between generalist and specialists. We compared AIC [78] fit between models and retained the best model (model 5 without hidden states) to estimate marginal probabilities of states at each node (Fig. 1d). We generated a 100 stochastic maps of ancestral character reconstruction using the R package *phytools* [79] with the same transition matrix.

### Simulations for positive selection analyses

To investigate the false positive rate (FPR) and sensitivity (power) to detect convergent positive selection on protein coding genes associated with host shifts, we simulated data with the program evolver (PAML, v.4.9; [80]; Fig. S3). We simulated alignments of 1,000 codons under the branch-site model, following the clownfish species tree (total of 28 tips) with the identified host shifts (4 to *Entacmaea* specialists, 3 to *Radianthus* specialists). We simulated 100 alignments with no (*ω* of 0.2, 0.5, and 1.0) or an increasing strength (*ω* of 2, 5, 10, 20, 50, and 100) of positive selection on the branches with host shifts (*Entacmaea* or *Radianthus*, Fig. S3A). We then tested for positive selection on the simulated sequences using the branch-site model implemented in codeml (PAML, v.4.9; [80]). We set the branches with the host shifts (*Entacmaea* or *Radianthus*) as foreground branches (Fig. S3B). We fitted a null model (H0), where the foreground *ω* is constrained to be smaller or equal to 1, and we compared it with the alternative model (H1), where the foreground *ω* is estimated but forced to be larger than 1. The best model was determined with a likelihood ratio test (LRT; df=1). We obtained the FPR and sensitivity (or power) by calculating the proportion of significant LRT (p-value < 0.05) for the alignments simulated without or with positive selection, respectively (Fig. S3).

We examined the sensitivity to detect postive selection associated with a single shift to *Entacmaea* or *Radianthus* (i.e., selection on a single branch, without considering the other branches with identical shifts, Fig. S3B). We used the alignment simulated with increasing strength (*ω* of 2, 5, 10, 20, 50, and 100) of positive selection on the branches with host shifts (*Entacmaea* or *Radianthus*, Fig. S3). We considered each shift separately and purged the additional equivalent shifts from the tree and the alignments (dotted lines in Fig S3B). We tested for positive selection and calculated the sensitivity of the analyses as described above, but by setting a single branch as the foreground. We sequentially tested each branch with a shift to *Entacmaea* or *Radianthus*.

False positives in convergent analyses could also arise from positive selection occurring on a single branch and driving the significance of the LRT. We tested this by simulating alignments with increasing strength of positive selection as described above, but with the positive selection only occurring in the longest branch with a shift (“Long branch” scenario, Fig. S3A) or in the branch where the shift is concerning an entire clade (“Clade branch” scenario, Fig. S3A). We then tested for positive selection on all branches with host shifts (*Entacmaea* or *Radianthus*, Fig. S3B) and calculated FPR (Fig S3C-D).

### Positive selection analyses on the real data

We verified that the alignments of the 18,390 protein-coding genes corresponded to codon alignments using the annotated protein sequences of *A. frenatus* (https://doi.org/10.5061/dryad.nv1sv;[53]) and the software Pal2Nal [81]. We then tested for positive selection associated with host shifts to *Entacmaea* or *Radianthus* sea anemones (Fig. 4) using the branchsite model in codeml (PAML, v.4.9; [80]). We set the shifts of hosts as foreground branches (4 shifts for *Entacmaea*, 3 shifts for *Radianthus*, Fig. 4). As for the simulated datasets, we compared for each gene the null model (with the foreground *ω* <=1) to the alternative model (with fore-ground *ω >* 1) with a likelihood ratio test (df=1). We corrected p-values for multiple testing using the Benjamin-Hochberg method implemented in the *qvalue* R package (FDR level of 0.05, method bootstrap, option “robust”; [82]).

Because simulations highlighted potential false positives arising from the presence of long branches in the tests, we retrieved the significant genes showing convergent positive selection associated with host shifts and tested again for positive selection without including the longest branches (Fig. S5). For each gene, the best model was chosen with a LRT (df=1), and significant results were categorized as “genes under convergent positive selection”. Genes that were not significant may be false positives due to the presence of the long branch or true positive with increased power of the analyses given by the long branch, and they were thus labeled as “genes potentially under convergent positive selection”.

For the genes showing significant positive selection associated with shifts to *Entacmaea* host, we performed gene ontology (GO) enrichment analyses using the *topGO* package (parameters: Fisher’s exact tests, weight01 algorithms, minimum node size of 5; v.2.26.0; [83]). We contrasted the annotations of the significant genes against those of the analyzed 18,390 protein-coding genes. Results were considered significant at p-values < 0.01. No corrections for multiple tests were performed, following recommendations from the topGO manual. The same analyses was not done for shits associated with *Radianthus* host because of the few number of significant genes obtained (see Results and Discussion).

### Phylogenetic comparative analysis

We performed Maximum Likelihood estimation of color and morphology evolutionary rates using alternative models. We evaluated the five color patterns and morphology datasets independently. First, we used a multivariate approach, taking all dimensions into account at the same time with simple models using the R package *mvMORPH* [84]. This approach aimed at testing whether color patterns and morphology are phylogenetically conserved among clades, (Brownian Motion (BM) = phylogenetic signal vs White Noise (WN) = no phylogenetic signal), whether they have an optimal value (Ornstein-Uhlenbeck (OU) = optimal color and/or morphology vs Brownian Motion = random evolution of coloration and/or morphology). Second, we used an univariate approach, independent for each dimension of the color and morphology analysis, using more complex models of evolution taking density effects or the effect of the host into account. We compared univariate OU, BM and WN models fitted with the R package *geiger* [85] to an OU model with several optima determined by the host anemone (OUM) fitted with the R package *OUwie* [86], and to three alternative models of density-dependent trait evolution all taking biogeographic stochastic mapping into account. The Matching Competition model (MC) is a directional repulsive model where trait value of a species is drifting away from the trait value of each co-occurring lineage with a parameter *S* representing the competition strength [87]. The density-dependent linear model (DDlin) assumes a linear relationship between the evolutionary rate and the number of co-occurring lineages at time *t* with an intercept 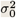 and a slope *r* [87]. The density-dependent exponential model (DDexp) assumes a linear relationship between the evolutionary rate and the number of cooccurring lineages at time *t* with an intercept 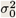 and a slope *r* [87]. We compared the models using Δ*AICc*. For each of these models, we used mean trait value per species and accounted for error in these estimates by incorporated the trait variance per species. We also performed sensitivity analysis by selecting 100 random datasets of biogeographic and ancestral host-use stochastic maps to replicate the model selection steps.

Finally, we investigated the influence that the host anemone has on clownfish morphology and color patterns. We used multivariate phylogenetic analysis of variance (MANOVA) [88, 89] to test the effect of host-use (*Radianthus* specialist, *Entacmaea* specialist, *Stichodactyla* specialist or generalist) on individual morphology and color. We first fitted linear models using penalized likelihood on generalized least square models (GLS) considering several alternative evolutionary models (BM, OU, EB or Pagel’s lambda) of the first four axes of the color PCA and the five morphology traits respectively. We compared the extended information criterion from the different evolutionary models with 100 bootstrap iterations [90] and applied phylogenetic multivariate analysis of variance on the best GLS model fit using Pillai test (Fig. 2).

## Acknowledgments

We thank Diego Hartasanchez and Thibault Latrille for comments on this manuscript.

## Declarations

- Funding The work was funded by a grant from the Swiss National Science Foundation to NS (grant 310030_185223) and from funding from the University of Lausanne.
- Conflict of interest/Competing interests (check journal-specific guidelines for which heading to use) We declare no conflicts of interest
- Availability of data and materials Raw reads are deposited at SRA (PRJNA1126266). Temporary link for reviewers https://dataview.ncbi.nlm.nih.gov/object/PRJNA1126266?reviewer=8a7h06nccmltpbmqa0gl9vnu4m Phenotypic and phylogenetic data are deposited as online supplementary material doi:10.5281/zenodo.12625956
- Code availability Code is available on the online supplementary material doi:10.5281/zenodo.12625956
- Authors’ contributions TG, BT and NS designed the study. AM, BM and SS gathered genomic data and ran phylogenetic analyses. AM and LF ran and interpreted gene selection analyses. AGJ gathered color data, BT gathered reproductive host associations data, TG gathered morphometrics data. TG ran and interpreted comparative phylogenetic and biogeographic analyses. TG wrote the manuscript with contributions of AM, BT and AGJ. All authors participated in reviewing, correcting, and approving the final version of the manuscript.

## Notes

https://dataview.ncbi.nlm.nih.gov/object/PRJNA1126266?reviewer=8a7h06nccmltpbmqa0gl9vnu4m

doi:10.5281/zenodo.12625956

### Competing Interest Statement

The authors have declared no competing interest.

