## Supplementary material for "Host-use Drives Convergent Evolution in Clownfish and Disentangles the Mystery of an Iconic Adaptive Radiation": S

#### Note on Reproductive-Host Associations

Our hypothesis is built off the following observations and rationale: It is well established clownfish species compete for "preferred" anemone hosts, some of which only serve as reproductive adult habitat, leaving the "less preferred" hosts to exclusively harbor juveniles and non-breeding subadults [1]. These preferences are thought to be partially driven by variation across host species in important characteristics such as size, tentacle shape, tentacle length, microhabitat, and toxicity. Host morphology has been shown to be linked to the distribution of adult and juvenile clownfish species across a reef, and also impact the ability of clownfish to shelter effectively at each life stage [1]. Host association dynamics are known to change over geographic space, as well as in the presence or absence of other clownfish species [2, 3]. Thus, 1) the frequency that clownfish associate with different host anemones on a reef captures the combined outcome of behavioral host preference and interspecific competition. 2) Natural selection acts on this outcome because different host characteristics impacts clownfish survival differently across all life stages. 3) Reproduction is clearly linked to the host anemone. Clownfishes lay and fertilize eggs on hard substrata directly next to or under anemone host tentacles, but not every host anemone species serves as adult reproductive habitat to every clownfish species. Those that do, typically fall into the "preferred" host species category. 4) Host imprinting impacts clownfish larval recruitment patterns, and clownfish larvae appear to prefer to recruit to the same species of host anemone their eggs were laid next to. 5) Less preferred hosts that only harbor juveniles and subadults are not true nursery habitats, and thus, do not have to be accounted for in phylogenetic analyses because they are not as ecologically and evolutionarily significant. To summarize, we hypothesize that adult reproductive host frequencies are capturing the most ecologically and evolutionarily significant aspects of the clownfish-sea anemone mutualism in a single variable: host preference, competition, reproduction, recruitment, growth, and survival. New host association data bin clownfishes into two broad host-use groups; host generalists and host specialists. Host generalists utilize $> 75\%$ of sea anemone hosts locally available in their respective ranges, while host specialists utilize $< 25\%$ of the sea anemone hosts locally available (Tab. 1).

##### Premnas Biaculeatus

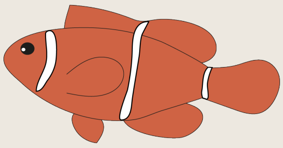

The spine-cheek anemonefish, *Premnas biaculeatus*, is famously known to only occupy solitary morphs of the bubble-tip anemone *Entacmaea quadricolor* across all life stages. Our data support prior host-use designations. Evaluation: *E. quadricolor* specialist.

##### Amphiprion akallopisos

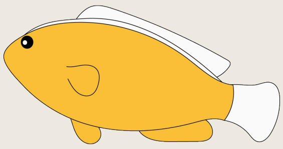

The skunk anemonefish, *Amphiprion akallopisos*, has only been documented in association with two anemone hosts: 1) the magnificent anemone *Radianthus magnifica*, and 2) Merten's carpet anemone *Stichodactyla mertensii*. Our dataset corroborates these prior associations. Further, while we find that both anemones host adult fish, our dataset shows that 94% of all *A. akallopisos* associations occur with *R. magnifica* at all life stages. Evaluation:*R. magnifica* specialist.

#### Amphiprion akindynos

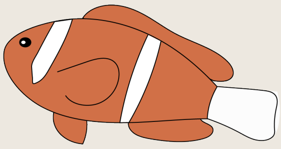

The Barrier Reef anemonefish has a complicated pattern of host use and exhibits variation in phenotype along its distribution, which is historically considered to stretch along the entire length of the Great Barrier Reef in Australia. In the southern extent of its range, it primarily hosts with the bubble-tip anemone *Entacmaea quadricolor*, takes on a darker orange body coloration, and exhibits thinner white stripes. In the North, body color tends to lighten, white stripes appear to be wider, and this species utilizes Merten's carpet anemone *Stichodactyla mertensii* and the letathery anemone *Radianthus crispa* most frequently- possibly due to the presence of other *E. quadricolor* specialists in this part of the GBR. We, and others, ultimately document adults associating with 5 host anemones, but primarily *E. quadricolor*, *R. crispa*, and *S. mertensii*. Evaluation: host generalist.

#### Amphiprion allardi

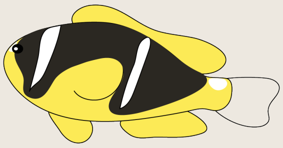

We provide major updates on patterns of host use for Allard's anemone fish. Our new data document three new host associations for this Western Indian Ocean species: 1) *Radianthus crispa*, 2) *Radianthus magnifica*, and 3) *Stichodactyla haddoni*. Adult fish are documented in association with four total host anemones (*E. quadricolor*, *R. magnifica, S. haddoni, S. mertensii*), and we designate all as primary adult hosts for this species. Evaluation: host generalist.

#### Amphiprion barberi

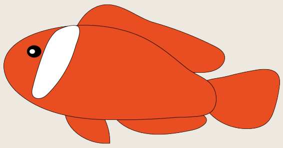

We document one new host association for Barber's anemonefish, *A. barberi*, via images from iNaturalist showing two occurrences of this species living with Merten's carpet anemone *Stichodactyla mertensii*. Otherwise, like all other members of the Tomato Clown Species Complex, adult *A. barberi* live primarily with the bubble tip anemone *Entacmaea quadricolor*. Individuals that associate with the leathery anemone *Radianthus crispa* are mostly juveniles. Evaluation: *E. quadricolor* specialist.

#### Amphiprion bicinctus

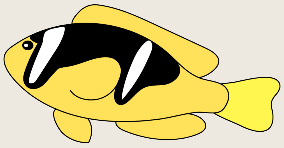

Historical misidentifications surrounding the sea anemone hosts in the Red Sea have plagued the host associations for the endemic Red Sea anemonefish, *Amphiprion bicintus*. Bennett-Smith et al. [4] recently revised these associations, showing that the giant carpet anemone *Stichodactyla gigantea*, long considered a host for this species, does not occur in the Red Sea. Instead, two other carpet anemones *S. haddoni* and *S. mertensii* are present. Our data show that a total of six anemone species (*E. quadricolor*, *H. aurora, R. crispa, R. magnifica, S. haddoni,* and *S. mertensii*) are found in association with *A. bicinctus*. Only *E. quadricolor, R. magnifica, S. haddoni*, and *S. mertensii* serve as adult habitat. Evaluation: host generalist.

#### Amphiprion chagosensis

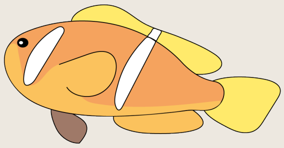

The Chagos anemonefish, *Amphiprion chagosensis*, is likely the most mysterious of all clownfish species. Endemic to the remote Chagos Archipelago South of the Maldives, they have been long thought to only host with the bubble tip anemone *Entacmaea quadricolor* [2, 5]. Recent scientific expeditions to the Chagos have provided photographic evidence that this species also hosts with *Radianthus crispa, R. magnifica,* and *Stichodactyla mertensii*. All but *R. crispa* appear serve as reproductive adult habitat for *A. chagosensis*. Further work on this species and its ecological associations are greatly needed. Evaluation: host generalist.

#### Amphiprion chrysogaster

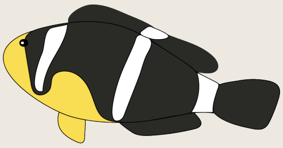

We document two new host associations between the Mauritian anemonefish, *Amphiprion chrysogaster*: 1) the bubble tip anemone *Entacmaea quadricolor*, and 2) the long tentacled anemone *Macrodactyla doreensis,* bringing the total number of sea anemone hosts for this species to six. However, our data could not independently confirm an association with the beaded anemone *Radianthus aurora* or Haddon's carpet anemone *Stichodactyla haddoni*. This is a poorly studied species and further work is needed on its ecological associations. However, we are confident that *E. quadricolor, R. magnifica,* and *S. mertensii* all serve as adult reproductive habitat. Evaluation: host generalist.

#### Amphiprion chrysopterus

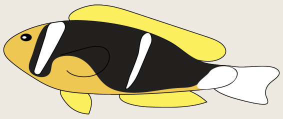

The orangefin anemonefish, *Amphiprion chysopterus*, has a large distributional range and as such, host associations in some parts of the range do not always match associations in others. However, in totality, we corroborate the host associations from prior studies, which list six anemone hosts. Our data show that *Radianthus crispa, R. magnifica,* and *Stichodactyla mertensii* serve as primary reproductive hosts, but *E. quadricolor* and *S. haddoni* can serve as adult hosts as well. Evaluation: host generalist.

#### Amphiprion clarkii

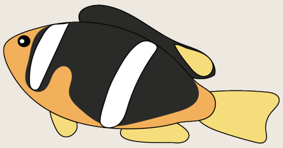

Clark's anemonefish, *Amphiprion clarkii* is the only species that is documented in association with all 10 host anemones. Our data corroborate this. Additionally, our data show adult *A. clarkii* associate with all host species except *Radianthus malu*. Evaluation: host generalist.

#### Amphiprion ephippium

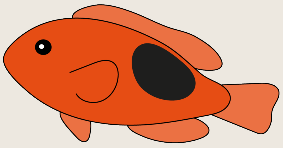

Part of the Tomato Clownfish Species Complex, the red saddleback anemonefish, *Amphiprion ephippium*, is only known to associate with the bubble-tip anemone *Entacmaea quadricolor* across all life stages. Our results corroborate this association. Evaluation: *E. quadricolor* specialist.

#### Amphiprion frenatus

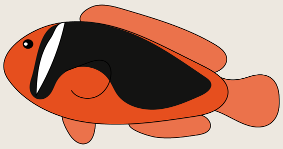

The Tomato Clownfish, *Amphiprion frenatus*, is only known to associate with the bubble-tip anemone *Entacmaea quadricolor* across all life stages. Our results corroborate this association. Evaluation: *E. quadricolor* specialist.

#### Amphiprion fuscocaudatus

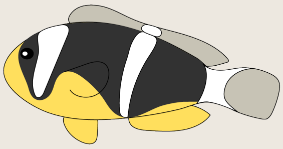

We provide major updates on patterns of host use for the Seychelles anemonefish, *Amphiprion fuscocaudatus*. Long thought to solely specialize on Merten's carpet anemone *Stichodactyla mertensii*, den Hartog [6] published updated host-use observations that have never been incorporated into the host-association literature. These include associations with 1) the bubble-tip anemone *Entacmaea quadricolor*, 2) *Radianthus aurora*, and 3) *Stichodactyla haddoni*. Our data corroborates den Hartog [6] and we also add a fourth new association with *Macrodactyla doreensis*. Taken together, *A. fuscocaudatus* is now known to associate with five species. Adults host in *S. mertensii, E. quadricolor,* and *S. haddoni*. Evaluation: host generalist.

#### Amphiprion latezonatus

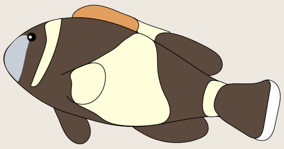

We provide major updates on patterns of host use for the wideband anemonefish, *Amphiprion latezonatus*. In their seminal field guide, Fautin and Allen [2] list only *Radianthus crispa* as a host for *A. latezonatus*. Scott et al., [7] updated host associations to include *Entacmaea quadricolor* and *Stichodactyla gigantea*. We further update associations for this species to include *S. haddoni* to bring the total number of hosts to four. We find adults in association with each of these species. While the majority of associations we observed for this species are with *E. quadricolor*, we interpret this pattern to be more of a reflection of this species' subtropical range in Southeast Australia, where *E. quadricolor* are very abundant, rather than a specialization on this host. Further, our observations of adults associating with multiple species in the genus *Stichodactyla*, as well as *R. crispa*, is a pattern we do not see in other *E. quadricolor* specialists. Evaluation: host generalist.

#### Amphiprion latifasciatus

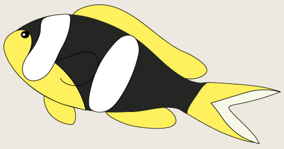

We provide major updates on patterns of host use for the Madagascar anemonefish, *Amphiprion latifasciatus*. Fautin and Allen [2] and Hoepner et al. [5] list *Stichodactyla mertensii* as the sole host for *A. latifasciatus*. Here we document four new sea anemone hosts for this species: 1) *Entacmaea quadricolor*, 2) *Radianthus aurora,* 3) *R. magnifica*, and 4) *S. haddoni*. We document each of these as hosts to adult *A. latifasciatus* anemonefish. Evaluation: host generalist.

#### Amphiprion mccullochi

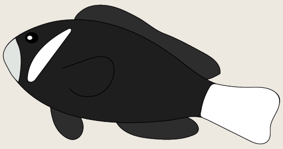

McCulloch's anemonefish, *Amphiprion mccullochi*, is a subtropical endemic species to Lord Howe Island, giving it the smallest range of any clownfish species. Our data corroborate prior host associations and show this species only hosts with *Entacmaea quadricolor*. Evaluation: *E. quadricolor* specialist.

#### Amphiprion melanopus

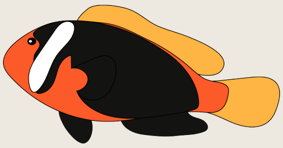

The red and black anemonefish, *Amphiprion melanopus*, has been documented in the literature to have been observed in association with four anemone hosts: *Entacmaea quadricolor, Radianthus crispa, R. magnifica,* and *S. gigantea* [5]. Here, we document only two: *E. quadricolor* and *R. crispa*. Among our newly collected data 98% of these observations occur in *E. quadricolor* at all life stages, which is the only host to harbor adult fish. We fail to observe this species in either *R. magnifica* or *S. gigantea*. Evaluation: *E. quadricolor* specialist.

#### Amphiprion nigripes

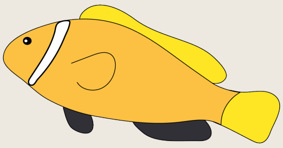

The blackfin anemonefish, *Amphiprion nigripes,* is endemic to the Maldives and has been previously documented to live solely with the magnificent anemone, *Radianthus magnifica*. Our data corroborate this prior host association. Evaluation: *R. magnifica* specialist.

#### Amphiprion ocellaris

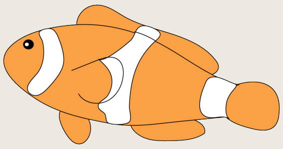

The Ocellaris clownfish, *Amphiprion ocellaris* has been previously documented with three host anemones: *Radianthus magnifica*, *Stichodactyla gigantea,* and *S. mertensii*. Our dataset only recovered *R. magnifica* and *S. gigantea* as hosts at all life stages- although we acknowledge there are plenty of photographs of this species in circulation demonstrating their associations with *S. mertensii* this does not appear to be one of their primary hosts. While *S. gigantea* appears to be an important host regionally, the majority of our observations across the entire range of this species show the majority of associations with *R. magnifica*. Evaluation: *R. magnifica* specialist.

#### Amphiprion omanensis

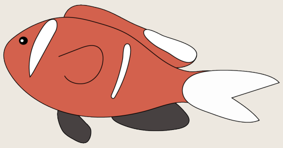

The Oman anemonefish, *Amphiprion omanensis*, is endemic to a small stretch of shallow water coastline in Oman that is famous for its seasonal upwelling and ephemeral kelp forests. It has been previously listed as having associations with three host species: *Entacmaea quadricolor, Radianthus crispa,* and *Stichodactyla haddoni*. This poorly studied species has limited photographic documentation. All verifiable host association information we could observe for this species was in *Entacmaea quadricolor*. Evaluation: *E. quadricolor* specialist.

#### Amphiprion pacificus

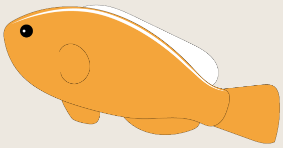

The Pacific anemonefish, *Amphiprion pacificus*, occupies a small geographic range surrounding Tonga and American Samoa. It was previously reported to host solely with *Radianthus magnifica*. Our data, while very limited, support this association. Evaluation: *R. magnifica* specialist.

#### Amphiprion percula

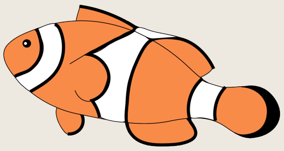

The true clownfish, *Amphiprion percula*, has long been listed as having associations with three anemone hosts: *Radianthus magnifica, R. crispa,* and *S. gigantea*. Our data recover associations with only *R. magnifica* and *S. gigantea* for fish at all life stages. While *S. gigantea* appears to be an important host regionally, *R. magnifica* serves as the primary host for the overwhelming majority of *A. percula*. Evaluation: *R. magnifica* specialist.

#### Amphiprion perideraion

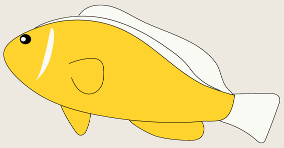

The pink-skunk anemonefish, *Amphiprion perideraion*, has long been listed as having associations with four host anemones: *Radianthus magnifica, R. crispa, Macrodactyla doreensis,* and *Stichodactyla gigantea*. Recently, Hoepner et al (2022) list *E. quadricolor* as a fifth host. Among the newly collected data here, *A. perideraion* had the largest sample size (N = 757 observations). We only observe associations with two hosts: *R. magnifica* and *R. crispa*. Of these 91% of all observations were in association with *R. magnifica*. Evaluation: *R. magnifica* specialist.

#### Amphiprion polymnus

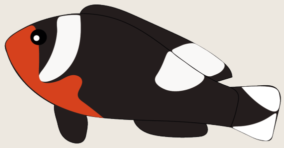

The saddle-back anemonefish, *Amphiprion polymnus*, has been previously listed as associating with three anemone hosts: *Stichodactyla haddoni, Radianthus crispa,* and *Macrodactyla doreensis*. Our dataset adds a new host association record with *H. aurora* but cannot confirm an association with *R. crispa*. Our dataset shows *S. haddoni* as the primary host for this species. Evaluation: *S. haddoni* specialist.

#### Amphiprion rubrocinctus

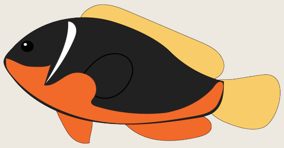

The Australian anemonefish, *Amphiprion rubrocintus*, has been previously listed as associating with two anemone hosts: *Entacmaea quadricolor* and *Stichodactyla gigantea*. Our dataset shows *Entacmaea quadricolor* as the sole host across all life stages. We could not independently confirm *S. gigantea* as a host. Evaluation: *E. quadricolor* specialist.

#### Amphiprion sandaracinos

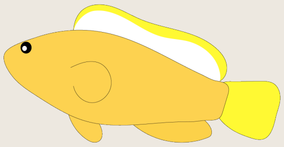

The orange skunk anemonefish, *Amphiprion sandaracinos*, has been previously listed as associating with two anemone host: *Stichodactyla mertensii* and *Radianthus crispa*. We confirm both of these as hosts in our dataset yet find 99% of all observations in association with *S. mertensii*. Evaluation: *S. mertensii* specialist

#### Amphiprion sebae

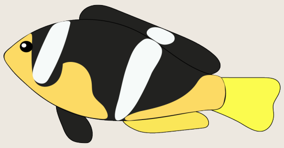

The Sebae anemonefish, *Amphiprion sebae*, has long been listed as solely hosting with *Stichodactyla haddoni*. Although few publicly available photographs exist, our dataset adds five new host associations: 1) *Entacmaea quadricolor,* 2) *Heteractis aurora,* 3) *Radianthus crispa*, 4) *Radianthus malu*, and 5) *Macrodactyla doreensis*. While sample sizes are low and the plurality are in association with *S. haddoni*, this pattern of association suggests host generalization. Evaluation: host generalist.

#### Amphiprion tricinctus

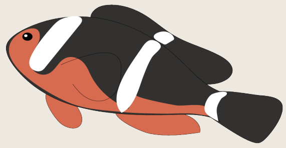

The three-band anemonefish, *Amphiprion tricinctus*, has been previously listed as having five anemone hosts: *Entacmaea quadricolor*, *Radianthus crispa, H. aurora, Stichodactyla mertensii,* and *Stichodactyla haddoni*. Our dataset confirms all but *R. crispa* as hosts and also adds three additional host anemones: 1) *R. magnifica,* 2) *R. malu*, and 3) *Macrodactyla doreensis* bringing the total number of hosts to eight. Evaluation: host generalist.

### Genomics

| Sample ID | Species | Sample Origin | Sequencing Year |
|---|---|---|---|
| NC016_AKI | A. akindynos | New Caledonia | 2017 |
| GA033_ALL | A. allardi | Comoro Islands | 2018 |
| GA071_BER | A. barberi | NA | 2018 |
| GA098_CHA | A. chagosensis | Chagos | 2017 |
| GA077_CRG | A. chrysogaster | Mauritius | 2018 |
| GA031_CRP | A. chrysopterus | Solomon Islands | 2018 |
| GA057_EPH | A. ephippium | Australia | 2018 |
| SC247_FUS | A. fuscocaudatus | Seychelles | 2018 |
| GA023_LAT | A. latezonatus | Australia | 2018 |
| MY284_LAT | A. latifasciatus | Mayotte | 2017 |
| GA056_MCC | A. mccullochi | Australia | 2018 |
| GB160_OCE | A. ocellaris | Indonesia | 2018 |
| GA051_OMA | A. omanensis | Oman | 2018 |
| GA069_PAC | A. pacificus | NA | 2018 |
| GB019_POL | A. polymnus | Indonesia | 2018 |
| GA070_RUB | A. rubrocinctus | NA | 2017 |
| GB033_SAN | A. sandaracinos | Indonesia | 2017 |
| GA058_TRI | A. tricinctus | Marshall Islands | 2018 |
| Ane19_PBI | P. biaculeatus | Australia | 2018 |

**Tab. S1**: Sample information for the individuals sequenced in this study. Raw sequencing data for these samples are available on SRA (PRJNA1126266). Raw sequencing data for the additional species was available from previous studies and was obtained from the SRA database: data for *Amphiprion akallopisos*, *A. bicinctus*, *A. melanopus*, *A. nigripes*, *A. sebae*, and the damselfish *Pomacentrus moluccensis* was obtained from PRJNA515163 [8] (samples 005_AKA, 006_BIC, 001_MEL, 004_NIG, 002_SEB and 1POMMO, respectively); data for *A. frenatus* was obtained from PRJNA433458 [9] (sample 001_FRE); data for *A. percula* and *A. perideraion* was obtained from PRJNA1022585 [10] (samples Ane03_PRC and GB059_PER, respectively); data for *A. clarkii* was obtained from PRJNA1025355 [11](sample GB081_CLA).

#### Sequencing and gene recovery

The number of obtained raw paired-end reads (PEs) for each individual ranged from ca. 61 to 158 million (Tab. S2). After trimming low-quality regions, the number of PEs per sample ranged from 57 to 151 million, corresponding to an estimated raw coverage between 6.5X and 17X (Tab. S2). Mapping of the reads resulted in ca. 23% to 36% of the reads mapping to *A. frenatus* protein-coding genes, and the number of SNPs retrieved for each species ranged from 242,351 for *A. ephippium* to 1,021,291 for *P. biaculeatus* (Tab. S2).

#### Phylogenetic reconstruction and dating

The species tree obtained with ASTRAL showed a considerable level of gene tree incongruence, as shown by the quartet support values (overall normalized quartet score of 0.76, Supplementary Fig. S1). Nevertheless, the tree was well supported, with all nodes having local posterior probabilities of 1 (Supplementary Fig. S1). The main clades of clownfishes were confirmed, and *Premnas biaculeatus* was placed as the basal lineage to all other clownfishes (Fig. S1). The dated phylogeny was also well supported, with all nodes with posterior probabilities of 1 and relatively narrow confidence intervals for the divergence times (Fig. S2). Divergence with the damselfish *P. moluccensis* occurred ca. 29 MYA, and the crown age of clownfishes was estimated to be ca. 10.5 MYA, with the main diversification occurring ca. 6.1 MYA (Fig. S2).

#### Simulation of positive selection

Simulations of convergent positive selection associated with shifts in hosts showed improved sensitivity to detect positive selection with increasing strength of selection (Supplementary Fig. S4). While the sensitivity was only 0.03 for a simulated $\omega$ of 2.0, it increased to 0.85 and 0.71 for *Entacmaea* and *Radianthus* shifts in case of a simulated $\omega$ of 100 (Fig. 4). When testing for positive selection on single branches, the sensitivity of the analyses drastically dropped, even for large $\omega$ (Supplementary Fig. S5, S6). Indeed, when considering the tests for positive selection on the longest branches, the highest sensitivity was 0.44, and it was obtained for a simulated $\omega$ of 100 (Supplementary Fig. S5). The sensitivity was further decreased for the tests on shorter branches, for which we had almost no power to detect positive selection (sensitivity ranging between 0 and 0.04; Supplementary Fig. S5, S6).

The false positives rates obtained from detecting a signal of convergent positive selection when no positive selection was simulated were low, with rates reaching maximum values of 0.03 (Fig. 4, Supplementary Fig. S4). False positives originating from positive selection occurring on a single internal branch but being wrongly recognized as convergent positive selection were also low, with maximum values reaching 0.05 (the “clade branch” scenario; Supplementary Fig. S7). However, false positive rates were increased when the positive selection was simulated only on the longest branches, with values reaching 0.17 (the “long branch” scenario; Supplementary Fig. S7).

#### Convergent positive selection associated with host shift

We tested for convergent positive selection associated with host shifts (Fig. 4) and found 165 and 8 potentially positively selected genes for shifts to *Entacmaea* and *Radianthus*, respectively (Tab. S3, S5). Because including the long branches potentially resulted in an increased number of false positives (Supplementary Fig. S7), we also performed more conservative tests (i.e., excluding the longest branches) and found a total of 76 and 3 genes under positive selection during shifts to *Entacmaea* and *Radianthus* hosts, respectively (Fig. 5, Tab. S3, S5).

Within the genes positively selected in shifts to *Radianthus*, we found the gene encoding for the protein crumbs homolog 1 (CRB1; UniProtKB ID: P82279) and the gene of the Serine protease FAM111A (FAM111A, UniProtKB ID: Q96PZ2). GO enrichment analysis for the genes positive selected in shifts to *Entacmaea* resulted in the enrichment of seven GO terms (Tab. S4), and within the significant genes, we observed the gene encoding for the protein $\beta$-carotene 15,15′-dioxygenase (BCO1; UniProtKB ID: Q9I993) and the gene for the olfactory receptor 11A1 (OR11A1, UniProtKB ID: Q9GZK7). Finally, the gene encoding for the retinitis-pigmentosa 1-like 1 protein (RP1L1, UniProtKB ID: Q8IWN7) was also found to be positively selected in shifts to *Entacmaea* hosts.

| Species | #Raw paired reads | #Trimmed paired reads | Estimated coverage | Mapped reads (%) | Number of SNPs |
|---|---|---|---|---|---|
| Newly Sequenced | | | | | |
| Amphiprion akindynos | 144468080 | 138544221 | 15.39 | 25.61 | 661’886 |
| Amphiprion allardi | 76442240 | 72228769 | 8.03 | 28.14 | 609’586 |
| Amphiprion barberi | 108546052 | 103969046 | 11.55 | 26.05 | 342’128 |
| Amphiprion chagosensis | 152079480 | 145485875 | 16.17 | 25.09 | 690’239 |
| Amphiprion chrysogaster | 89744484 | 84929015 | 9.44 | 26.17 | 643’108 |
| Amphiprion chrysopterus | 71462888 | 68101057 | 7.57 | 29.93 | 591’290 |
| Amphiprion clarkii | 86018456 | 79416119 | 8.82 | 25.70 | 611’680 |
| Amphiprion ephippium | 62946856 | 59487787 | 6.61 | 27.24 | 242’351 |
| Amphiprion fuscocaudatus | 91422892 | 87105870 | 9.68 | 27.10 | 646’589 |
| Amphiprion latezonatus | 65498508 | 61764821 | 6.86 | 26.11 | 728’488 |
| Amphiprion latifasciatus | 112477464 | 107293992 | 11.92 | 25.82 | 655’073 |
| Amphiprion mccullochi | 61348420 | 57387779 | 6.38 | 26.41 | 546’758 |
| Amphiprion ocellaris | 110898502 | 106144476 | 11.79 | 27.70 | 954’157 |
| Amphiprion omanensis | 84299930 | 80329429 | 8.93 | 27.95 | 634’039 |
| Amphiprion pacificus | 82251614 | 78243803 | 8.69 | 27.65 | 599’881 |
| Amphiprion percula | 100728420 | 96827262 | 10.76 | 28.41 | 919’552 |
| Amphiprion perideraion | 100637256 | 93022043 | 10.34 | 26.65 | 608’877 |
| Amphiprion polymnus | 73487236 | 70161629 | 7.8 | 28.51 | 614’260 |
| Amphiprion rubrocinctus | 157625280 | 150948795 | 16.77 | 26.28 | 372’012 |
| Amphiprion sandaracinos | 139139056 | 133257053 | 14.81 | 26.29 | 628’095 |
| Amphiprion tricinctus | 99952268 | 95795743 | 10.64 | 26.50 | 593’471 |
| Premnas biaculeatus | 108606766 | 103259564 | 11.47 | 22.65 | 1’021’291 |
| Previously sequenced (PRJNA515163) | | | | | |
| Amphiprion akallopisos | 463748734 | 426225412 | 47.36 | 25.84 | 651’649 |
| Amphiprion bicinctus | 413869256 | 386739640 | 42.97 | 25.32 | 721’376 |
| Amphiprion melanopus | 374346876 | 351324431 | 39.04 | 35.80 | 366’952 |
| Amphiprion nigripes | 458974486 | 422383027 | 46.93 | 24.97 | 708’753 |
| Amphiprion sebae | 313768062 | 298402708 | 33.16 | 26.94 | 709’255 |

**Tab. S2**: Sequencing, read processing, mapping and SNP calling statistics. Results are reported for the new samples and the samples retrieved from Marcionetti et al. ([8]; PRJNA515163). The total number of reads before and after quality filtering is reported. The estimated coverage is reported (read size: 100 bp, approximate reference genome size: 900 Mb). The percentage of mapped reads and the number of SNPs are also reported.

| Gene | Annotation | UniProtID | Gene Names |
|---|---|---|---|
| g32583.t1 | sp\|A2ASS6\|TITIN_MOUSE | A2ASS6 | Ttn |
| g39907.t1 | sp\|A4IF87\|GNPAT_BOVIN | A4IF87 | GNPAT |
| g11002.t1 | sp\|A4IFA3\|GT2D2_BOVIN | A4IFA3 | GTF2IRD2 |
| g50975.t1 | sp\|A4IFA3\|GT2D2_BOVIN | A4IFA3 | GTF2IRD2 |
| g41425.t1 | sp\|A7X406\|LECM1_PHIOL | A7X406 | NA |
| g35555.t1 | sp\|E9PYL2\|PRR12_MOUSE | E9PYL2 | Prr12 Kiaa1205 |
| g35556.t1 | sp\|E9PYL2\|PRR12_MOUSE | E9PYL2 | Prr12 Kiaa1205 |
| g54201.t1 | sp\|F1QRC1\|DRC1_DANRE | F1QRC1 | drc1 ccdc164 si:dkey-65b13.6 |
| g60372.t1 | sp\|O02839\|MCP_PIG | O02839 | CD46 MCP |
| g14082.t1 | sp\|O42249\|GBLP_ORENI | O42249 | gnb2l1 rack1 |
| g9184.t1 | sp\|O54750\|CP2J6_MOUSE | O54750 | Cyp2j6 |
| g27519.t1 | sp\|O75417\|DPOLQ_HUMAN | O75417 | POLQ POLH |
| g20072.t1 | sp\|O75443\|TECTA_HUMAN | O75443 | TECTA |
| g9758.t1 | sp\|O76031\|CLPX_HUMAN | O76031 | CLPX |
| g49223.t1 | sp\|P01133\|EGF_HUMAN | P01133 | EGF |
| g28343.t1 | sp\|P14784\|IL2RB_HUMAN | P14784 | IL2RB IL15RB |
| g55710.t1 | sp\|P15979\|HA1F_CHICK | P15979 | NA |
| g9544.t1 | sp\|P19180\|HV03_CARAU | P19180 | NA |
| g56300.t1 | sp\|P19256\|LFA3_HUMAN | P19256 | CD58 LFA3 |
| g38968.t1 | sp\|P20273\|CD22_HUMAN | P20273 | CD22 SIGLEC2 |
| g2209.t1 | sp\|P26285\|F262_BOVIN | P26285 | PFKFB2 |
| g20171.t1 | sp\|P38571\|LICH_HUMAN | P38571 | LIPA |
| g7242.t1 | sp\|P51608\|MECP2_HUMAN | P51608 | MECP2 |
| g32026.t1 | sp\|P55849\|DSC1_MOUSE | P55849 | Dsc1 |
| g8905.t1 | sp\|P70120\|HES5_MOUSE | P70120 | Hes5 Hes-5 |
| g32080.t1 | sp\|P78549\|NTH_HUMAN | P78549 | NTHL1 NTH1 OCTS3 |
| g45938.t1 | sp\|P97784\|CRY1_MOUSE | P97784 | Cry1 |
| g49787.t1 | sp\|Q00657\|CSPG4_RAT | Q00657 | Cspg4 Ng2 |
| g36803.t1 | sp\|Q00839\|HNRPU_HUMAN | Q00839 | HNRNPU C1orf199 HNRPU SAFA U21.1 |
| g36289.t1 | sp\|Q05556\|CP2AB_RABIT | Q05556 | CYP2A11 |
| g19145.t1 | sp\|Q08BC6\|ENO4_DANRE | Q08BC6 | eno4 zgc:153973 |
| g50240.t1 | sp\|Q08CS6\|MOXD2_DANRE | Q08CS6 | moxd2 moxd1l si:ch211-203k16.5 |
| g46992.t1 | sp\|Q09666\|AHNK_HUMAN | Q09666 | AHNAK PM227 |
| g23382.t1 | sp\|Q13018\|PLA2R_HUMAN | Q13018 | PLA2R1 CLEC13C |
| g60015.t1 | sp\|Q14126\|DSG2_HUMAN | Q14126 | DSG2 CDHF5 |
| g52450.t1 | sp\|Q14315\|FLNC_HUMAN | Q14315 | FLNC ABPL FLN2 |
| g43755.t1 | sp\|Q14AT5\|ANO7_MOUSE | Q14AT5 | Ano7 Ngep Tmem16g |
| g29958.t1 | sp\|Q15058\|KIF14_HUMAN | Q15058 | KIF14 KIAA0042 |
| g33612.t1 | sp\|Q15466\|NR0B2_HUMAN | Q15466 | NR0B2 SHP |
| g18186.t1 | sp\|Q15842\|KCNJ8_HUMAN | Q15842 | KCNJ8 |
| g35122.t1 | sp\|Q1JQB2\|BUB3_BOVIN | Q1JQB2 | BUB3 |
| g9680.t1 | sp\|Q1LY77\|SE1BA_DANRE | Q1LY77 | setd1ba setd1b si:dkey-237o15.4 |
| g39250.t1 | sp\|Q2Z1W2\|CENPU_CHICK | Q2Z1W2 | CENPU CENP50 MLF1IP |
| g16099.t1 | sp\|Q3UK37\|CN080_MOUSE | Q3UK37 | Tedc1 |
| g54746.t1 | sp\|Q3ZMH1\|SC5A8_DANRE | Q3ZMH1 | slc5a8 slc5a8l smcte zgc:152716 |
| g20109.t1 | sp\|Q4R9E0\|TECT3_MACFA | Q4R9E0 | TCTN3 TECT3 QtsA-10216 |
| g2343.t1 | sp\|Q502W7\|CCD38_HUMAN | Q502W7 | CCDC38 |
| g51034.t1 | sp\|Q5I0R6\|RL22L_XENTR | Q5I0R6 | rpl22l1 |
| g30789.t1 | sp\|Q5RIU9\|PORED_DANRE | Q5RIU9 | srd5a3 si:ch211-278f21.3 |
| g53593.t1 | sp\|Q5TCS8\|KAD9_HUMAN | Q5TCS8 | AK9 AKD1 AKD2 C6orf199 C6orf224 |
| g18063.t1 | sp\|Q5VW36\|FOCAD_HUMAN | Q5VW36 | FOCAD KIAA1797 |
| g51869.t1 | sp\|Q5W0U4\|BSPRY_HUMAN | Q5W0U4 | BSPRY |
| g28883.t1 | sp\|Q61656\|DDX5_MOUSE | Q61656 | Ddx5 Tnz2 |
| g35507.t1 | sp\|Q63184\|E2AK2_RAT | Q63184 | Eif2ak2 Prkr |
| g33501.t1 | sp\|Q6GVH4\|GGNB2_CHICK | Q6GVH4 | GGNBP2 ZNF403 RCJMB04_15h5 |
| g39065.t1 | sp\|Q6IND6\|HMCES_XENLA | Q6IND6 | hmces srapd1 |
| g60382.t1 | sp\|Q6J9G1\|STYK1_MOUSE | Q6J9G1 | Styk1 Nok |
| g29535.t1 | sp\|Q6NWF6\|K2C8_DANRE | Q6NWF6 | krt8 krt2-8 |
| g37503.t1 | sp\|Q6UXI9\|NPNT_HUMAN | Q6UXI9 | NPNT EGFL6L POEM UNQ295/PRO334 |
| g8568.t1 | sp\|Q6Y228\|LTOR3_PAGMA | Q6Y228 | lamtor3 |

| Gene | Annotation | UniProtID | Gene Names |
|---|---|---|---|
| g15983.t1 | sp\|Q6Y288\|B3GLT_HUMAN | Q6Y288 | B3GLCT B3GALTL B3GTL |
| g10744.t1 | sp\|Q7RTR2\|NLRC3_HUMAN | Q7RTR2 | NLRC3 NOD3 |
| g25736.t1 | sp\|Q7RTR2\|NLRC3_HUMAN | Q7RTR2 | NLRC3 NOD3 |
| g54659.t1 | sp\|Q7SXC6\|SLAIL_DANRE | Q7SXC6 | zgc:66447 |
| g26382.t1 | sp\|Q7TPD3\|ROBO2_MOUSE | Q7TPD3 | Robo2 Kiaa1568 |
| g12969.t1 | sp\|Q7TSA3\|BTLA_MOUSE | Q7TSA3 | Btla |
| g13944.t1 | sp\|Q7TSI1\|PKHM1_MOUSE | Q7TSI1 | Plekhm1 |
| g45834.t1 | sp\|Q803I8\|SYVN1_DANRE | Q803I8 | syvn1 hrd1 zgc:55735 zgc:77108 |
| g54124.t1 | sp\|Q86WI1\|PKHL1_HUMAN | Q86WI1 | PKHD1L1 |
| g22131.t1 | sp\|Q8AV61\|INSI1_DANRE | Q8AV61 | insig1 |
| g29330.t1 | sp\|Q8BHG1\|NRDC_MOUSE | Q8BHG1 | Nrdc Nrd1 |
| g35236.t1 | sp\|Q8BR93\|HARB1_MOUSE | Q8BR93 | Harbi1 |
| g48817.t1 | sp\|Q8BX90\|FND3A_MOUSE | Q8BX90 | Fndc3a D14Ertd453e Fndc3 Kiaa0970 |
| g26226.t1 | sp\|Q8BXA7\|PHLP2_MOUSE | Q8BXA7 | Phlpp2 Phlppl |
| g6970.t1 | sp\|Q8BYM5\|NLGN3_MOUSE | Q8BYM5 | Nlgn3 |
| g165.t1 | sp\|Q8CJ11\|AGRG2_RAT | Q8CJ11 | Adgrg2 Gpr64 Re6 |
| g37331.t1 | sp\|Q8I7P9\|POL5_DROME | Q8I7P9 | pol |
| g39693.t1 | sp\|Q8IWN7\|RP1L1_HUMAN | Q8IWN7 | RP1L1 |
| g27241.t1 | sp\|Q8IXT5\|RB12B_HUMAN | Q8IXT5 | RBM12B |
| g8163.t1 | sp\|Q8IYE1\|CCD13_HUMAN | Q8IYE1 | CCDC13 |
| g55469.t1 | sp\|Q8N3K9\|CMYA5_HUMAN | Q8N3K9 | CMYA5 C5orf10 DTNBP2 SPRYD2 TRIM76 |
| g5404.t1 | sp\|Q8N3R3\|TCAIM_HUMAN | Q8N3R3 | TCAIM C3orf23 TOAG1 |
| g25997.t1 | sp\|Q8NDN9\|RCBT1_HUMAN | Q8NDN9 | RCBTB1 CLLD7 E4.5 |
| g28973.t1 | sp\|Q8NHM4\|TRY6_HUMAN | Q8NHM4 | PRSS3P2 T6 TRY6 |
| g38621.t1 | sp\|Q8NHV1\|GIMA7_HUMAN | Q8NHV1 | GIMAP7 IAN7 |
| g45936.t1 | sp\|Q8QG61\|CRY1_CHICK | Q8QG61 | CRY1 |
| g7357.t1 | sp\|Q8TB03\|CX038_HUMAN | Q8TB03 | CXorf38 |
| g18659.t1 | sp\|Q8TD26\|CHD6_HUMAN | Q8TD26 | CHD6 CHD5 KIAA1335 RIGB |
| g13771.t1 | sp\|Q8TD55\|PKHO2_HUMAN | Q8TD55 | PLEKHO2 PLEKHQ1 PP9099 |
| g32480.t1 | sp\|Q8UWF0\|SC5A7_TORMA | Q8UWF0 | CHT1 |
| g57821.t1 | sp\|Q91ZI0\|CELR3_MOUSE | Q91ZI0 | Celsr3 |
| g50877.t1 | sp\|Q922H2\|PDK3_MOUSE | Q922H2 | Pdk3 |
| g54233.t1 | sp\|Q96IZ5\|RBM41_HUMAN | Q96IZ5 | RBM41 |
| g4249.t1 | sp\|Q96PZ2\|F111A_HUMAN | Q96PZ2 | FAM111A KIAA1895 |
| g52042.t1 | sp\|Q99928\|GBRG3_HUMAN | Q99928 | GABRG3 |
| g19673.t1 | sp\|Q99PG6\|TS1R1_MOUSE | Q99PG6 | Tas1r1 Gpr70 T1r1 Tr1 |
| g51790.t1 | sp\|Q99PP9\|TRI16_MOUSE | Q99PP9 | Trim16 Ebbp |
| g39602.t1 | sp\|Q9BGS7\|MOG_MACFA | Q9BGS7 | MOG QflA-14648 |
| g54147.t1 | sp\|Q9BQF6\|SENP7_HUMAN | Q9BQF6 | SENP7 KIAA1707 SSP2 SUSP2 |
| g60555.t1 | sp\|Q9BYJ1\|LOXE3_HUMAN | Q9BYJ1 | ALOXE3 |
| g28611.t1 | sp\|Q9CY64\|BIEA_MOUSE | Q9CY64 | Blvra |
| g11107.t1 | sp\|Q9D4J7\|PHF6_MOUSE | Q9D4J7 | Phf6 Kiaa1823 |
| g15821.t1 | sp\|Q9D666\|SUN1_MOUSE | Q9D666 | Sun1 Unc84a |
| g60053.t1 | sp\|Q9EPE9\|AT131_MOUSE | Q9EPE9 | Atp13a1 Atp13a |
| g39078.t1 | sp\|Q9GLY5\|ITIH3_RABIT | Q9GLY5 | ITIH3 |
| g5120.t1 | sp\|Q9GZK7\|O11A1_HUMAN | Q9GZK7 | OR11A1 OR11A2 |
| g51478.t1 | sp\|Q9H0U3\|MAGT1_HUMAN | Q9H0U3 | MAGT1 IAG2 PSEC0084 UNQ628/PRO1244 |
| g54915.t1 | sp\|Q9H2Y7\|ZN106_HUMAN | Q9H2Y7 | ZNF106 SH3BP3 ZFP106 ZNF474 |
| g42823.t1 | sp\|Q9H422\|HIPK3_HUMAN | Q9H422 | HIPK3 DYRK6 FIST3 PKY |
| g11594.t1 | sp\|Q9H9J4\|UBP42_HUMAN | Q9H9J4 | USP42 |
| g48006.t1 | sp\|Q9HCU0\|CD248_HUMAN | Q9HCU0 | CD248 CD164L1 TEM1 |
| g40477.t1 | sp\|Q9I993\|BCDO1_CHICK | Q9I993 | BCO1 BCDO BCMO1 |
| g39323.t1 | sp\|Q9IBD8\|PTPRC_CYPCA | Q9IBD8 | ptprc |
| g7361.t1 | sp\|Q9JHX4\|CASP8_RAT | Q9JHX4 | Casp8 |
| g23024.t1 | sp\|Q9NR16\|C163B_HUMAN | Q9NR16 | CD163L1 CD163B M160 UNQ6434/PRO23202 |
| g4118.t1 | sp\|Q9NRM0\|GTR9_HUMAN | Q9NRM0 | SLC2A9 GLUT9 |
| g50578.t1 | sp\|Q9NUV9\|GIMA4_HUMAN | Q9NUV9 | GIMAP4 IAN1 IMAP4 MSTP062 |
| g60462.t1 | sp\|Q9NZN9\|AIPL1_HUMAN | Q9NZN9 | AIPL1 AIPL2 |
| g24824.t1 | sp\|Q9P0M4\|IL17C_HUMAN | Q9P0M4 | IL17C UNQ561/PRO1122 |
| g13333.t1 | sp\|Q9QXX8\|NUFP1_MOUSE | Q9QXX8 | Nufip1 |
| g19389.t1 | sp\|Q9UK61\|TASOR_HUMAN | Q9UK61 | TASOR C3orf63 FAM208A KIAA1105 |

| Gene | Annotation | UniProtID | Gene Names |
|---|---|---|---|
| g49278.t1 | sp\|Q9UQE7\|SMC3_HUMAN | Q9UQE7 | SMC3 BAM BMH CSPG6 SMC3L1 |
| g28370.t1 | sp\|Q9W6V5\|PTPRJ_CHICK | Q9W6V5 | PTPRJ |
| g23998.t1 | sp\|Q9Y2G3\|AT11B_HUMAN | Q9Y2G3 | ATP11B ATPIF ATPIR KIAA0956 |
| g37726.t1 | NA | NA | NA |
| g37333.t1 | NA | NA | NA |
| g40543.t1 | NA | NA | NA |
| g40986.t1 | NA | NA | NA |
| g41444.t1 | NA | NA | NA |
| g54519.t1 | NA | NA | NA |
| g56094.t1 | NA | NA | NA |
| g9953.t1 | NA | NA | NA |
| g36641.t1 | NA | NA | NA |
| g35574.t1 | NA | NA | NA |
| g4617.t1 | NA | NA | NA |
| g58238.t1 | NA | NA | NA |
| g38877.t1 | NA | NA | NA |
| g42164.t1 | NA | NA | NA |
| g60891.t1 | NA | NA | NA |
| g4615.t1 | NA | NA | NA |
| g5935.t1 | NA | NA | NA |
| g17993.t1 | NA | NA | NA |
| g31107.t1 | NA | NA | NA |
| g37989.t1 | NA | NA | NA |
| g16112.t1 | NA | NA | NA |
| g18524.t1 | NA | NA | NA |
| g32125.t1 | NA | NA | NA |
| g39334.t1 | NA | NA | NA |
| g29607.t1 | NA | NA | NA |
| g33802.t2 | NA | NA | NA |
| g40179.t1 | NA | NA | NA |
| g41924.t1 | NA | NA | NA |
| g1667.t1 | NA | NA | NA |
| g19724.t1 | NA | NA | NA |
| g36153.t1 | NA | NA | NA |
| g42764.t1 | NA | NA | NA |
| g11975.t1 | NA | NA | NA |
| g14916.t1 | NA | NA | NA |
| g26520.t1 | NA | NA | NA |
| g362.t1 | NA | NA | NA |
| g12769.t1 | NA | NA | NA |
| g9746.t1 | NA | NA | NA |
| g54534.t1 | NA | NA | NA |
| g59417.t1 | NA | NA | NA |
| g7606.t1 | NA | NA | NA |

**Tab. S3**: List of genes positively selected in branches corresponding to a specialisation event towards *Entacmea* reproductive association. Red-highlighted rows correspond to genes found to positively selected in analyses excluding the basal branch of *Premnas biaculeatus*

| GO.ID | Term | classicFisher |
|---|---|---|
| GO:0048712 | negative regulation of astrocyte differentiation | 0.00073 |
| GO:0009394 | 2′-deoxyribonucleotide metabolic process | 0.00088 |
| GO:0043517 | positive regulation of DNA damage response | 0.00088 |
| GO:0035357 | peroxisome proliferator activated receptor signaling pathway | 0.00121 |
| GO:0032369 | negative regulation of lipid transport | 0.00179 |
| GO:0043651 | linoleic acid metabolic process | 0.00328 |
| GO:0006757 | ATP generation from ADP | 0.00359 |

**Tab. S4**: Go enriched analysis results for the genes under positive selection in branches linked to specialisation to *Entacmaea* reproductive association

| Gene | Annotation | UniProtID | Gene Names |
|---|---|---|---|
| g4249.t1 | sp\|Q96PZ2\|F111A_HUMAN | Q96PZ2 | Serine protease FAM111A |
| g50522.t1 | sp\|Q80XC3\|US6NL_MOUSE | Q80XC3 | USP6 N-terminal-like protein |
| g4184.t1 | sp\|P82279\|CRUM1_HUMAN | P82279 | Protein crumbs homolog 1 |
| g45007.t1 | sp\|P35031\|TRY1_SALSA | P35031 | Trypsin-1 |
| g1988.t1 | sp\|P13942\|COBA2_HUMAN | P13942 | Collagen alpha-2(XI) chain |
| g36611.t1 | sp\|O43157\|PLXB1_HUMAN | O43157 | Plexin-B1 |
| g12194.t1 | NA | NA | NA |
| g6554.t1 | sp\|A7MD70 \|SPDLY_DANRE | A7MD70 | Protein Spindly |

**Tab. S5**: List of genes positively selected in branches corresponding to a specialisation event towards *Radianthus* reproductive association. Red-highlighted rows correspond to genes found to positively selected in analyses excluding the basal branch of *Amphiprion percula* and *ocellaris*

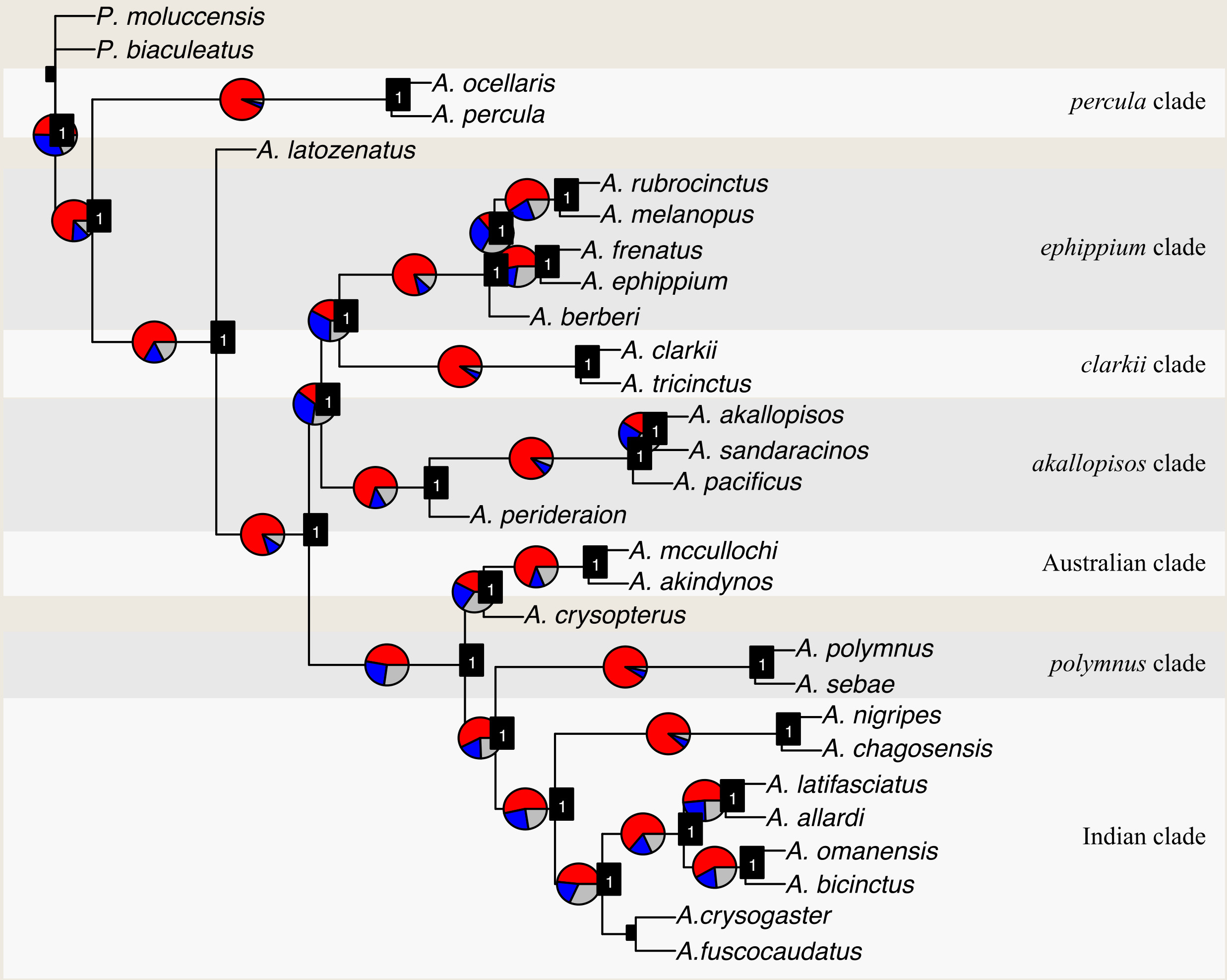


**Fig. S1**: Clownfishes' ASTRAL-III tree. Pie charts represent the proportion of the gene trees agreeing with the topology of the main species tree (red) and the two other alternative topologies (blue and grey). The local posterior probabilities for the branches are reported in black squared at each node. Plotting was done with the R package AstralPlane (v.0.1.1; https://github.com/chutter/AstralPlane.git). The main clownfish clades are reported [12].

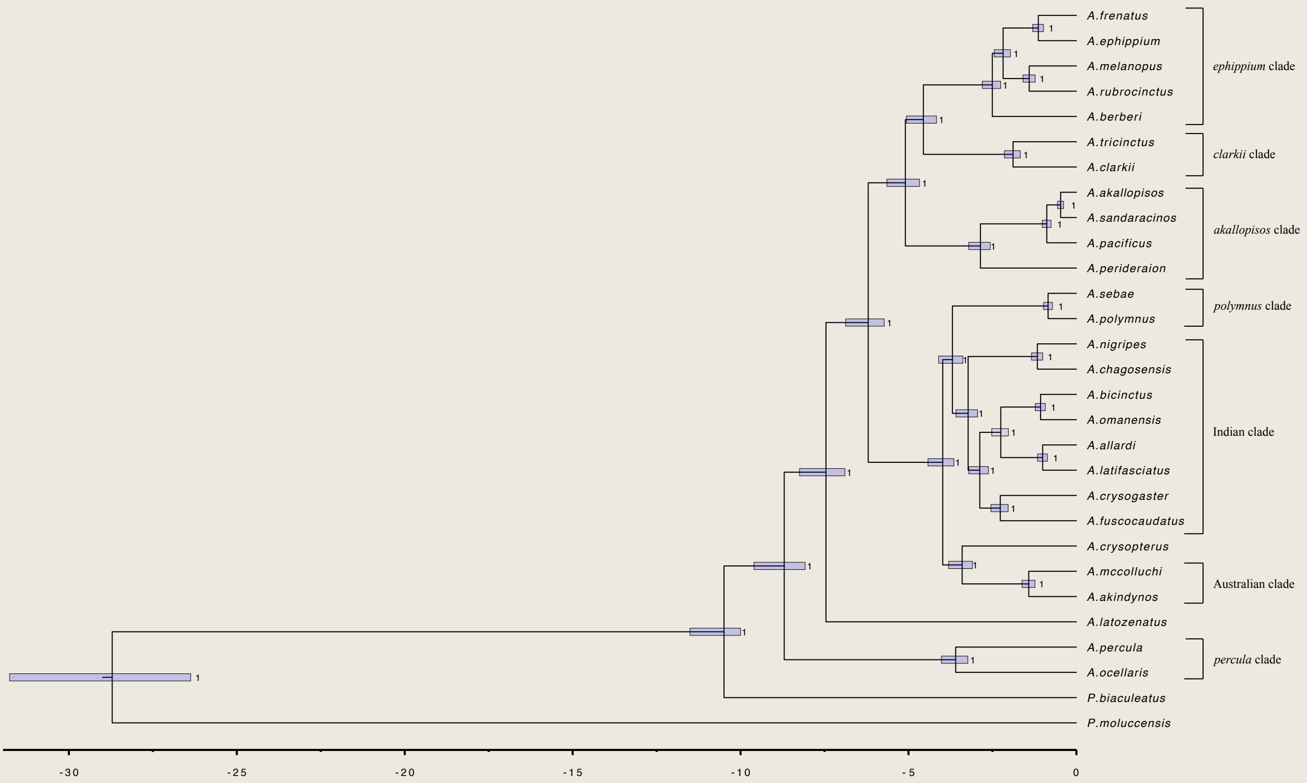


**Fig. S2**: Clownfishes' dated phylogeny. The dated phylogeny was obtained with BEAST2 (v.2.6.3) using the 20 most informative genes. Error bars represent the 95% HPD (height posterior density) confidence interval for the divergence time estimates. Numbers at the nodes indicate the Bayesian posterior probability. The time scale is given in million years before the present (MYA). Main clownfish clades are reported [12]

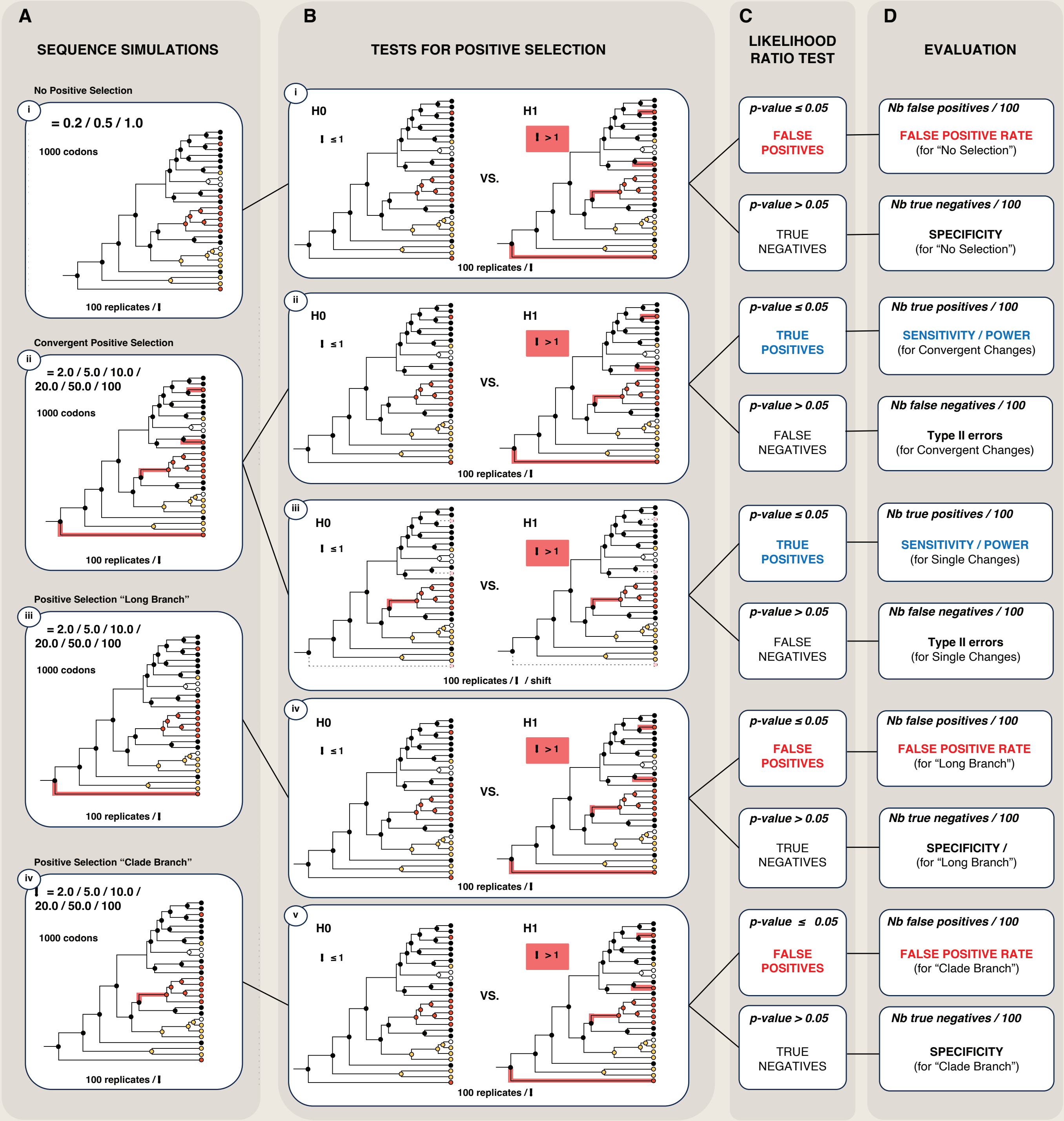


**Fig. S3**: Scheme of the simulations performed to examine sensitivity, specificity, and false positive rate of positive selection (PS) analyses. The simulations are depicted for the shifts to *Entacmaea* sea anemone host, but the same approach was employed for shifts to *Radianthus*. A) Sequences of 1,000 codons were simulated under the branch-site model for different evolutionary scenarios (i-iv), using the species tree and following the identified shifts of sea anemone hosts. We simulated no ($\omega = 0.2, 0.5, 1.0$) or increasing strength ($2 \leq \omega \leq 100$) of "convergent PS", i.e., PS on branches with convergent host shifts (ii). We also simulated an increasing strength of PS on specific branches: a "long branch" (iii) and a "clade branch" (iv) with host shifts. For each scenario and each $\omega$, we simulated 100 sequence alignments (i.e., 100 replicates). B) PS analyses were performed for each simulated alignment using the branch-site model implemented in codeml and the species tree. The H0 model - with $\omega$ constrained to be smaller or equal to 1- was compared to the H1 model, where $\omega$ was estimated and forced to be larger than 1 in the foreground branches (in red). For all simulated scenarios, we tested for "convergent PS" (i, ii, iv, v). For the scenarios simulated with increasing strength of "convergent PS", we also tested for PS on individual branches with a host shift (after purging the additional PS branches, as reported by the dotted lines in figure iii). C, D) The best model was determined with a likelihood ratio test (LTR; df=1). The significance of the LRT, together with the information of the simulated scenario, allowed the calculation of the sensitivity (true positives/total number of tests), specificity (true negatives/ total number of tests), and false positive rate (false positives/total number of tests).

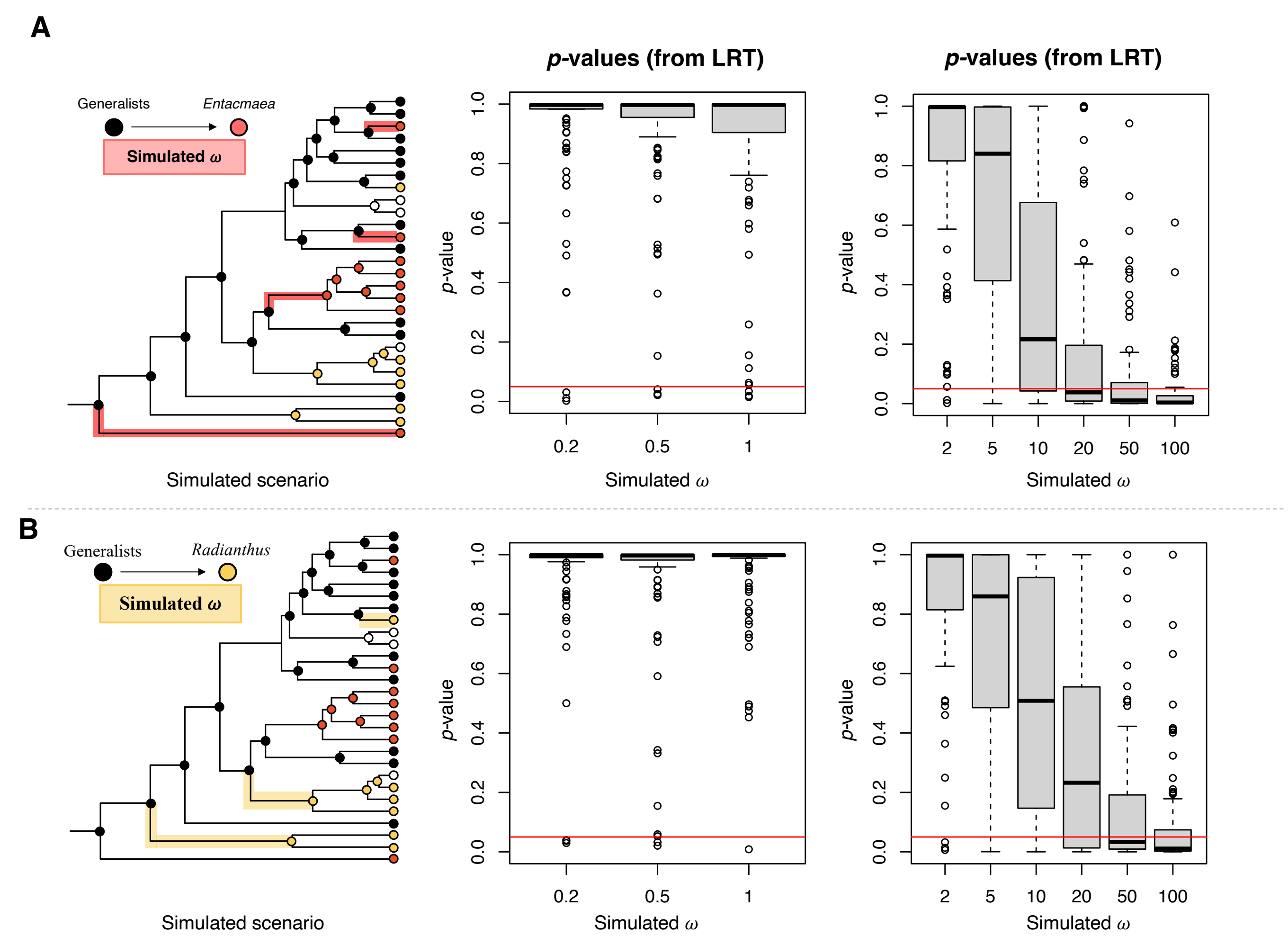


**Fig. S4**: p-values obtained for the likelihood ratio test (LRT) between the model H0 (no positive selection) and H1 (convergent positive selection) performed on simulated sequences. Sequences were simulated under scenarios of presence ($\omega > 1$) or absence ($\omega \leq 1$) of convergent positive selection associated with shifts to *Entacmaea* (A) or *Radianthus* (B) sea anemone hosts. For each $\omega$ value, we simulated 100 sequences with a length of 1000 codons. Tests for positive selection were performed with the branch-site model implemented in codeml (PAML, [13]), and simulations were performed with evolver (PAML, [13]). Red lines in the plots correspond to the significance level (a = 0.05). For scenarios simulated with no positive selection ($\omega \leq 1$), the LRT is expected to be non-significant, and the proportion of significant tests determines the false positive rate (Fig. 5). Contrarily, for scenarios simulated with convergent positive selection ($\omega > 1$), the LRT is expected to be significant, and the proportion of significant tests determines the sensitivity (Fig. 5).

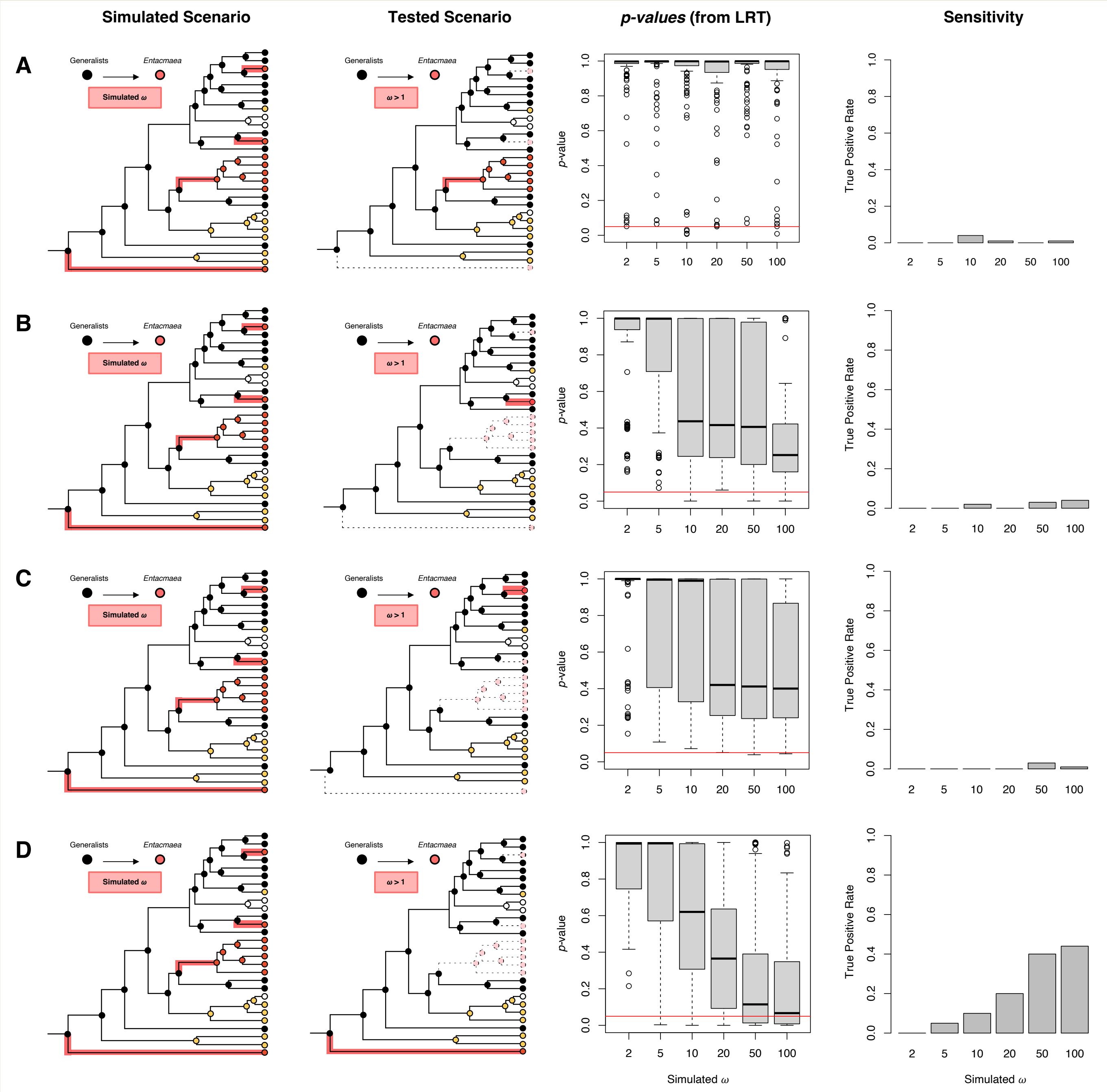


**Fig. S5**: Sensitivity (power) for the detection of positive selection on a single branch with a shift to *Entacmaea* sea anemone host. A-D, first panel) Sequences were simulated under scenarios of increasing strength positive selection ($2 < \omega < 100$) associated with shifts to *Entacmaea* hosts. Colored branches represent foreground branches in simulations. Simulations were performed with evolver (PAML, [13]). For each $\omega$ value, we simulated 100 sequences with a length of 1000 codons. A-D, second panel) Simulated sequences were tested for positive selection on single branches, as depicted in the schemes. The colored branch was set as the foreground, and the dotted branches were removed from the analyses. A-D, third panel) p-values obtained for the likelihood ratio test (LRT) between the model H0 (no positive selection) and H1 (positive selection on foreground branches) performed on the simulated data. Tests for positive selection were performed with the branch-site model implemented in codeml (PAML, [13]). Red lines in the plots correspond to the significance level (a = 0.05). A-D, fourth panel) Power was obtained as the proportion of true positives (simulated $\omega > 1$, estimated $\omega > 1$, p-value LTR < 0.05) for each $\omega$ value and considered scenario.

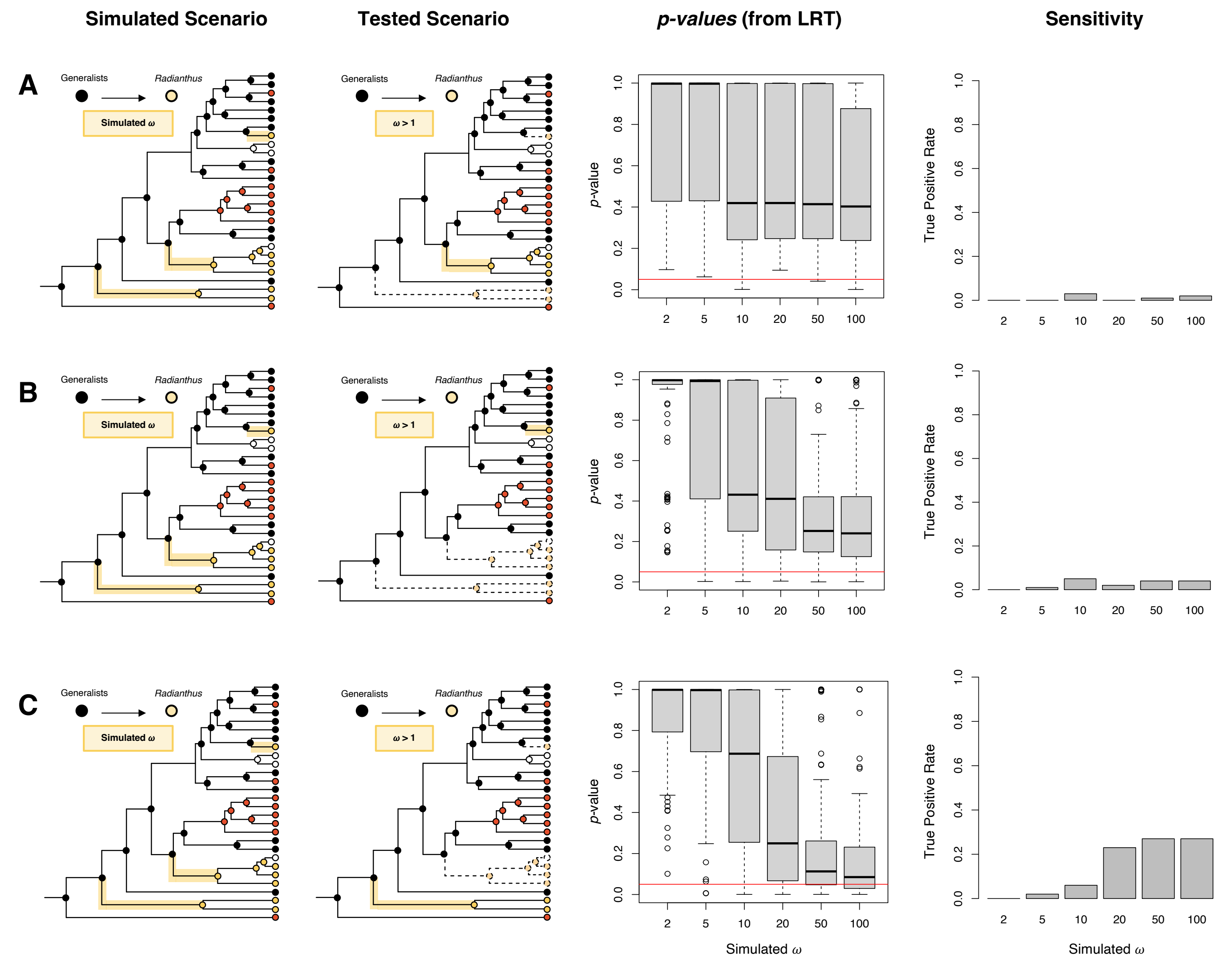


**Fig. S6**: Sensitivity (power) for the detection of positive selection on a single branch with a shift to *Radianthus* sea anemone host. A-D, first panel) Sequences were simulated under scenarios of increasing strength positive selection ($2 < \omega < 100$) associated with shifts to *Radianthus* hosts. Colored branches represent foreground branches in simulations. Simulations were performed with evolver (PAML, [13]). For each $\omega$ value, we simulated 100 sequences with a length of 1000 codons. A-D, second panel) Simulated sequences were tested for positive selection on single branches, as depicted in the schemes. The colored branch was set as the foreground, and the dotted branches were removed from the analyses. A-D, third panel) p-values obtained for the likelihood ratio test (LRT) between the model H0 (no positive selection) and H1 (positive selection on foreground branches) performed on the simulated data. Tests for positive selection were performed with the branch-site model implemented in codeml (PAML, [13]). Red lines in the plots correspond to the significance level (a = 0.05). A-D, fourth panel) Power was obtained as the proportion of true positives (simulated $\omega > 1$, estimated $\omega > 1$, p-value LTR < 0.05) for each $\omega$ value and considered scenario.

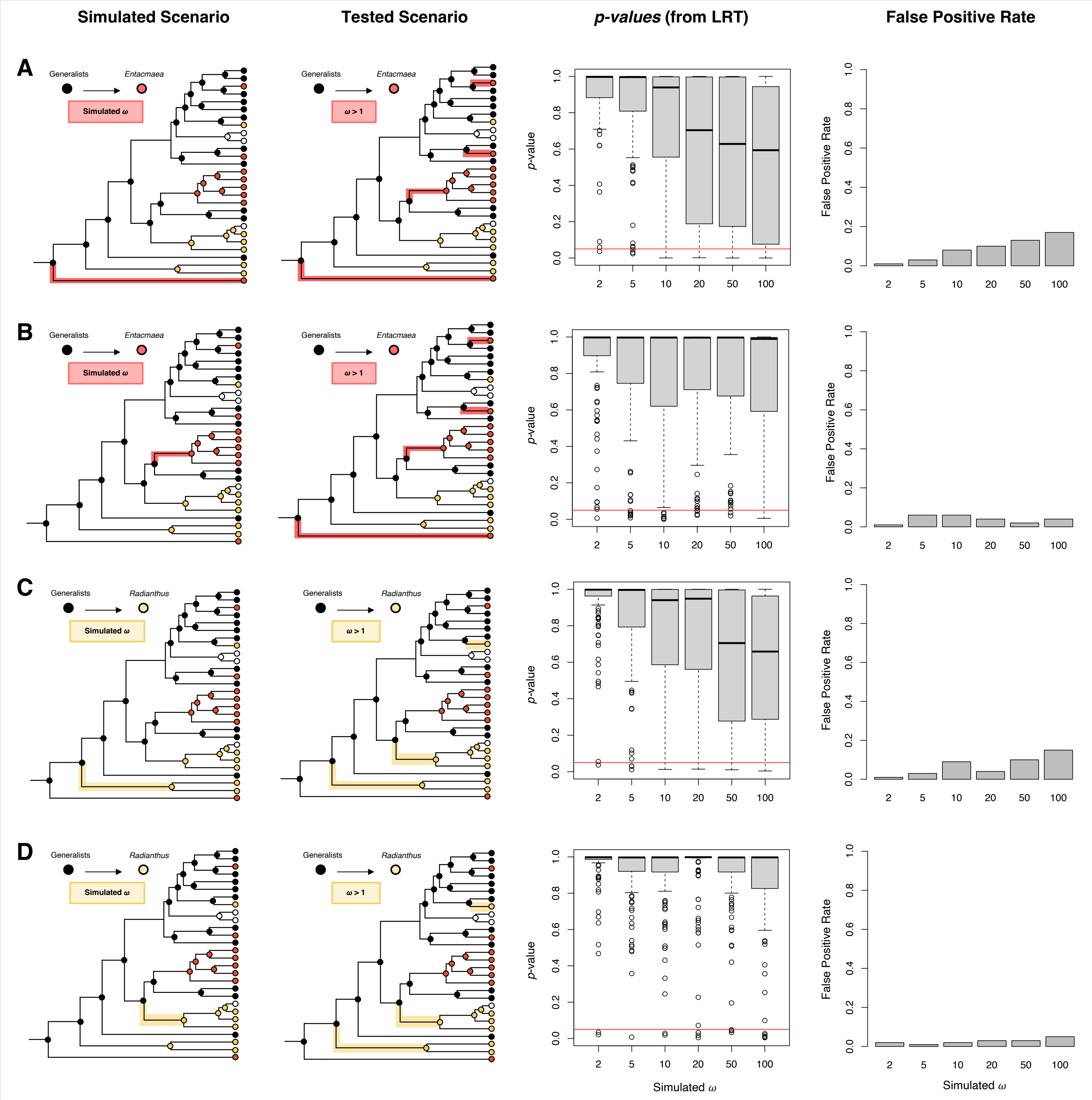


**Fig. S7**: False positive rate for the detection of convergent positive selection originating from positive selection on a single long branch (A, C) or on a "clade" branch (B, D). Shifts from generalists to *Entacmaea* (A, B) or *Radianthus* (C, D) were considered. A-C, first panel) Sequences were simulated under scenarios of increasing strength positive selection ($2 < \omega < 100$) on the longest branch with shifts to *Entacmaea* (A) or *Radianthus* (C) hosts, and on the branch leading to a clade of *Entacmaea* (B) or *Radianthus* (D) specialists (i.e., "clade branch"). For each $\omega$ value and scenario, we simulated 100 sequences with a length of 1000 codons. Simulations were performed with evolver (PAML, [13]). A-C, second panel) Simulated sequences were tested for convergent positive selection linked with shift to *Entacmaea* (A, B) or *Radianthus* (C, D) hosts. The colored branches were set as the foreground. A-D, third panel) p-values obtained for the likelihood ratio test (LRT) between the model H0 (no positive selection) and H1 (convergent positive selection) performed on the simulated data. Tests for positive selection were performed with the branch-site model implemented in codeml (PAML, [13]). Red lines in the plots correspond to the significance level (a = 0.05). A-D, fourth panel) False positive rate was obtained as the proportion of false positives (simulated $\omega$ on a single specific branch > 1, estimated $\omega$ on all foreground branches > 1, p-value LTR < 0.05) for each $\omega$ value and scenario.

#### Biogeography

Using Species Distribution Modelling [14], we observe that clownfish communities are composed by similar phenotypic assemblages. Fig. S8 illustrates that clownfish species presenting similar color patterns rarely co-occur in the same communities. Further, we observe that species specialized in *Entacmaea quadricolor* (Fig. S9) and *Radianthus magnifica* (Fig. S10) only co-occur when distantly related in the phylogeny and presenting divergent white bar patterning. Our Biogeographic reconstruction based on the new phylogeny and updated species distributions confirm the findings from Litsios et al. [15] of a Indo-Australian Archipelago ancestor and a stepping-stone colonization of marginal Indo-Pacific regions (Fig. S13).

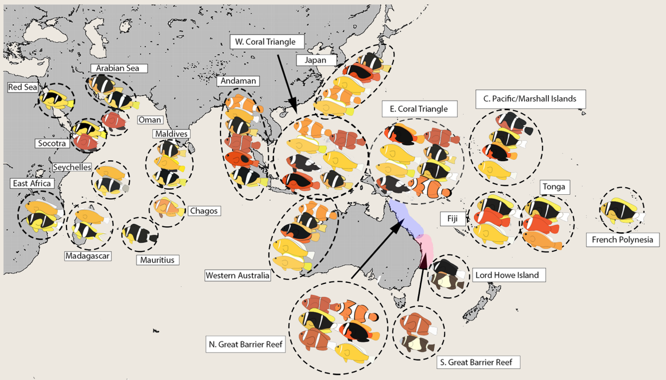


**Fig. S8**: Similar clownfishes communities in terms of coloration and host-use are identified in different areas of the Indo-Pacific

| Model | $\Delta AIC$ |
|---|---|
| DEC | 6.77 |
| DEC+J | 5.11 |
| DIVA | 0 |
| DIVA+J | 1.99 |
| BayArea | 33.59 |
| BayArea+J | 14.69 |

**Tab. S6**: AIC comparison between biogeographic models.

#### Discussion: Phenotyping

##### Morphology

We estimated the morphology of 214 adult clownfish individuals based on museum standardised pictures. We used pictures of the left side of samples and placed 29 landmarks (Fig. S14). We can see from the landmark positions (Fig. S15) that some landmarks (4, 7, 15,16) are more affected than others by reproductive association categorization. Landmarks placed on gills, mouth, eyes, pectoral fins and caudal fins seem unaffected. However, landmarks placed at the pelvic and dorsal fins seem more divergent between categories. This affects estimates of body ratio and peduncle ratio as well as snout angle (Fig. S17) and generates the variation along PC1 axis of the PCA that is driven

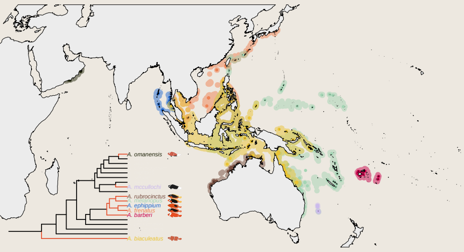

**Fig. S9**: Geographical distribution of Entacmaea specialists. Shaded areas are estimated species distributions. Dots represent occurrences. Red branches in the phylogenetic tree represent lineages estimated to be Entacmaea specialists

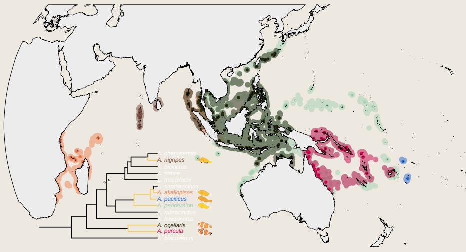

**Fig. S10**: Geographical distribution of Radianthus specialists. Shaded areas are estimated species distributions. Dots represent occurrences. Yellow branches in the phylogenetic tree represent lineages estimated to be Radianthus specialists

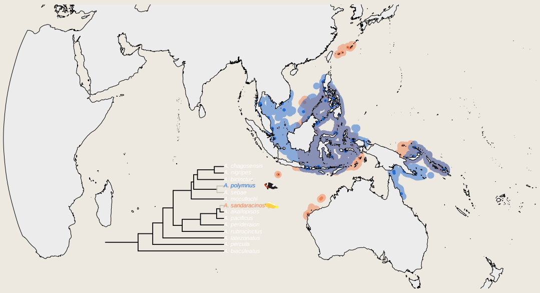

**Fig. S11**: Geographical distribution of Stichodactyla specialists. Shaded areas are estimated species distributions. Dots represent occurrences. Grey branches in the phylogenetic tree represent lineages estimated to be Stichodactyla specialists

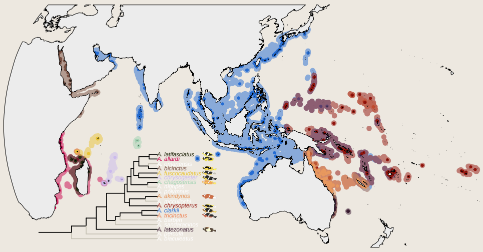

**Fig. S12**: Geographical distribution of generalists. Shaded areas are estimated species distributions. Dots represent occurrences. Black branches in the phylogenetic tree represent lineages estimated to be generalists

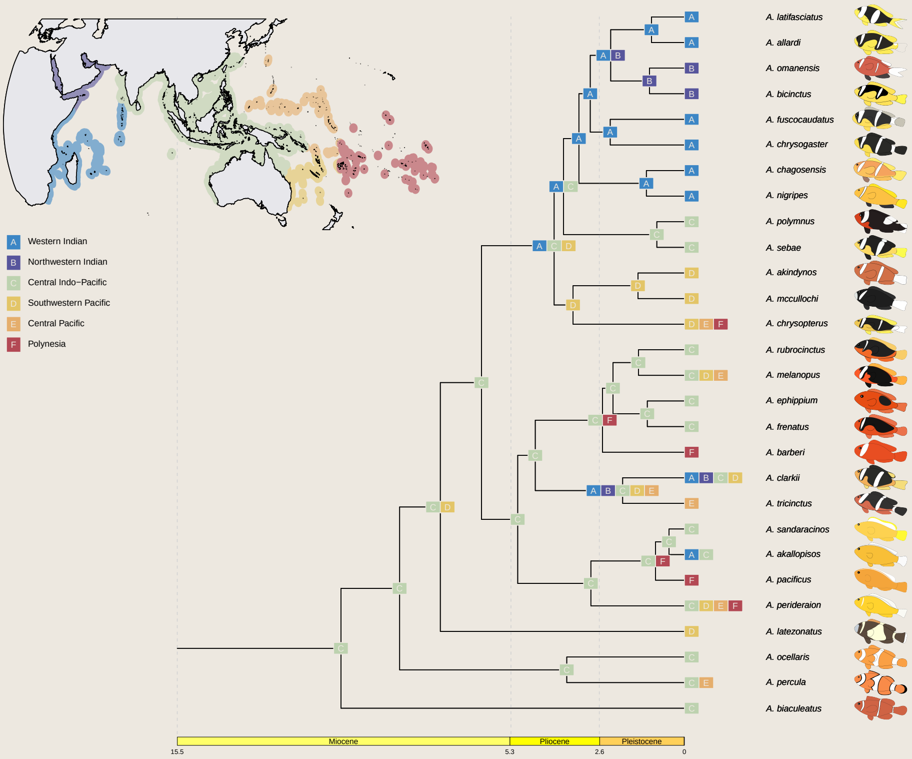


**Fig. S13**: Results of the ML estimation of biogeography ancestral state using the DIVA model.

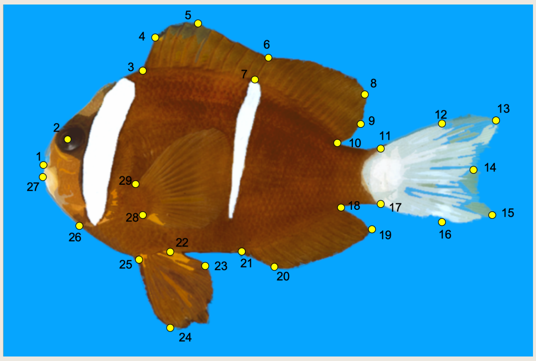

**Fig. S14**: Position of the 29 landmarks on a *Amphiprion akindynos* specimen

by elongation (Fig. S16). Individuals with small values for PC1 are more elongated and individuals with large values are deep-bodied. PC2 discriminates individuals with a smaller snout-angle (small values) to those with a stronger snout angle (large values). PC3 represents variation around the caudal fin with larger lower parts for small values along the axis and thinner lowers parts for large values. Finally PC4 represents variation along the pelvic fins with lower pelvic fins for small values and higher pelvic fins for large values.

#### Color patterns

Two types of analyses have been done on color patterns. First, images were separated in three channels (Red, Green, Blue). We concatenated the three RGB layers and performed a PCA on the matrix. With this approach, each pixel was contributing three times to PCA axis variation. The first four axis, that were used in the following analyses represented 47% of the total variation. To understand color patterns variation along axes, we summed respectively the negative and positive contributions of each pixel to each axis. For the graphic representations of PCAs on the RGB dataset, we represented the strength of pixel-wise negative contribution with red intensity and the strength of pixel-wise positive contribution with blue intensity. We observe that off-stripe patterns drive the negative contribution to PC1 while the head stripe drives the positive contribution to PC1. Individuals with low PC1 values are characterized by bright orange off-stripe patterns and by an absence of a white head stripe. Individuals with high PC1 values are characterized by dark off-stripe patterns and the presence of a white head stripe. PC2 is driven by dorsal fin brightness for negative contribution and the presence of a second white bar for positive contribution. PC3 is driven by bright caudal fin and ventral patterns for negative contribution and the width of the second stripe for positive contribution. Finally, PC4 is driven by the width of the head stripe for negative contribution and by dorsal patterns for the positive contribution.

Second, we identified color patterns from three specific color channels using the R package *Patternize* [16], black (RGB color code: 0,0,0), orange (RGB color code: 255,165,0) and white (RGB color code: 255,255,255). We performed a PCA for each channel independently and quantified the contribution of each pixel to each PCA axis. For black patterns, the first four axes explain 53.9% of the variation. For orange patterns, the first four axes explain 49% of axis variation. For white patterns, the first four axes explain 45.1% of the variation. For both black and orange patterns PC1 is driven by whole body coloration excluding stripes while PC2 is driven by fin coloration. For white patterns, PC1 is driven by the three stripe patterns and dorsal patterns. PC2 is driven by off-stripe patterns and the caudal fin.

#### Discussion: Phylogenetic Comparative Methods

##### Color patterns by color:

Using multivariate comparative approaches, we found phylogenetic signal for white and black color patterns (Tab. S7). However, phylogenetic independence could not be rejected for orange patterns (Tab. S7). With this approach, we find an association with white patterns evolutionary optima and reproductive host association. By fitting phylogenetic MANOVAs [17], using the best selected model from each dataset, we found a significant association between host association and white (Pillai: $S = 0.8506, p = 0.003$) and black (Pillai: $S = 0.8218, p = 0.006$) patterns, but not for orange patterns (Pillai: $S = 0.5915, p = 0.112$).

Using univariate analyses, white patterns best fit a OUM model (optima driven by reproductive associations) for the two first PC axes (tab. S8) and black patterns best fit an OU model (single optimum) for PC1 and a BMM model (evolutionary rates driven by reproductive host associations) for PC2 S10. The fit to PC2 axis of black patterns

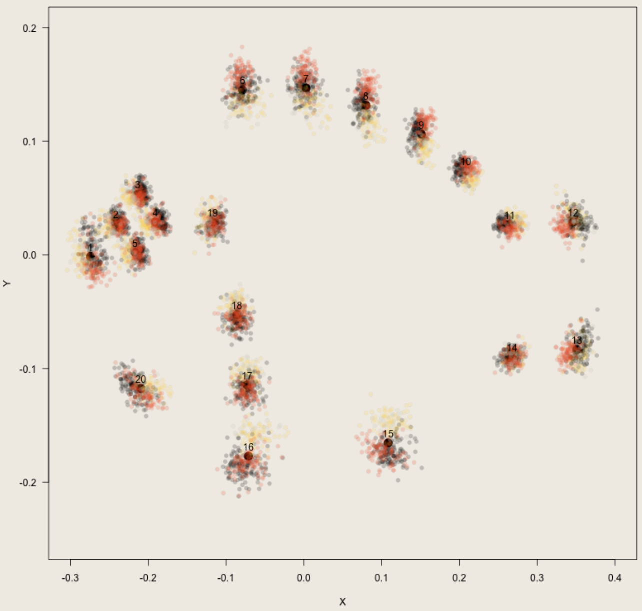


**Fig. S15**: Position of landmarks after Procrustes superimposition for each phenotyped sample. Numbered black dots represent the mean position of landmarks, Transparent point represent landmark position by sample. Colors represent the reproductive host association associated with the sample species (Red: Entacmaea specialist, Yellow: Radianthus specialist, Grey: Stichodactyla specialist, Black: Generalist)

| Dataset | BM | WN | OU | OUM |
|---|---|---|---|---|
| Color patterns (white) | 49.45 | 10.45 | 14.71 | **0** |
| Color patterns (orange) | 34.32 | **0** | 12.40 | 4.62 |
| Color patterns (black) | 39.44 | 20.03 | **0** | 4.17 |
| Color patterns (rgb) | **0.70** | 16.42 | 22.27 | 0 |
| Morphospace | 21.79 | 37.58 | 46.17 | **0** |
| Morphology | 58.17 | 89.55 | 43.32 | **0** |

**Tab. S7**: $\Delta AICc$ comparison for multivariate evolutionary models on phenotypes. We bold the selected model based on $\Delta Aicc$ criterion

seem to be driven by high evolutionary rates in *Entacmaea* specialists while species that have a different type of association have low evolutionary rates. Both results seem concordant with multivariate analyses. However univariate models fitted on orange patterns seem discordant with multivariate models (Tab. S8). Both PC1 and PC2 of orange patterns best fit an OUM model. PC3 and PC4 have likely driven the absence of signal in multivariate analyses as they are treated equally but represent a small portion of orange color patterns variation.

##### RGB color patterns

Using multivariate models, the dataset including the three RGB channels best fitted an OUM model (Tab. S7, multiple evolutionary optima driven by reproductive host associations). However, we did not have enough power to reject the BM model. Using the BM as a selected model, phylogenetic MANOVA detected an influence of reproductive host association on RGB color patterns (Pillai: $S = 0.8181, p = 0.003$). A discrepancy between those two approaches could be due their different design. The multivariate models are assuming that each category has a different optima while, the phylogenetic MANOVA is testing whether one of the categories is different from the others. If a reproductive host category is not evolving towards a different optima, adding a parameter for this category will not improve the likelihood but will reduce AICc values.

Univariate models confirm the observed trend (Tab. S11). The two first PC axes of RGB color patterns best fit a OUM model (different optima driven by reproductive host association). We conclude that the presence or absence of a head bar and a second bar, as well as the orange intensity of off-stripe colors are likely driven by reproductive host associations. However PC3 and PC4 best fit density-dependent models. PC3, which measures the width of both the head stripe and the second stripe and the brightness of the caudal fin, best fits a Matching-Competition where phenotypes of co-occurring species diverges from the mean community value. PC4, that measures dorsal patterns and the width of the head bar, best fits a density-dependent model where evolutionary rates are decreasing when more species co-occur (Tab. S11).

##### Morphology

Both morphology datasets (traits and PCA) fit best a multivariate OUM model (Tab. S7, several optima driven

| Model | $\Delta AICc$ | $\theta_0$ | $\sigma^2$ | $\sigma^2_S$ | $\sigma^2_E$ | $\sigma^2_R$ | $\sigma^2_G$ | $\alpha$ | $\theta_S$ | $\theta_E$ | $\theta_R$ | $\theta_G$ | $b_{lin}$ | $r_{exp}$ | $S$ |
|---|---|---|---|---|---|---|---|---|---|---|---|---|---|---|---|
| PC1 White | | | | | | | | | | | | | | | |
| BM | 5.9 | -0.24 | 0.19 | | | | | | | | | | | | |
| BMM | 1.1 | -0.89 | | 0.47 | 0.0012 | 0.58 | 0.11 | | | | | | | | |
| WN | 2.9 | -0.57 | 0.73 | | | | | | | | | | | | |
| OU | 3.6 | -0.53 | 0.73 | | | | | 0.5 | | | | | | | |
| **OUM** | **0** | | **31** | | | | | **33** | **-1.6** | **-1** | **-0.005** | **-0.42** | | | |
| $MC$ | 7.6 | -0.25 | 0.18 | | | | | | | | | | | | -1.2e-08 |
| $DD_{lin}$ | 4.9 | -0.57 | 1.5e-08 | | | | | | | | | | 9.4e-10 | | |
| $DD_{exp}$ | 4.9 | -0.64 | 0.0018 | | | | | | | | | | | 0.51 | |
| PC2 White | | | | | | | | | | | | | | | |
| BM | 180 | -0.91 | 7.9e-57 | | | | | | | | | | | | |
| BMM | 25 | -0.33 | | 1.6 | 0.54 | 0.24 | 0.36 | | | | | | | | |
| WN | 180 | -0.91 | 7.9e-57 | | | | | | | | | | | | |
| OU | 4.8 | -0.58 | 5.1 | | | | | 2.5 | | | | | | | |
| **OUM** | **0** | | **36** | | | | | **32** | **-1.9** | **0.34** | **-0.52** | **-0.77** | | | |
| $MC$ | 4.8 | -0.59 | 7.9e-10 | | | | | | | | | | | | -0.00043 |
| $DD_{lin}$ | 4.8 | -0.59 | 2e-04 | | | | | | | | | | -1.5e-05 | | |
| $DD_{exp}$ | 4.8 | -0.59 | 4.6e-08 | | | | | | | | | | | -1 | |

**Tab. S8**: $\Delta AICc$ comparison and parameter estimates for univariate evolutionary models on white color patterns. We bold the selected model based on $\Delta Aicc$ criterion

| Model | $\Delta AICc$ | $\theta_0$ | $\sigma^2$ | $\sigma^2_S$ | $\sigma^2_E$ | $\sigma^2_R$ | $\sigma^2_G$ | $\alpha$ | $\theta_S$ | $\theta_E$ | $\theta_R$ | $\theta_G$ | $b_{lin}$ | $r_{exp}$ | $S$ |
|---|---|---|---|---|---|---|---|---|---|---|---|---|---|---|---|
| PC1 Orange | | | | | | | | | | | | | | | |
| BM | 14 | -0.2 | 0.33 | | | | | | | | | | | | |
| BMM | 18 | -0.56 | | 1e-09 | 0.69 | 0.43 | 0.32 | | | | | | | | |
| WN | 320 | -1.7 | 7.9e-10 | | | | | | | | | | | | |
| OU | 2.8 | -0.61 | 2 | | | | | 1.3 | | | | | | | |
| **OUM** | **0** | | **8.4** | | | | | **7.8** | **-1.7** | **-0.48** | **-0.012** | **-0.81** | | | |
| $MC$ | 16 | -0.21 | 0.31 | | | | | | | | | | | | -2.7e-08 |
| $DD_{lin}$ | 15 | -0.16 | 0.49 | | | | | | | | | | -0.035 | | |
| $DD_{exp}$ | 0.78 | -0.14 | 24 | | | | | | | | | | | -2.9 | |
| PC2 Orange | | | | | | | | | | | | | | | |
| BM | 6.9 | -0.0013 | 0.1 | | | | | | | | | | | | |
| BMM | 12 | 0.84 | | 0.24 | 1e-09 | 0.072 | 0.17 | | | | | | | | |
| WN | 110 | 0.23 | 7.9e-10 | | | | | | | | | | | | |
| OU | 6.2 | -0.028 | 2.8 | | | | | 2.7 | | | | | | | |
| **OUM** | **0** | | **14** | | | | | **22** | **-1.1** | **0.8** | **0.1** | **-0.32** | | | |
| $MC$ | 6.1 | -0.029 | 2.1e-30 | | | | | | | | | | | | -2.6 |
| $DD_{lin}$ | 6.4 | -0.022 | 0.0032 | | | | | | | | | | -0.00016 | | |
| $DD_{exp}$ | 2 | 0.11 | 8.4e+8 | | | | | | | | | | | -90 | |

**Tab. S9**: $\Delta AICc$ comparison and parameter estimates for univariate evolutionary models on orange color patterns. We bold the selected model based on $\Delta Aicc$ criterion

| Model | $\Delta AICc$ | $\theta_0$ | $\sigma^2$ | $\sigma^2_S$ | $\sigma^2_E$ | $\sigma^2_R$ | $\sigma^2_G$ | $\alpha$ | $\theta_S$ | $\theta_E$ | $\theta_R$ | $\theta_G$ | $b_{lin}$ | $r_{exp}$ | $S$ |
|---|---|---|---|---|---|---|---|---|---|---|---|---|---|---|---|
| PC1 Black | | | | | | | | | | | | | | | |
| BM | 89 | -1.4 | 2.4e-8 | | | | | | | | | | | | |
| BMM | 15 | -0.3 | | 1e-09 | 0.69 | 0.22 | 0.35 | | | | | | | | |
| WN | 88 | -1.4 | 2.2e-05 | | | | | | | | | | | | |
| **OU** | **0.77** | **-0.57** | **2.8** | | | | | **2.1** | | | | | | | |
| OUM | 0 | | 15 | | | | | 15 | -1.4 | -0.2 | -0.15 | -0.8 | | | |
| $MC$ | 18 | -0.19 | 0.33 | | | | | | | | | | | | -1e-08 |
| $DD_{lin}$ | 16 | -0.16 | 0.49 | | | | | | | | | | -0.035 | | |
| $DD_{exp}$ | 2.8 | -0.58 | 4.3e+10 | | | | | | | | | | | -24 | |
| PC2 Black | | | | | | | | | | | | | | | |
| BM | 18 | -0.13 | 2.4e-8 | | | | | | | | | | | | |
| **BMM** | **0** | **-0.13** | | **1e-09** | **0.43** | **1e-09** | **1e-09** | | | | | | | | |
| WN | 18 | -0.13 | 2.4e-8 | | | | | | | | | | | | |
| OU | 20 | -0.13 | 1.7e-21 | | | | | 4.7e-182 | | | | | | | |
| OUM | 26 | | 1e-09 | | | | | 2.7e-07 | -22000 | -27000 | -0.053 | -27000 | | | |
| $MC$ | 33 | -0.34 | 0.0022 | | | | | | | | | | | | -0.37 |
| $DD_{lin}$ | 36 | -0.38 | 0.029 | | | | | | | | | | -0.0021 | | |
| $DD_{exp}$ | 32 | -0.21 | 1.8 | | | | | | | | | | | -1.8 | |

**Tab. S10**: $\Delta AICc$ comparison and parameter estimates for univariate evolutionary models on black color patterns. We bold the selected model based on $\Delta Aicc$ criterion

| Model | $\Delta AICc$ | $\theta_0$ | $\sigma^2$ | $\sigma_S^2$ | $\sigma_E^2$ | $\sigma_R^2$ | $\sigma_G^2$ | $\alpha$ | $\theta_S$ | $\theta_E$ | $\theta_R$ | $\theta_G$ | $b_{lin}$ | $r_{exp}$ | $S$ |
|---|---|---|---|---|---|---|---|---|---|---|---|---|---|---|---|
| PC1 | | | | | | | | | | | | | | | |
| BM | 8.8 | -0.2 | 0.078 | | | | | | | | | | | | |
| BMM | 11 | 0.14 | | 1e-09 | 0.26 | 1e-09 | 0.052 | | | | | | | | |
| WN | 92 | 0.49 | 7.9e-7 | | | | | | | | | | | | |
| OU | 11 | -0.18 | 0.1 | | | | | 0.057 | | | | | | | |
| **OUM** | **0** | | **0.18** | | | | | **0.8** | **0.32** | **-1.3** | **0.5** | **0.41** | | | |
| $MC$ | 11 | -0.15 | 0.069 | | | | | | | | | | | | -0.038 |
| $DD_{lin}$ | 10 | -0.19 | 5.4e-15 | | | | | | | | | | 0.015 | | |
| $DD_{exp}$ | 10 | -0.17 | 0.0097 | | | | | | | | | | | 0.29 | |
| PC2 | | | | | | | | | | | | | | | |
| BM | 3.4 | 0.15 | 0.15 | | | | | | | | | | | | |
| BMM | 3.5 | -0.15 | | 0.32 | 1e-09 | 0.051 | 0.25 | | | | | | | | |
| WN | 220 | -0.3 | 7.9e-16 | | | | | | | | | | | | |
| OU | 5.4 | 0.16 | 0.18 | | | | | 0.033 | | | | | | | |
| **OUM** | **0** | | **0.2** | | | | | **0.21** | **2.7** | **-0.034** | **-1.4** | **1** | | | |
| $MC$ | 5 | 0.42 | 0.063 | | | | | | | | | | | | -0.2 |
| $DD_{lin}$ | 5.4 | 0.13 | 0.21 | | | | | | | | | | -0.011 | | |
| $DD_{exp}$ | 5.4 | 0.14 | 0.21 | | | | | | | | | | | -0.07 | |
| PC3 | | | | | | | | | | | | | | | |
| BM | 44 | 0.016 | 7.9e-16 | | | | | | | | | | | | |
| BMM | 9.2 | 1.6 | | 1e-09 | 0.01 | 1e-09 | 0.11 | | | | | | | | |
| WN | 44 | 0.016 | 7.9e-16 | | | | | | | | | | | | |
| OU | 5.9 | 0.73 | 0.046 | | | | | 7.1e-16 | | | | | | | |
| OUM | 10 | | 0.037 | | | | | 3.8e-08 | 7.8 | -2.2 | 1.4 | -2.5 | | | |
| **MC** | **0** | **0.28** | **0.00064** | | | | | | | | | | | | **-0.43** |
| $DD_{lin}$ | 5.1 | 0.77 | 0.068 | | | | | | | | | | -0.0049 | | |
| $DD_{exp}$ | 4.5 | 0.78 | 0.17 | | | | | | | | | | | -0.46 | |
| PC4 | | | | | | | | | | | | | | | |
| BM | 10 | 0.14 | 0.14 | | | | | | | | | | | | |
| BMM | 10 | 0.21 | | 1e-09 | 0.074 | 0.48 | 0.041 | | | | | | | | |
| WN | 270 | 1.2 | 7.9e-7 | | | | | | | | | | | | |
| OU | 6.5 | 0.12 | 0.57 | | | | | 0.55 | | | | | | | |
| OUM | 9.8 | | 3.5 | | | | | 4 | 0.36 | 0.49 | 0.36 | -0.23 | | | |
| $MC$ | 7.3 | 0.14 | 3.7e-12 | | | | | | | | | | | | -1.2 |
| $DD_{lin}$ | 7.9 | 0.14 | 0.046 | | | | | | | | | | -0.0033 | | |
| **$DD_{exp}$** | **0** | **0.0035** | **4.1e+14** | | | | | | | | | | | **-34** | |

**Tab. S11**: $\Delta AICc$ comparison and parameter estimates for univariate evolutionary models on rgb color patterns. We bold the selected model based on $\Delta Aicc$ criterion

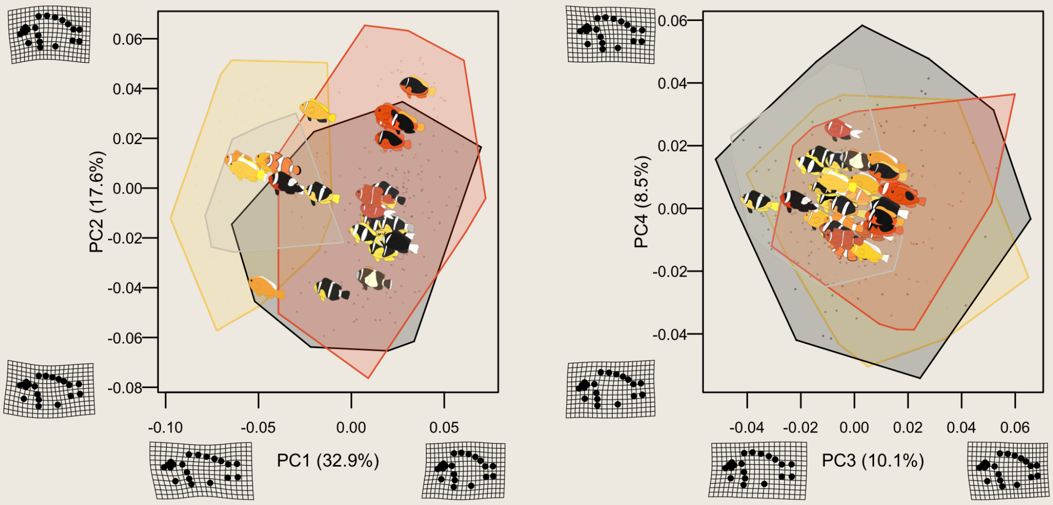


**Fig. S16**: Position of each clownfish species in the morphospace. Morphospace was generated by calculating a PCA on landmarks coordinates corrected by the Procrustes analysis. The grid deformation plot indicate the direction of morphological variation represented by each PCA axis. Fish images represent the mean position of each species in the morphospace. Hull polygons represent the part of the morphospace occupied by each reproductive host association categories (Red: Entacmaea specialist, Yellow: Radianthus specialist, Grey: Stichodactyla specialist, Black: Generalist)

by host associations). Phylogenetic MANOVAs give concordant results for the trait dataset (Pillai: $S = 1.2989, p = 0.001$) and the PCA dataset (Pillai: $S = 1.5279, p = 0.001$).

Trait data best fit an univariate OUM model for body ratio, head ratio, peduncle ratio and snout angle. We did not find phylogenetic signal for eye-height ratio (Tab. S12). The PCA dataset, which reflects better the total morphological variation between individuals gives contrasted results. PC1 and PC3 best fit a OUM model, while PC2 fits better a DDexp model (density dependent model) indicating that this morphology axis is evolving faster where more species co-occur (Tab. S13). Finally, we did not find phylogenetic signal for PC4 (Tab. S13).

#### Robustness to ASR uncertainties

For univariate analyses, we find that some results are not robust to uncertainties relative to ancestral state reconstruction. For instance, RGB color patterns PC2 best fits a BM with 40% of stochastic maps (Fig. S23) while it better fits an OUM model with the marginal reproductive host association reconstruction. Black patterns PC1 and PC2 only fit better a OUM model with 2% and 11% of stochastic maps respectively (Fig. S24). Conversely, eye height ratio fits better an OUM with 60% of stochastic maps as opposed to the marginal reconstruction that best fits a WN model. These uncertainties can also affect parameter estimation. For PC4 of RGB color patterns, there is a high uncertainty on the estimation of r which ranges from $-2$ to 2. Although stochastic maps lead to the same selected model, its interpretation can range from an accelerated evolution where more species co-occur to a decelerated evolution in the same areas (Fig. S33).

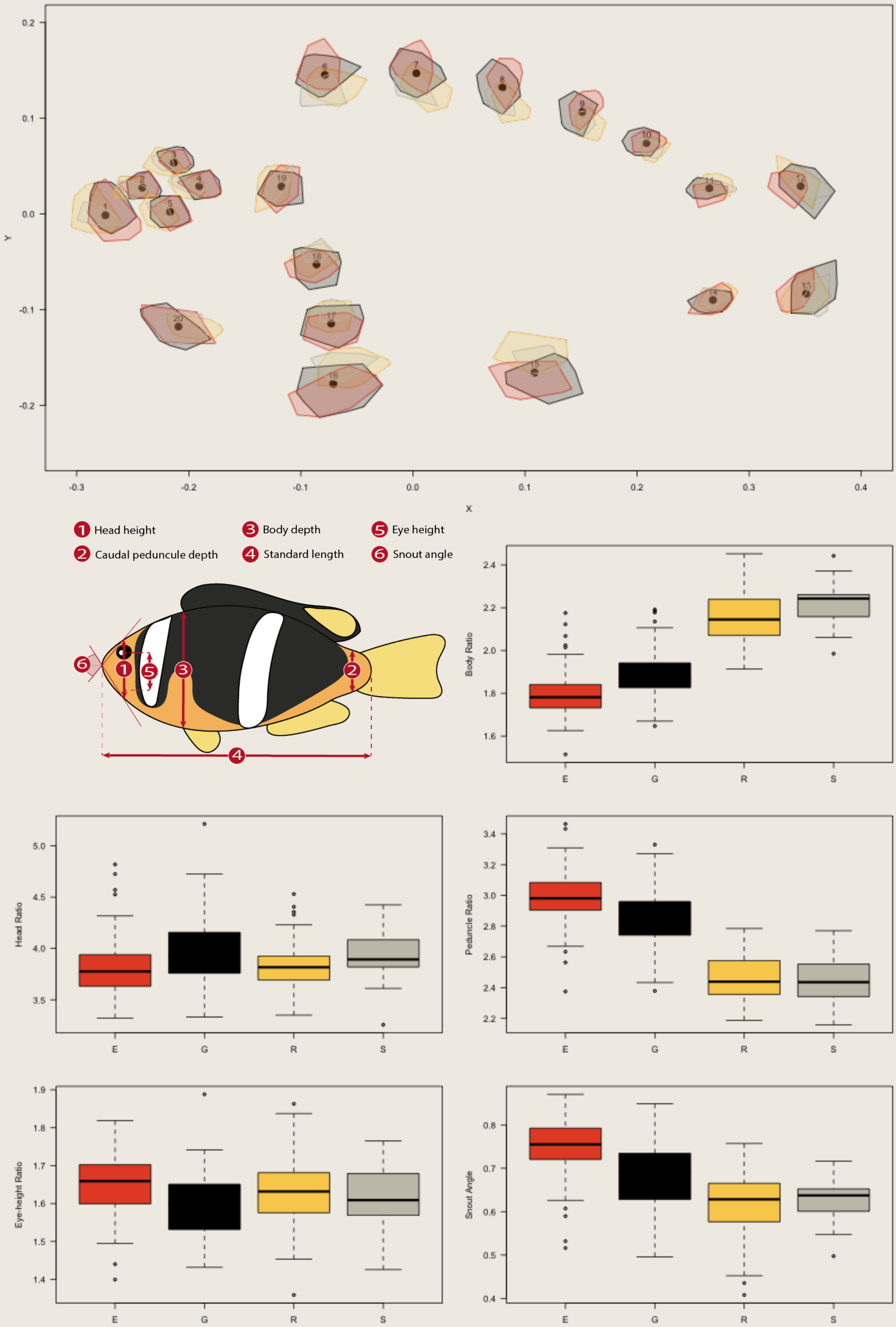


**Fig. S17**: Effect of reproductive host association categorization on morphology. Top plot represent the sample variation of each landmark position by reproductive host association. Boxplots represent the distribution of five traits of interest by reproductive host association (Red: Entacmaea specialist, Yellow: Radianthus specialist, Grey: Stichodactyla specialist, Black: Generalist). The five traits are derived from measures obtained from the landmarks shown on the top left schematic view (Body Ratio: 3/4, Head Ratio: 3/1, Peduncle Ratio: 3/2, Eye-height Ratio: 1/5)

| Model | $\Delta AICc$ | $\theta_0$ | $\sigma^2$ | $\sigma^2_S$ | $\sigma^2_E$ | $\sigma^2_R$ | $\sigma^2_G$ | $\alpha$ | $\theta_S$ | $\theta_E$ | $\theta_R$ | $\theta_G$ | $b_{lin}$ | $r_{exp}$ | $S$ |
|---|---|---|---|---|---|---|---|---|---|---|---|---|---|---|---|
| Body ratio | | | | | | | | | | | | | | | |
| BM | 40 | 2 | 0.0023 | | | | | | | | | | | | |
| BMM | 29 | 1.9 | | 0.015 | 0.0088 | 0.00052 | 1e-09 | | | | | | | | |
| WN | 54 | 1.9 | 0.018 | | | | | | | | | | | | |
| OU | 42 | 2 | 0.0023 | | | | | 1.4e-16 | | | | | | | |
| **OUM** | **0** | | **1e-09** | | | | | **0.62** | **2.2** | **2.1** | **1.8** | **1.9** | | | |
| $MC$ | 39 | 2 | 0.00063 | | | | | | | | | | | | -0.3 |
| $DD_{lin}$ | 40 | 2 | 1.5e-11 | | | | | | | | | | 0.00042 | | |
| $DD_{exp}$ | 41 | 2 | 0.00047 | | | | | | | | | | | 0.23 | |
| Head ratio | | | | | | | | | | | | | | | |
| BM | 30 | 3.6 | 2.4e-8 | | | | | | | | | | | | |
| BMM | 13 | 3.9 | | 1e-09 | 1e-09 | 0.0072 | 1e-09 | | | | | | | | |
| WN | 30 | 3.6 | 2.4e-8 | | | | | | | | | | | | |
| OU | 10 | 3.9 | 0.0065 | | | | | 0.19 | | | | | | | |
| **OUM** | **0** | | **1e-09** | | | | | **0.31** | **4** | **3.8** | **3.5** | **4** | | | |
| $MC$ | 11 | 3.8 | 0.00088 | | | | | | | | | | | | -0.13 |
| $DD_{lin}$ | 10 | 3.9 | 3.2e-09 | | | | | | | | | | 0.00049 | | |
| $DD_{exp}$ | 7.3 | 3.9 | 1.3e-16 | | | | | | | | | | | 2.7 | |
| Peduncle ratio | | | | | | | | | | | | | | | |
| BM | 34 | 2.7 | 0.006 | | | | | | | | | | | | |
| BMM | 27 | 2.9 | | 0.029 | 0.014 | 0.0039 | 1e-09 | | | | | | | | |
| WN | 46 | 2.8 | 0.038 | | | | | | | | | | | | |
| OU | 36 | 2.7 | 0.006 | | | | | 4.4e-15 | | | | | | | |
| **OUM** | **0** | | **1e-09** | | | | | **0.25** | **2.1** | **2.4** | **3.1** | **2.9** | | | |
| $MC$ | 35 | 2.7 | 0.003 | | | | | | | | | | | | -0.19 |
| $DD_{lin}$ | 34 | 2.7 | 1.2e-11 | | | | | | | | | | 0.0011 | | |
| $DD_{exp}$ | 35 | 2.7 | 0.0012 | | | | | | | | | | | 0.23 | |

This table continues next page

| Model | $\Delta AICc$ | $\theta_0$ | $\sigma^2$ | $\sigma^2_S$ | $\sigma^2_E$ | $\sigma^2_R$ | $\sigma^2_G$ | $\alpha$ | $\theta_S$ | $\theta_E$ | $\theta_R$ | $\theta_G$ | $b_{lin}$ | $r_{exp}$ | $S$ |
|---|---|---|---|---|---|---|---|---|---|---|---|---|---|---|---|
| Eye-Height ratio | | | | | | | | | | | | | | | |
| BM | 2 | 1.6 | 5e-04 | | | | | | | | | | | | |
| BMM | 2.1 | 1.6 | | 0.0015 | 0.00099 | 0.00072 | 1e-09 | | | | | | | | |
| **WN** | **0.39** | **1.6** | **0.0024** | | | | | | | | | | | | |
| OU | 1.7 | 1.6 | 0.0017 | | | | | 0.35 | | | | | | | |
| OUM | 0.85 | | 1e-09 | | | | | 0.093 | 1.9 | 1.4 | 1.8 | 1.4 | | | |
| $MC$ | 2.4 | 1.6 | 2.7e-12 | | | | | | | | | | | | -0.11 |
| $DD_{lin}$ | 1.9 | 1.6 | 1.1e-10 | | | | | | | | | | 4.8e-05 | | |
| $DD_{exp}$ | 0 | 1.6 | 5.4e-81 | | | | | | | | | | | 14 | |
| Snout Angle | | | | | | | | | | | | | | | |
| BM | 7.7 | 0.66 | 0.00048 | | | | | | | | | | | | |
| BMM | 3.9 | 0.69 | | 1e-09 | 0.0015 | 0.00055 | 1e-09 | | | | | | | | |
| WN | 13 | 0.68 | 0.0029 | | | | | | | | | | | | |
| OU | 9.5 | 0.67 | 0.00066 | | | | | 0.063 | | | | | | | |
| **OUM** | **0** | | **0.00038** | | | | | **0.38** | **0.65** | **0.58** | **0.78** | **0.69** | | | |
| $MC$ | 8.5 | 0.67 | 0.00016 | | | | | | | | | | | | -0.24 |
| $DD_{lin}$ | 8.3 | 0.66 | 1.9e-11 | | | | | | | | | | 8.5e-05 | | |
| $DD_{exp}$ | 6.2 | 0.65 | 4.3e-16 | | | | | | | | | | | 2.5 | |

**Tab. S12**: $\Delta AICc$ comparison and parameter estimates for univariate evolutionary models on morphological traits. We bold the selected model based on $\Delta Aicc$ criterion.

| Model | $\Delta AICc$ | $\theta_0$ | $\sigma^2$ | $\sigma^2_S$ | $\sigma^2_E$ | $\sigma^2_R$ | $\sigma^2_G$ | $\alpha$ | $\theta_S$ | $\theta_E$ | $\theta_R$ | $\theta_G$ | $b_{lin}$ | $r_{exp}$ | $S$ |
|---|---|---|---|---|---|---|---|---|---|---|---|---|---|---|---|
| PC1 | | | | | | | | | | | | | | | |
| BM | 37 | -0.0087 | 1e-04 | | | | | | | | | | | | |
| BMM | 29 | 0.017 | | 0.00029 | 5e-04 | 3.4e-05 | 1e-09 | | | | | | | | |
| WN | 51 | -0.00087 | 0.00075 | | | | | | | | | | | | |
| OU | 39 | -0.0087 | 1e-04 | | | | | 1.1e-15 | | | | | | | |
| **OUM** | **0** | | **1e-09** | | | | | **0.59** | **-0.03** | **-0.051** | **0.038** | **0.017** | | | |
| $MC$ | 37 | -0.0031 | 5e-05 | | | | | | | | | | | | -0.19 |
| $DD_{lin}$ | 38 | -0.0093 | 3.4e-11 | | | | | | | | | | 1.9e-05 | | |
| $DD_{exp}$ | 38 | -0.0092 | 4.5e-05 | | | | | | | | | | | 0.13 | |
| PC2 | | | | | | | | | | | | | | | |
| BM | 7.1 | -0.0029 | 4.6e-05 | | | | | | | | | | | | |
| BMM | 6.4 | -0.012 | | 1e-09 | 0.00033 | 0.00012 | 1e-09 | | | | | | | | |
| WN | 7.5 | -0.0018 | 0.00027 | | | | | | | | | | | | |
| OU | 7.6 | -0.0022 | 0.00014 | | | | | 0.23 | | | | | | | |
| OUM | 3.7 | | 0.00017 | | | | | 0.75 | 0.0032 | 2.4e-06 | 0.021 | -0.018 | | | |
| $MC$ | 7.9 | -0.0013 | 1e-05 | | | | | | | | | | | | -0.15 |
| $DD_{lin}$ | 7.3 | -0.0024 | 6.8e-12 | | | | | | | | | | 5e-06 | | |
| **DD$_{\mathbf{exp}}$** | **0** | **-0.0092** | **6.1e-95** | | | | | | | | | | | **16** | |
| PC3 | | | | | | | | | | | | | | | |
| BM | 8.3 | -0.00024 | 4.6e-06 | | | | | | | | | | | | |
| BMM | 7.4 | 0.003 | | 0.00022 | 1e-09 | 3.1e-06 | 1e-09 | | | | | | | | |
| WN | 9.3 | 0.0011 | 1.6e-05 | | | | | | | | | | | | |
| OU | 10 | -0.00014 | 9e-06 | | | | | 0.1 | | | | | | | |
| **OUM** | **0** | | **1e-09** | | | | | **0.82** | **-0.034** | **0.0046** | **0.0092** | **-0.0046** | | | |
| $MC$ | 5 | 0.00076 | 7.2e-15 | | | | | | | | | | | | -1.3 |
| $DD_{lin}$ | 9.6 | -0.00064 | 1e-12 | | | | | | | | | | 1.1e-06 | | |
| $DD_{exp}$ | 9.7 | -0.00096 | 3.7e-07 | | | | | | | | | | | 0.36 | |
| PC4 | | | | | | | | | | | | | | | |
| BM | 0 | 0.0056 | 2.4e-16 | | | | | | | | | | | | |
| BMM | 6 | 0.0056 | | 1e-09 | 1e-09 | 1.4e-08 | 1e-09 | | | | | | | | |
| **WN** | **0** | **0.0056** | **2.4e-16** | | | | | | | | | | | | |
| OU | 2 | 0.0056 | 1.1e-21 | | | | | 1.4e-09 | | | | | | | |
| OUM | 5.2 | | 1e-09 | | | | | 4.3e-08 | -9.1 | -1.5 | 0.00048 | 0.21 | | | |
| $MC$ | 2 | 0.0056 | 1.5e-16 | | | | | | | | | | | | -0.39 |
| $DD_{lin}$ | 2 | 0.0056 | 4.8e-13 | | | | | | | | | | 4.9e-13 | | |
| $DD_{exp}$ | 2 | 0.0056 | 5e-13 | | | | | | | | | | | -23 | |

**Tab. S13**: $\Delta AICc$ comparison and parameter estimates for univariate evolutionary models on morphospace. We bold the selected model based on $\Delta Aicc$ criterion.

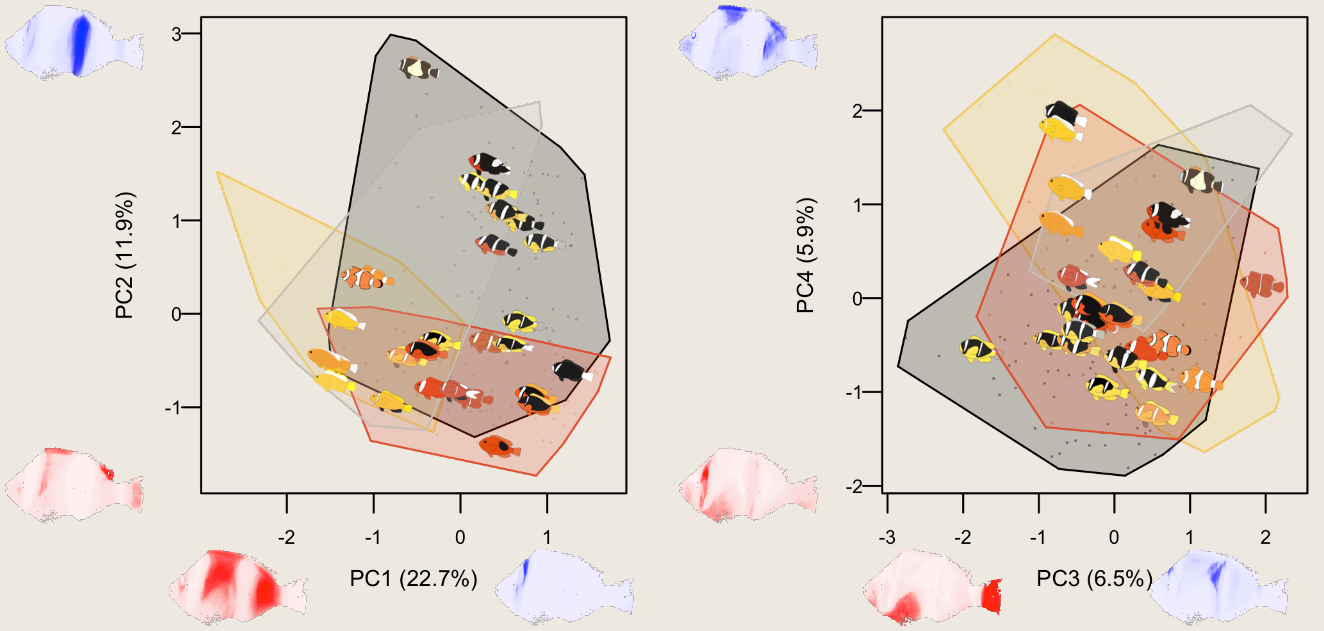


**Fig. S18**: Position of each clownfish species in the RGB color pattern space. RGB color pattern space was generated by calculating a PCA on a concatenation of Red Green and Blue channels of standardised clownfish images. The fish images along axes represent the individual contribution of each pixel to the corresponding axis on a standardised clownfish shape. Intensity of red represent the magnitude of negative contribution of each pixel to the corresponding axis. Intensity of blue represent the magnitude of positive contribution of each pixel to the corresponding axis. Fish images represent the mean position of each species in the RGB color pattern space. Hull polygons represent the part of the RGB color pattern space occupied by each reproductive host association categories (Red: Entacmaea specialist, Yellow: Radianthus specialist, Grey: Stichodactyla specialist, Black: Generalist)

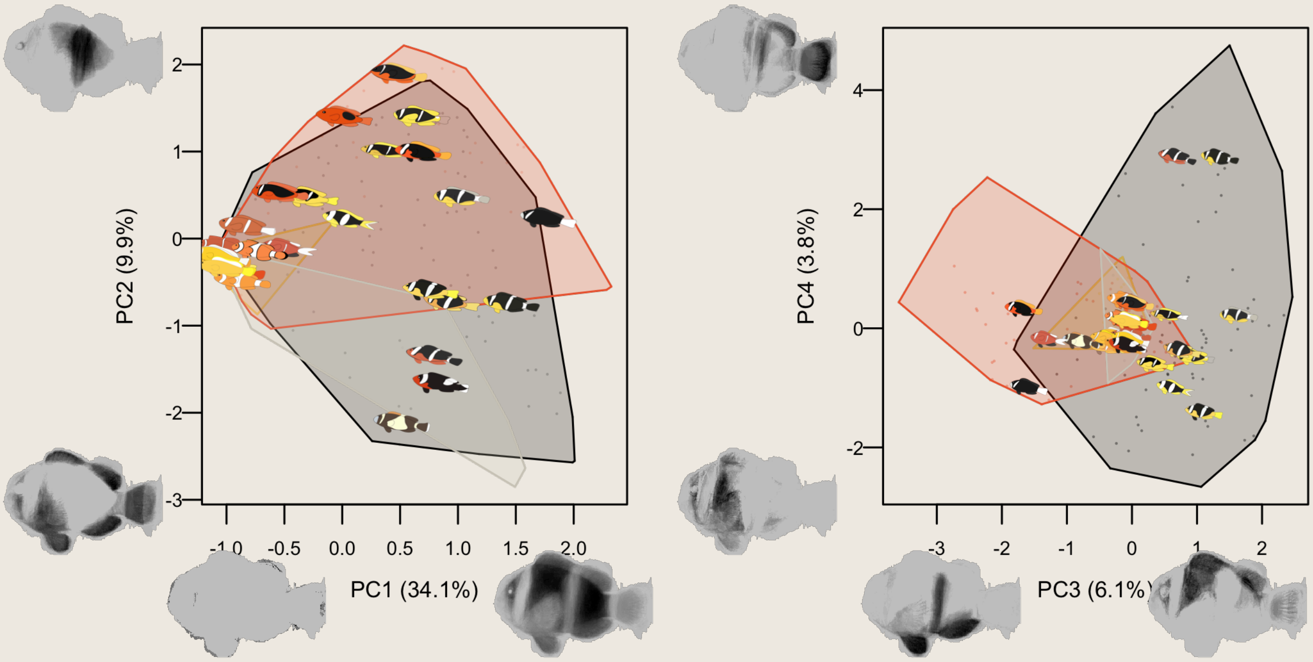


**Fig. S19**: Position of each clownfish species in the black pattern space. black pattern space was generated by calculating a PCA on the black channel (RGB color code: 0,0,0) of standardised clownfish images. The fish images along axes represent the individual contribution of each pixel to the corresponding axis on a standardised clownfish shape. Intensity of black represent the magnitude of the contribution of each pixel to the corresponding axis. Fish images represent the mean position of each species in the black pattern space. Hull polygons represent the part of the black color pattern space occupied by each reproductive host association categories (Red: Entacmaea specialist, Yellow: Radianthus specialist, Grey: Stichodactyla specialist, Black: Generalist)

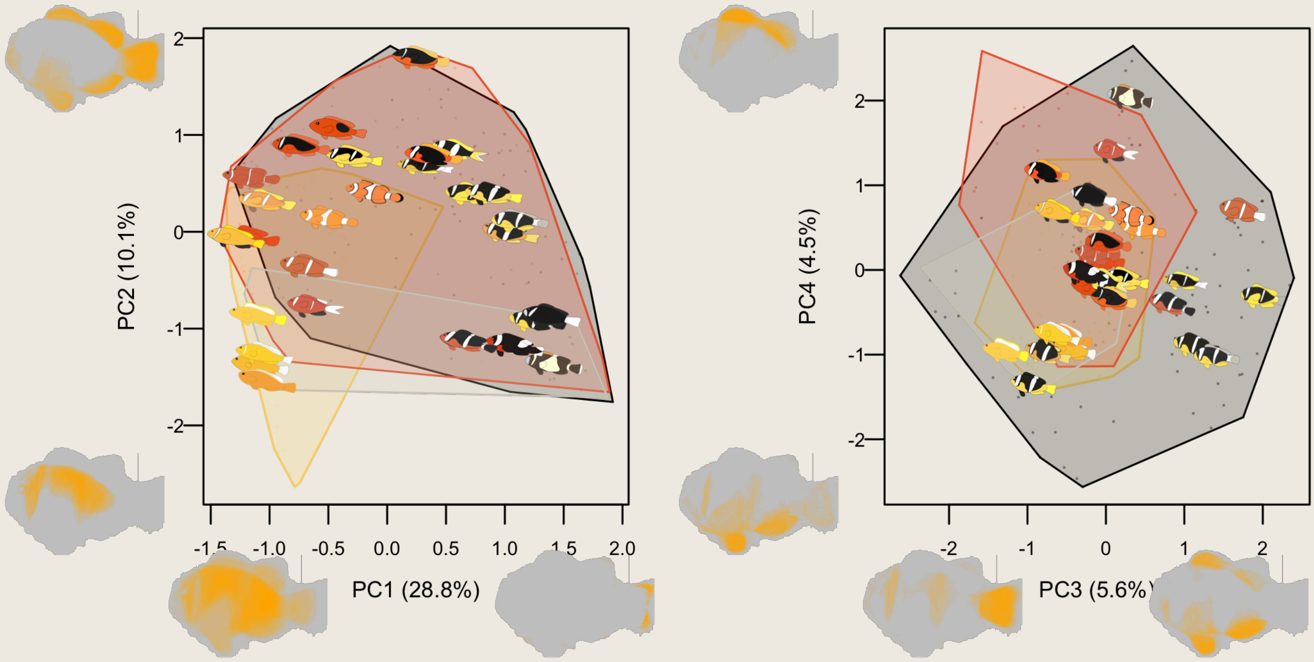


**Fig. S20**: Position of each clownfish species in the orange pattern space. orange pattern space was generated by calculating a PCA on the orange channel (RGB color code: 255,165,0) of standardised clownfish images. The fish images along axes represent the individual contribution of each pixel to the corresponding axis on a standardised clownfish shape. Intensity of orange represent the magnitude of the contribution of each pixel to the corresponding axis. Fish images represent the mean position of each species in the orange pattern space. Hull polygons represent the part of the orange color pattern space occupied by each reproductive host association categories (Red: Entacmaea specialist, Yellow: Radianthus specialist, Grey: Stichodactyla specialist, orange: Generalist)

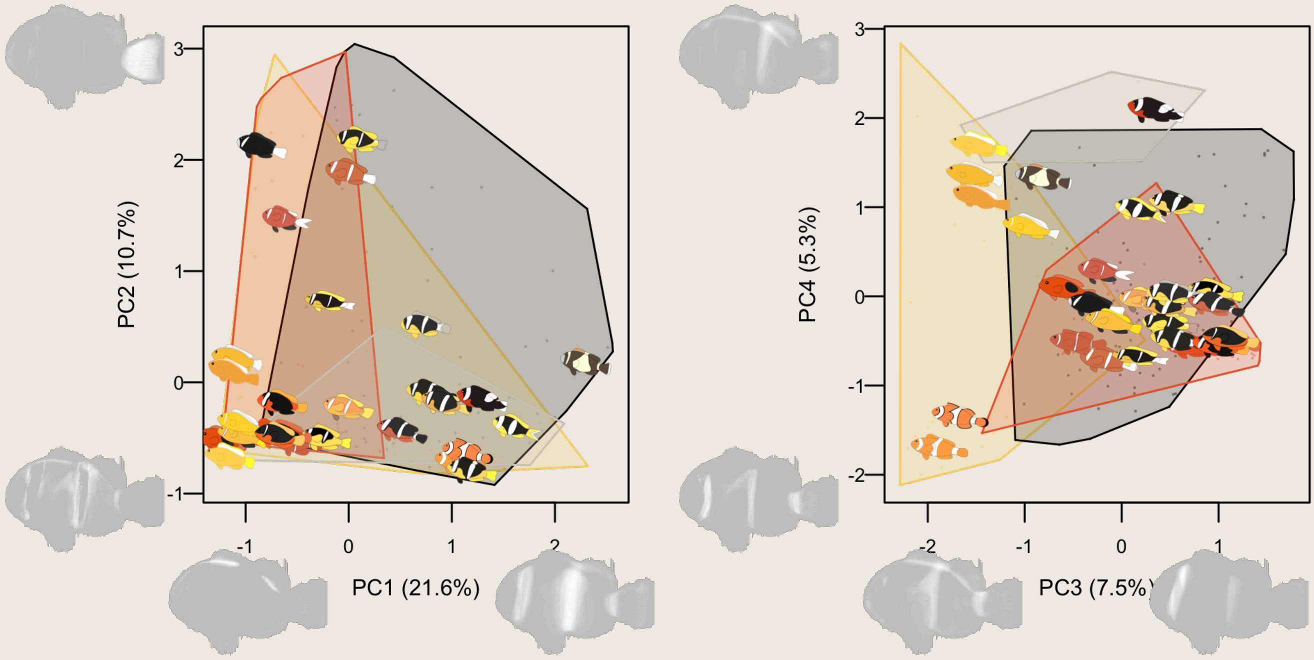


**Fig. S21**: Position of each clownfish species in the white pattern space. white pattern space was generated by calculating a PCA on the white channel (RGB color code: 255,255,255) of standardised clownfish images. The fish images along axes represent the individual contribution of each pixel to the corresponding axis on a standardised clownfish shape. Intensity of white represent the magnitude of the contribution of each pixel to the corresponding axis. Fish images represent the mean position of each species in the white pattern space. Hull polygons represent the part of the white color pattern space occupied by each reproductive host association categories (Red: Entacmaea specialist, Yellow: Radianthus specialist, Grey: Stichodactyla specialist, white: Generalist)

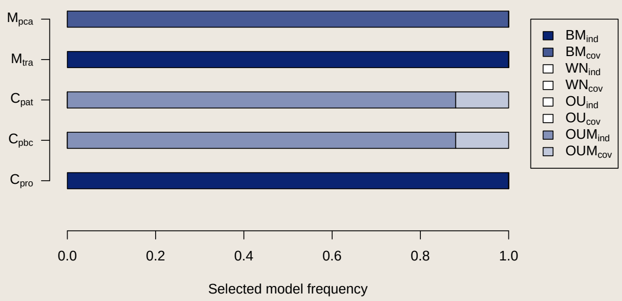


**Fig. S22**: Best multivariate PCM models for host-use stochastic maps. For each trait dataset, we ran multivariate PCM models on 100 stochastic maps of host-use ancestral state reconstructions for host-dependent models (OUM, BMM) or 100 stochastic maps of biogeography history for density-dependent models (MC, DDl, DDe). We compared alternative models using AICc. With a $\Delta AICc < 2$, we retained the NULL model. Each histogram represents the frequency at which models have been selected. $M_{pca}$: Morphospace, $M_{tra}$: Morphology traits, $C_{pat}$: coloration patterns, $C_{pbc}$: color-specific patterns, $C_{pro}$: color proportions.

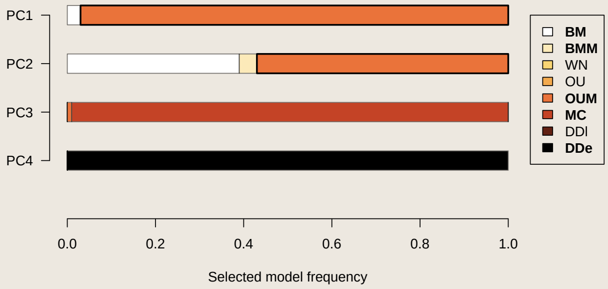


**Fig. S23**: Best univariate PCM models for host-use stochastic maps. For color patterns dataset, we ran univariate PCM models on 100 stochastic maps of host-use ancestral state reconstructions for host-dependent models (OUM, BMM) or 100 stochastic maps of biogeography history for density-dependent models (MC, DDl, DDe). We compared alternative models using AICc. With a $\Delta AICc < 2$, we retained the NULL model. Each histogram represents the frequency at which models have been selected.

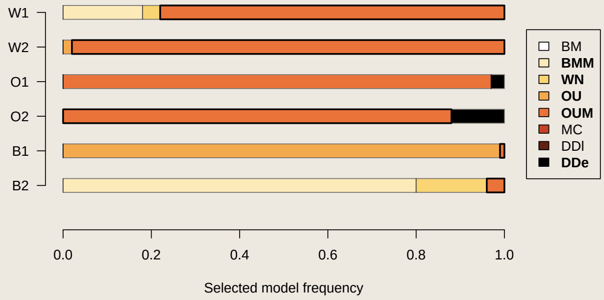


**Fig. S24**: Best univariate PCM models for host-use stochastic maps. For color specific patterns dataset, we ran univariate PCM models on 100 stochastic maps of host-use ancestral state reconstructions for host-dependent models (OUM, BMM) or 100 stochastic maps of biogeography history for density-dependent models (MC, DDl, DDe). We compared alternative models using AICc. With a $\Delta AICc < 2$, we retained the NULL model. Each histogram represents the frequency at which models have been selected. $O1$: PC1 of orange patterns, $O2$: PC2 of orange patterns, $W1$: PC1 of white patterns, $W2$: PC2 of white patterns, $B1$: PC1 of black patterns, $B2$: PC2 of black patterns

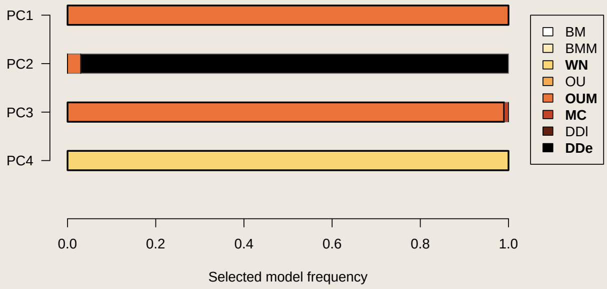


**Fig. S25**: Best univariate PCM models for host-use stochastic maps. For each morphospace axis, we ran univariate PCM models on 100 stochastic maps of host-use ancestral state reconstructions for host-dependent models (OUM, BMM) or 100 stochastic maps of biogeography history for density-dependent models (MC, DDl, DDe). We compared alternative models using AICc. With a $\Delta AICc < 2$, we retained the NULL model. Each histogram represents the frequency at which models have been selected.

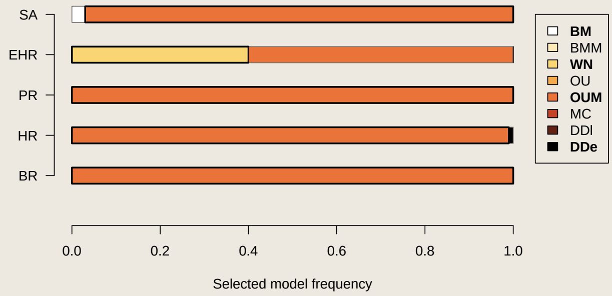


**Fig. S26**: Best univariate PCM models for host-use stochastic maps. For each morphological trait, we ran univariate PCM models on 100 stochastic maps of host-use ancestral state reconstructions for host-dependent models (OUM, BMM) or 100 stochastic maps of biogeography history for density-dependent models (MC, DDl, DDe). We compared alternative models using AICc. With a $\Delta AICc < 2$, we retained the NULL model. Each histogram represents the frequency at which models have been selected. BR: Body ratio, PR: Peduncule ratio, HR: Height ratio EHR: Eye-height ratio, SA: Snout angle

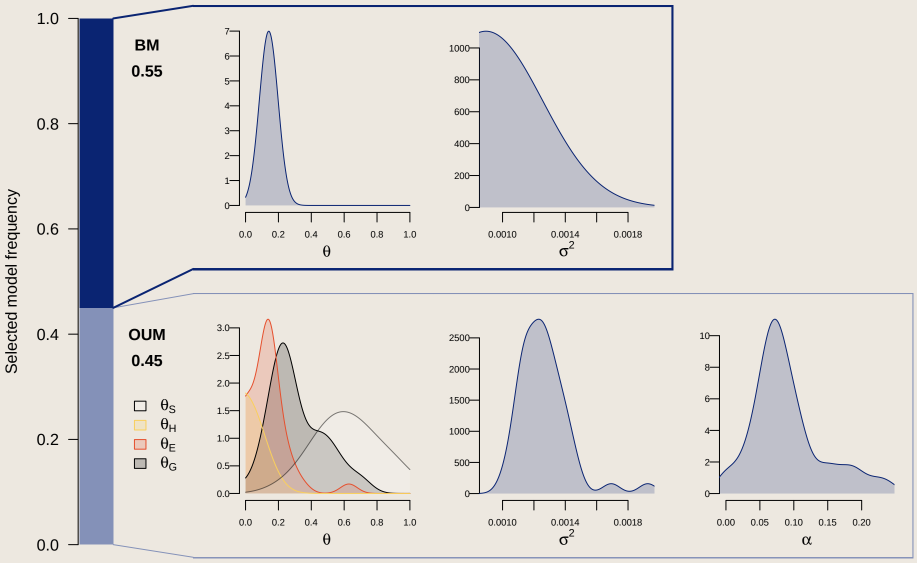


**Fig. S27**: Evolutionary rates for the proportion of white estimated from host-use stochastic maps. We show the distribution of parameters from univariate PCM models ran on 100 stochastic maps of host-use ancestral state reconstructions for host-dependent models (OUM, BMM) or 100 stochastic maps of biogeography history for density-dependent models (MC, DDl, DDe). We show parameter estimation for models selected from more than 25% of stochastic maps. The highlighted frame represents the selected model from the marginal ancestral state reconstructions.

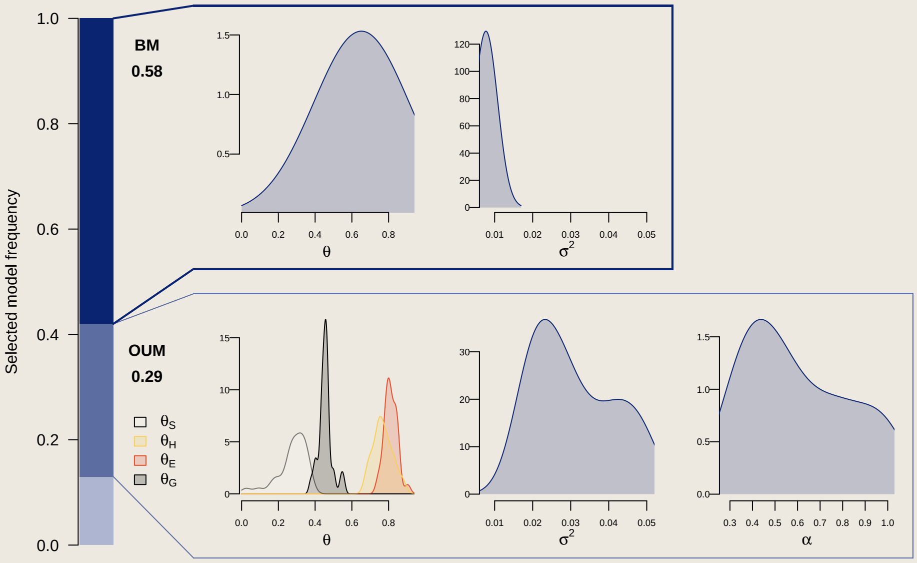


**Fig. S28**: Evolutionary rates for the proportion of orange estimated from host-use stochastic maps. We show the distribution of parameters from univariate PCM models ran on 100 stochastic maps of host-use ancestral state reconstructions for host-dependent models (OUM, BMM) or 100 stochastic maps of biogeography history for density-dependent models (MC, DDl, DDe). We show parameter estimation for models selected from more than 25% of stochastic maps. The highlighted frame represents the selected model from the marginal ancestral state reconstructions.

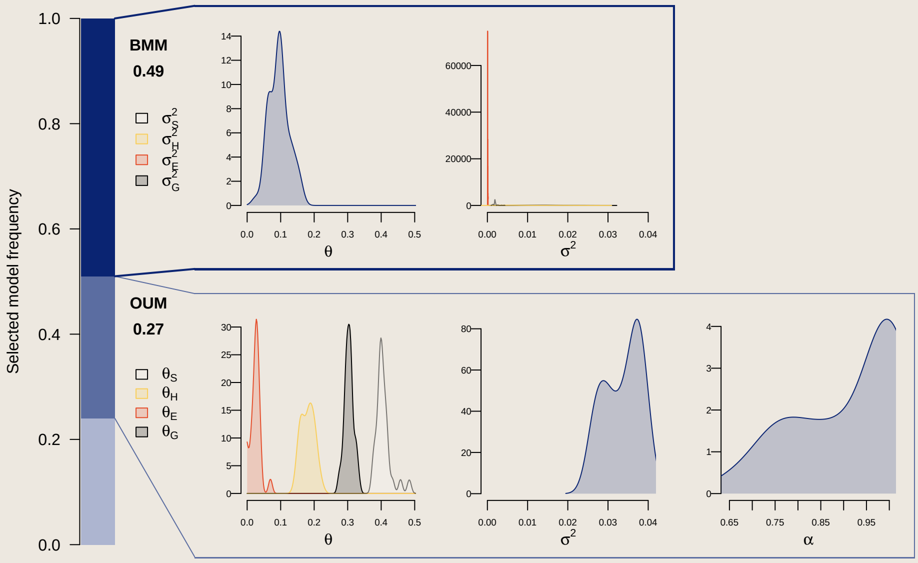


**Fig. S29**: Evolutionary rates for the proportion of black estimated from host-use stochastic maps. We show the distribution of parameters from univariate PCM models ran on 100 stochastic maps of host-use ancestral state reconstructions for host-dependent models (OUM, BMM) or 100 stochastic maps of biogeography history for density-dependent models (MC, DDl, DDe). We show parameter estimation for models selected from more than 25% of stochastic maps. The highlighted frame represents the selected model from the marginal ancestral state reconstructions.

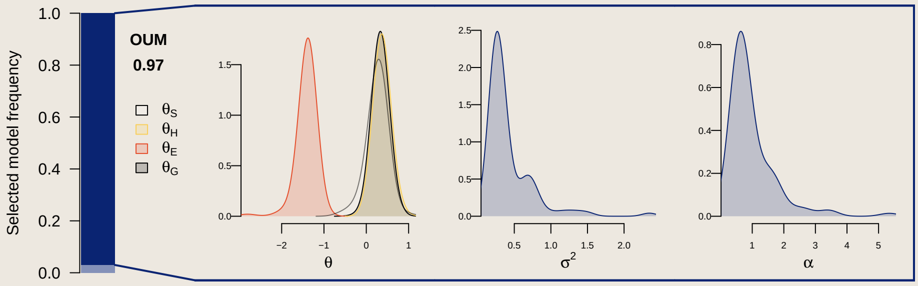


**Fig. S30**: Evolutionary rates for the first axis of color patterns estimated from host-use stochastic maps. We show the distribution of parameters from univariate PCM models ran on 100 stochastic maps of host-use ancestral state reconstructions for host-dependent models (OUM, BMM) or 100 stochastic maps of biogeography history for density-dependent models (MC, DDl, DDe). We show parameter estimation for models selected from more than 25% of stochastic maps. The highlighted frame represents the selected model from the marginal ancestral state reconstructions.

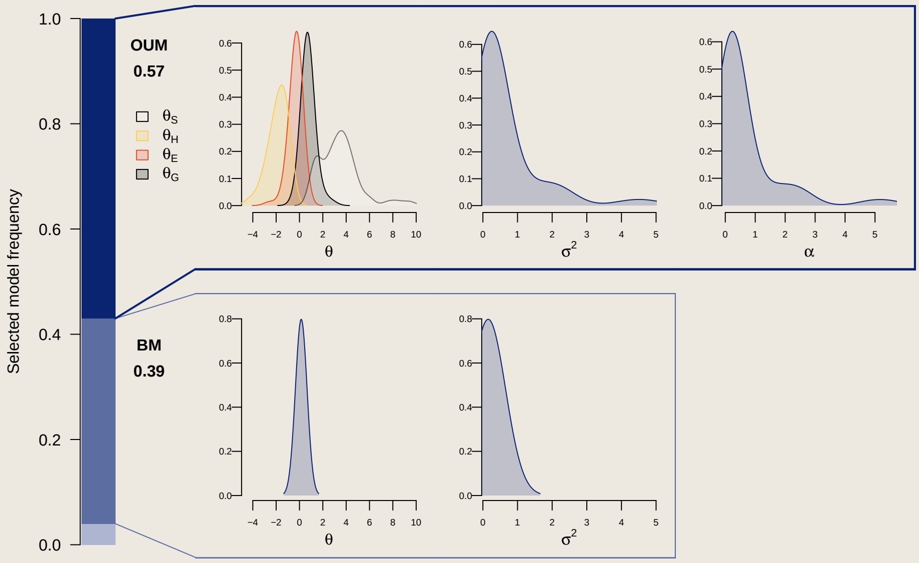


**Fig. S31**: Evolutionary rates for the second axis of color patterns estimated from host-use stochastic maps. We show the distribution of parameters from univariate PCM models ran on 100 stochastic maps of host-use ancestral state reconstructions for host-dependent models (OUM, BMM) or 100 stochastic maps of biogeography history for density-dependent models (MC, DDl, DDe). We show parameter estimation for models selected from more than 25% of stochastic maps. The highlighted frame represents the selected model from the marginal ancestral state reconstructions.

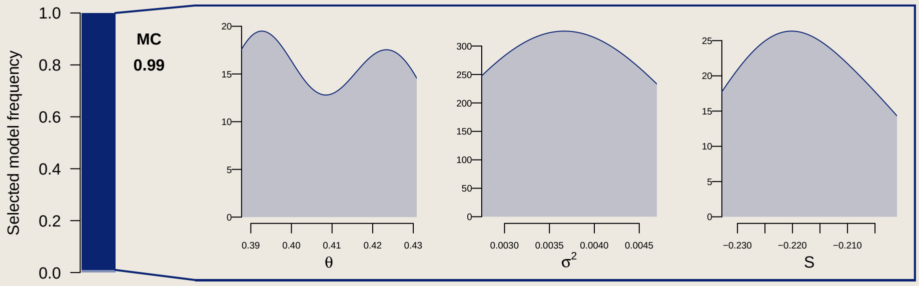


**Fig. S32**: Evolutionary rates for the third axis of color patterns estimated from host-use stochastic maps. We show the distribution of parameters from univariate PCM models ran on 100 stochastic maps of host-use ancestral state reconstructions for host-dependent models (OUM, BMM) or 100 stochastic maps of biogeography history for density-dependent models (MC, DDl, DDe). We show parameter estimation for models selected from more than 25% of stochastic maps. The highlighted frame represents the selected model from the marginal ancestral state reconstructions.

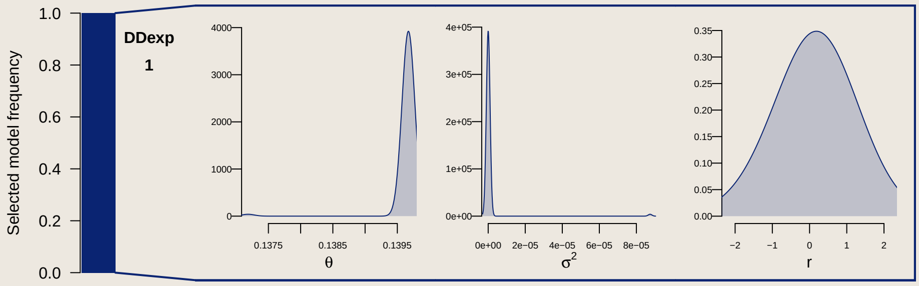


**Fig. S33**: Evolutionary rates for the fourth axis of color patterns estimated from host-use stochastic maps. We show the distribution of parameters from univariate PCM models ran on 100 stochastic maps of host-use ancestral state reconstructions for host-dependent models (OUM, BMM) or 100 stochastic maps of biogeography history for density-dependent models (MC, DDl, DDe). We show parameter estimation for models selected from more than 25% of stochastic maps. The highlighted frame represents the selected model from the marginal ancestral state reconstructions.

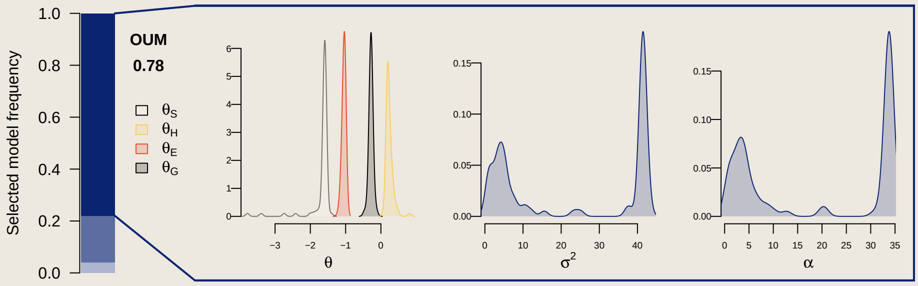


**Fig. S34**: Evolutionary rates for the first axis of white patterns estimated from host-use stochastic maps. We show the distribution of parameters from univariate PCM models ran on 100 stochastic maps of host-use ancestral state reconstructions for host-dependent models (OUM, BMM) or 100 stochastic maps of biogeography history for density-dependent models (MC, DDl, DDe). We show parameter estimation for models selected from more than 25% of stochastic maps. The highlighted frame represents the selected model from the marginal ancestral state reconstructions.

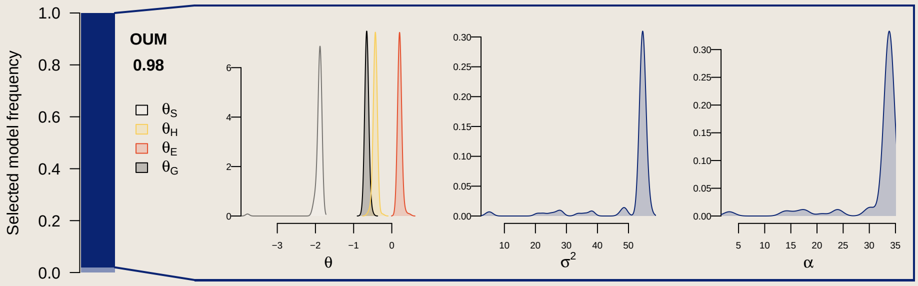


**Fig. S35**: Evolutionary rates for the second axis of white patterns estimated from host-use stochastic maps. We show the distribution of parameters from univariate PCM models ran on 100 stochastic maps of host-use ancestral state reconstructions for host-dependent models (OUM, BMM) or 100 stochastic maps of biogeography history for density-dependent models (MC, DDl, DDe). We show parameter estimation for models selected from more than 25% of stochastic maps. The highlighted frame represents the selected model from the marginal ancestral state reconstructions.

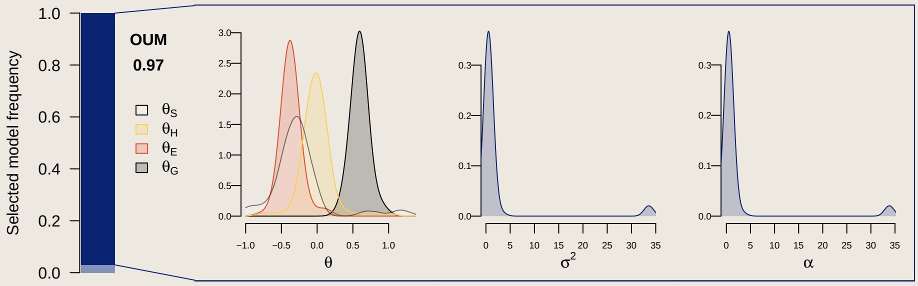


**Fig. S36**: Evolutionary rates for the first axis of orange patterns estimated from host-use stochastic maps. We show the distribution of parameters from univariate PCM models ran on 100 stochastic maps of host-use ancestral state reconstructions for host-dependent models (OUM, BMM) or 100 stochastic maps of biogeography history for density-dependent models (MC, DDl, DDe). We show parameter estimation for models selected from more than 25% of stochastic maps. The highlighted frame represents the selected model from the marginal ancestral state reconstructions.

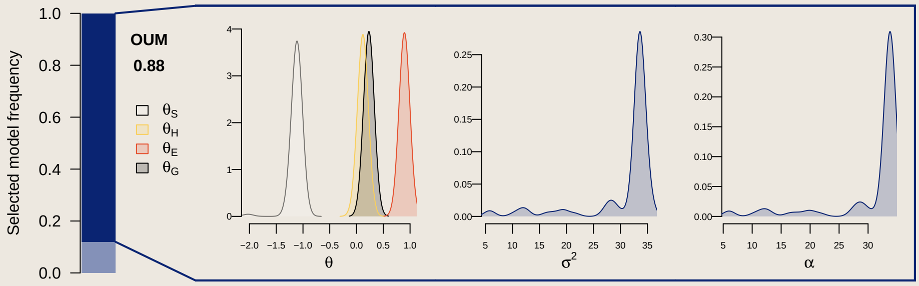


**Fig. S37**: Evolutionary rates for the second axis of orange patterns estimated from host-use stochastic maps. We show the distribution of parameters from univariate PCM models ran on 100 stochastic maps of host-use ancestral state reconstructions for host-dependent models (OUM, BMM) or 100 stochastic maps of biogeography history for density-dependent models (MC, DDl, DDe). We show parameter estimation for models selected from more than 25% of stochastic maps. The highlighted frame represents the selected model from the marginal ancestral state reconstructions.

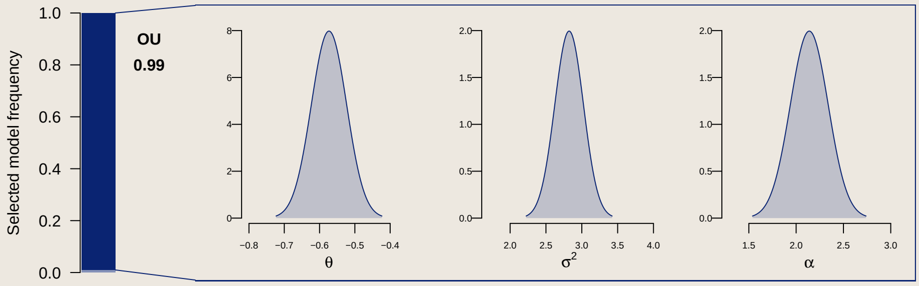


**Fig. S38**: Evolutionary rates for the first axis of black patterns estimated from host-use stochastic maps. We show the distribution of parameters from univariate PCM models ran on 100 stochastic maps of host-use ancestral state reconstructions for host-dependent models (OUM, BMM) or 100 stochastic maps of biogeography history for density-dependent models (MC, DDl, DDe). We show parameter estimation for models selected from more than 25% of stochastic maps. The highlighted frame represents the selected model from the marginal ancestral state reconstructions.

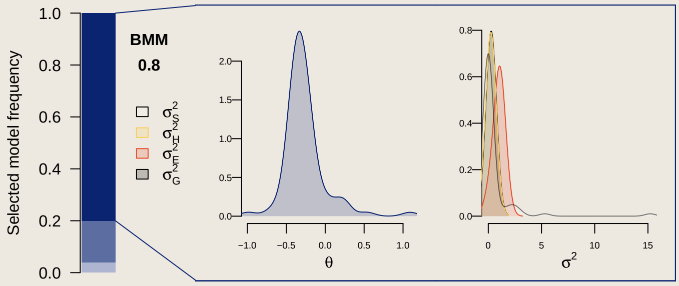


**Fig. S39**: Evolutionary rates for the second axis of black patterns estimated from host-use stochastic maps. We show the distribution of parameters from univariate PCM models ran on 100 stochastic maps of host-use ancestral state reconstructions for host-dependent models (OUM, BMM) or 100 stochastic maps of biogeography history for density-dependent models (MC, DDl, DDe). We show parameter estimation for models selected from more than 25% of stochastic maps. The highlighted frame represents the selected model from the marginal ancestral state reconstructions.

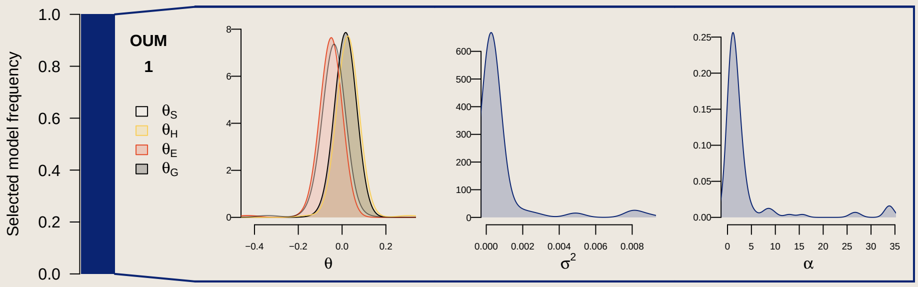


**Fig. S40**: Evolutionary rates for the first axis of morphospace estimated from host-use stochastic maps. We show the distribution of parameters from univariate PCM models ran on 100 stochastic maps of host-use ancestral state reconstructions for host-dependent models (OUM, BMM) or 100 stochastic maps of biogeography history for density-dependent models (MC, DDl, DDe). We show parameter estimation for models selected from more than 25% of stochastic maps. The highlighted frame represents the selected model from the marginal ancestral state reconstructions.

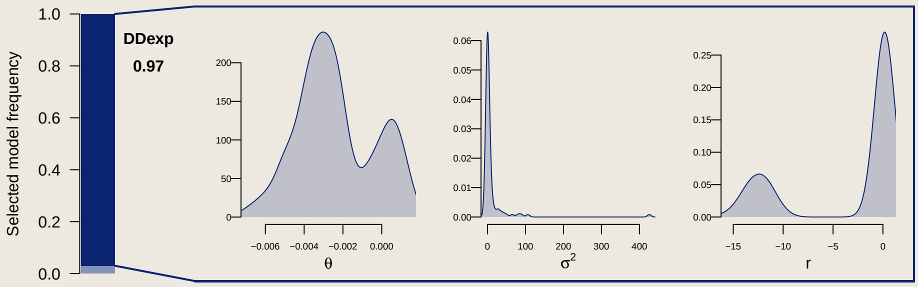


**Fig. S41**: Evolutionary rates for the second axis of morphospace estimated from host-use stochastic maps. We show the distribution of parameters from univariate PCM models ran on 100 stochastic maps of host-use ancestral state reconstructions for host-dependent models (OUM, BMM) or 100 stochastic maps of biogeography history for density-dependent models (MC, DDl, DDe). We show parameter estimation for models selected from more than 25% of stochastic maps. The highlighted frame represents the selected model from the marginal ancestral state reconstructions.

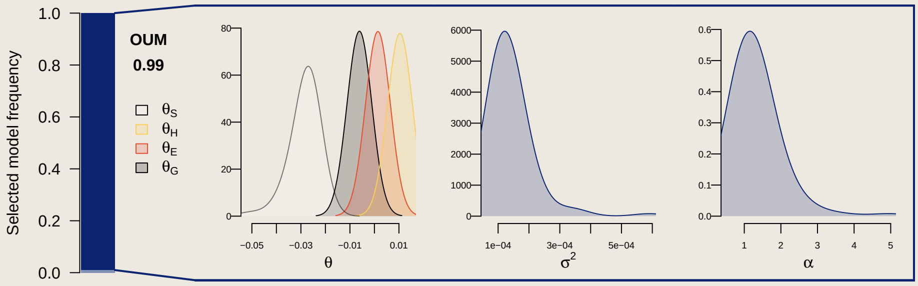


**Fig. S42**: Evolutionary rates for the third axis of morphospace estimated from host-use stochastic maps. We show the distribution of parameters from univariate PCM models ran on 100 stochastic maps of host-use ancestral state reconstructions for host-dependent models (OUM, BMM) or 100 stochastic maps of biogeography history for density-dependent models (MC, DDl, DDe). We show parameter estimation for models selected from more than 25% of stochastic maps. The highlighted frame represents the selected model from the marginal ancestral state reconstructions.

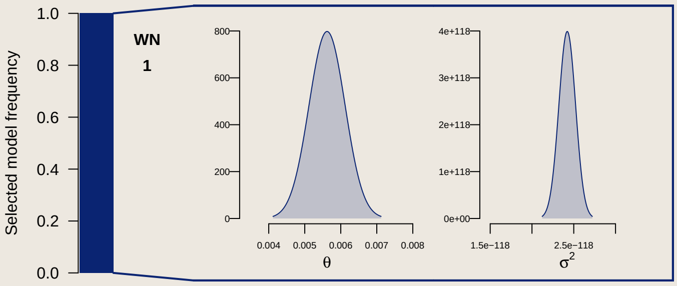


**Fig. S43**: Evolutionary rates for the fourth axis of morphospace estimated from host-use stochastic maps. We show the distribution of parameters from univariate PCM models ran on 100 stochastic maps of host-use ancestral state reconstructions for host-dependent models (OUM, BMM) or 100 stochastic maps of biogeography history for density-dependent models (MC, DDl, DDe). We show parameter estimation for models selected from more than 25% of stochastic maps. The highlighted frame represents the selected model from the marginal ancestral state reconstructions.

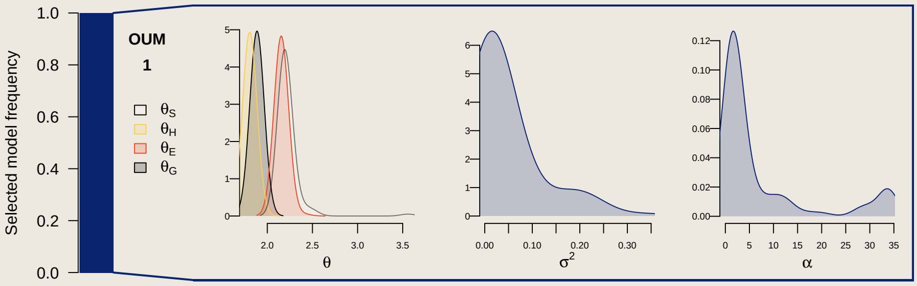


**Fig. S44**: Evolutionary rates for body ratio estimated from host-use stochastic maps. We show the distribution of parameters from univariate PCM models ran on 100 stochastic maps of host-use ancestral state reconstructions for host-dependent models (OUM, BMM) or 100 stochastic maps of biogeography history for density-dependent models (MC, DDl, DDe). We show parameter estimation for models selected from more than 25% of stochastic maps. The highlighted frame represents the selected model from the marginal ancestral state reconstructions.

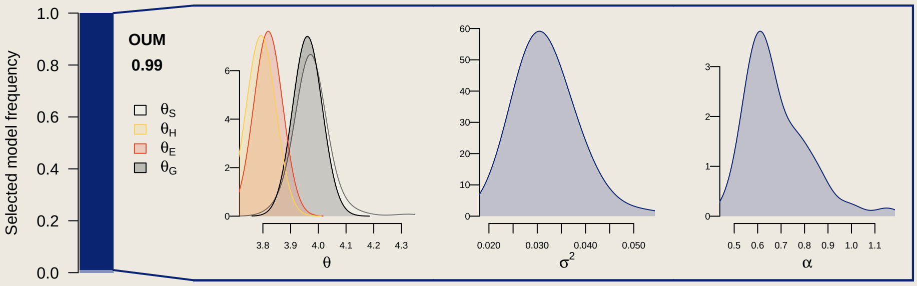


**Fig. S45**: Evolutionary rates for height ratio estimated from host-use stochastic maps. We show the distribution of parameters from univariate PCM models ran on 100 stochastic maps of host-use ancestral state reconstructions for host-dependent models (OUM, BMM) or 100 stochastic maps of biogeography history for density-dependent models (MC, DDl, DDe). We show parameter estimation for models selected from more than 25% of stochastic maps. The highlighted frame represents the selected model from the marginal ancestral state reconstructions.

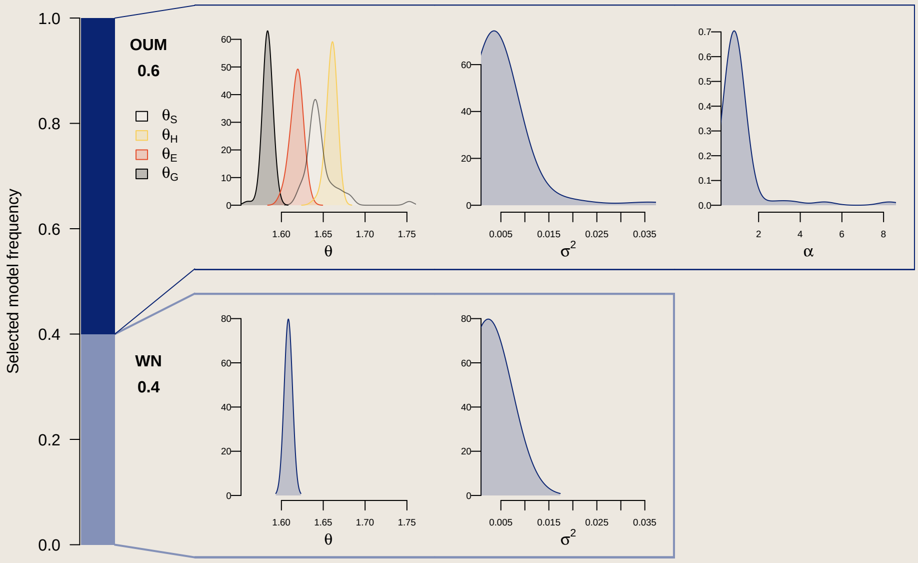


**Fig. S46**: Evolutionary rates for eye height ratio estimated from host-use stochastic maps. We show the distribution of parameters from univariate PCM models ran on 100 stochastic maps of host-use ancestral state reconstructions for host-dependent models (OUM, BMM) or 100 stochastic maps of biogeography history for density-dependent models (MC, DDl, DDe). We show parameter estimation for models selected from more than 25% of stochastic maps. The highlighted frame represents the selected model from the marginal ancestral state reconstructions.

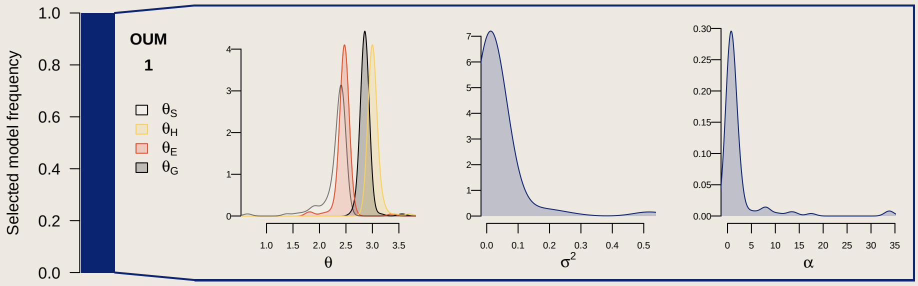


**Fig. S47**: Evolutionary rates for peduncule ratio estimated from host-use stochastic maps. We show the distribution of parameters from univariate PCM models ran on 100 stochastic maps of host-use ancestral state reconstructions for host-dependent models (OUM, BMM) or 100 stochastic maps of biogeography history for density-dependent models (MC, DDl, DDe). We show parameter estimation for models selected from more than 25% of stochastic maps. The highlighted frame represents the selected model from the marginal ancestral state reconstructions.

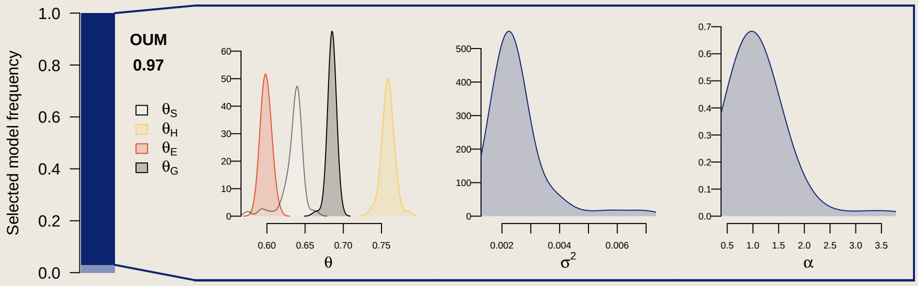


**Fig. S48**: Evolutionary rates for snout angle estimated from host-use stochastic maps. We show the distribution of parameters from univariate PCM models ran on 100 stochastic maps of host-use ancestral state reconstructions for host-dependent models (OUM, BMM) or 100 stochastic maps of biogeography history for density-dependent models (MC, DDl, DDe). We show parameter estimation for models selected from more than 25% of stochastic maps. The highlighted frame represents the selected model from the marginal ancestral state reconstructions.
